# NovAST: a deep learning framework for automated label transfer and novel cell type discovery in spatial transcriptomics

**DOI:** 10.64898/2026.11.18.767106

**Authors:** Yu Zhu, Ritambhara Singh, Ying Ma

## Abstract

Spatially resolved transcriptomics enables the mapping of gene expression within tissue architecture, transforming our understanding of cellular organization and function. However, its interpretation critically depends on accurate cell type annotation. As recent technologies now achieve single-cell resolution, annotation has become essential for linking molecular profiles to tissue architecture and disease mechanisms. However, reference-based annotation methods rely on the assumption that all cell types in the target tissue are present in the reference, which is frequently violated in complex or pathological contexts, causing systematic misclassification and masking novel or context-specific cell populations. Here we introduce **NovAST** (**Nov**el cell type discovery and **A**nnotation in **S**patial **T**ranscriptomics), a fully automated deep learning framework that jointly performs reference-guided annotation and novel cell type discovery. NovAST learns a cluster-aware latent space that preserves cell-type structure, supports graph-based label propagation, and explicitly quantifies assignment uncertainty. This model design enables NovAST to automatically infer whether novel cell populations are absent, singular, or multiple, a question that cannot be specified a priori, thereby avoiding the incorrect assumption of user-defined novel cell types required by existing approaches. Applied to eleven spatial transcriptomics reference-target pairs spanning MERFISH, CosMx SMI, and 10x Xenium platforms, NovAST consistently recovers known cell types while uncovering previously unannotated, disease-associated novel cell types, including immune-associated cell states in idiopathic pulmonary fibrosis and inflammatory cell populations in human colon tissue.

## 1 Introduction

Recent advances in high-resolution spatially resolved transcriptomics (SRT) technologies now enable gene expression profiling at single-cell or subcellular resolution, providing unprecedented opportunities to chart cellular heterogeneity and tissue architecture *in situ*. Accurate cell type annotation is essential for interpreting these data [1, 2]. Without reliable annotation, expression signals from mixed or closely related populations become confounded, limiting the interpretation of transcriptional variation within its biological and spatial context. Conversely, accurate annotation enables SRT data to reveal how molecular programs are organized across tissues and how specific cell populations and microenvironments contribute to disease processes. As a result, cell type annotation underpins a wide range of downstream analyses, including cell type-specific differential expression (DE) and spatial variable genes (SVGs) detection [3, 4, 5, 6], cell–cell communication inference [7, 8], and developmental trajectory analysis [9].

Two major strategies are currently used for cell type annotation in SRT. The first is unsupervised clustering-based annotation, where cells are embedded in a low-dimensional space (e.g., via principal component analysis, PCA) and grouped based on local similarity captured by a *`k`*-nearest neighbors (kNN) graph. The resulting clusters are subsequently manually labeled using canonical marker genes. This standard workflow (PCA → kNN graph → clustering → marker-based annotation) is implemented in widely used toolkits such as Seurat and Scanpy [10, 11], and has been broadly applied across SRT platforms, including MERFISH [12, 13, 14], CosMx SMI [15], and 10x Xenium [16]. While intuitive and scalable, clustering-based annotation is highly sensitive to choices in dimensionality reduction, graph construction, and hyperparameters (e.g., resolution parameter in Leiden clustering [17, 18, 19]). Moreover, marker-based labeling requires substantial biological expertise and introduces subjectivity, leading to annotation errors that propagate to down-stream analyses.

The second approach is reference-based label transfer, which assigns cell types in the SRT datasets by aligning to a transcriptomics reference atlas with known annotations. Early approaches developed for single-cell RNA sequencing, such as correlation- or distance-based matching methods [20, 21], laid the conceptual foundation for reference-guided annotation but performed poorly on SRT data due to their limited robustness to technical and biological variation and constrained computational efficiency and scalability [22, 23]. More recent frameworks employ deep learning to achieve greater accuracy and scalability [22, 24, 25, 26]. For example, Spatial-ID [22] applies graph-based transfer learning refined using pseudo-labels from scRNA-seq pretraining, STELLAR [25] integrates graph neural networks with supervised classification and contrastive learning to incorporate spatial neighborhood information, and scBOL [26] employs a neighbor-aware prototypical learning strategy that uses local spatial context to shape the latent representation for annotation. Beyond these approaches, alignment-based methods such as Tangram [27] further enable projection of single-cell annotations onto spatial locations by learning probabilistic mappings between modalities. Collectively, these methods automate much of the annotation process and can scale efficiently to handle large datasets.

Despite these advances, reference-based methods face fundamental limitations. Their performance depends on the completeness and relevance of the chosen reference, which cannot be guaranteed in complex tissues or disease settings where emergent, transitional, or pathological cell populations are often of greatest biological interest. When novel cell types are absent from the reference, existing methods tend to misclassify them as the most similar known population, masking distinct molecular programs and obscuring biologically meaningful discoveries. This limitation is particularly consequential in disease contexts, where emerging or altered cell populations may drive pathological progression or influence treatment response [1, 28]. Although reference-based methods such as scBOL [26], Spatial-ID [22], and STELLAR [25] attempt to address this limitation by flagging cells that poorly align with known references as potential novel populations, they rely on heuristic thresholds [22] or user-defined numbers of novel clusters [25] — quantities that cannot be specified a priori in real biological settings. Moreover, none of these approaches explicitly quantify assignment uncertainty, limiting interpretability and making it difficult to distinguish ambiguous assignments from genuinely novel cell populations. These limitations highlight the need for a unified, automatic framework that preserves the strengths of reference-based annotation while enabling adaptive, uncertainty-aware discovery of novel cell populations.

To overcome these limitations, we develop **NovAST**, **Nov**el cell type discovery and **A**nnotation in **S**patial **T**ranscriptomics, a fully automated deep learning framework that jointly performs reference-based annotation and data-driven discovery of novel populations (Figure 1a). NovAST integrates reference and target cells into a shared representation while enforcing well-structured clusters for known cell types. Each target cell is assigned a quantitative confidence score reflecting its likelihood of belonging to an existing reference population. Cells that align confidently to the reference are directly annotated, whereas cells with weak alignment or exhibit inconsistent local neighborhoods are flagged as candidate novel populations. This model design enables NovAST to automatically infer whether novel cell populations are absent, singular, or multiple, a question that is fundamentally unknown in practice, thereby eliminating reliance on user-defined parameters required by existing approaches. By explicitly modeling both explained and unexplained variation, NovAST unifies reference alignment with discovery and captures continuity with known biology while revealing divergence into previously unannotated cellular states.

**Figure 1:**
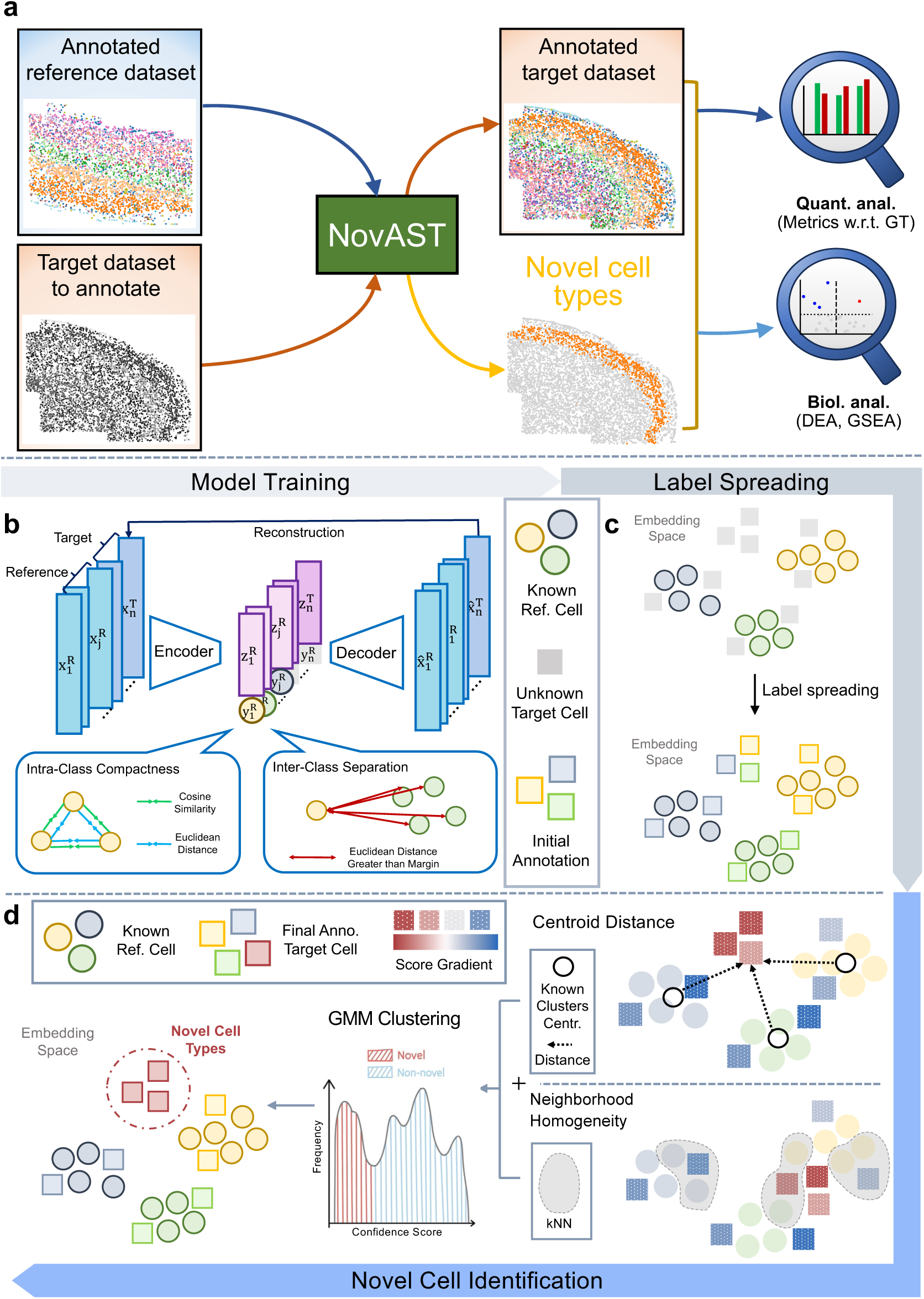
NovAST’s pipeline. **(a)** Overview of the NovAST pipeline. NovAST takes as input an annotated reference spatially resolved transcriptomics (SRT) dataset and an unannotated target SRT dataset, and outputs cell type annotations for the target, including predicted novel cell populations. The resulting annotations are evaluated quantitatively using diverse performance metrics and examined biologically through differential expression analysis (DEA) and gene set enrichment analysis (GSEA). **(b)-(d)** Schematic illustrations of each stage in detail. **(b)** Autoencoder training. The model is trained jointly on reference and target datasets for reconstruction, and additionally on the labeled reference cells to enforce intra-class compactness and inter-class separation. **(c)** Label transfer. Labels are propagated across the latent embedding space from reference to target cells via label spreading. **(d)** Novel cell identification. In the UMAP projection of the latent embedding, a confidence score is computed for each target cell based on its distance to the centroids of known cell types and the neighborhood homogeneity defined as 1/(number of existing cell types in neighborhood). Gaussian mixture model (GMM) clustering is then applied to the confidence scores, with the cluster of lowest mean designated as novel cells.

We systematically evaluated NovAST across nine reference–target dataset pairs spanning MERFISH, CosMx SMI, and 10x Xenium platforms and diverse tissue types. Across these settings, NovAST consistently achieves improved or comparable performance in cell type annotation across metrics, demonstrating its broad applicability. Beyond recovering known cell type populations, NovAST enables the identification of disease-associated novel cell populations, including immune-related cells localized to fibrotic lesions in idiopathic pulmonary fibrosis and fibroblast-like populations enriched in inflamed colonic tissue, revealing biologically meaningful structure that is not captured by existing reference-based methods. Together, these results show that NovAST supports accurate annotation of known cell types while enabling principled discovery of novel populations without relying on assumptions of reference completeness, thereby facilitating more reliable interpretation of tissue architecture and disease mechanisms. NovAST is provided as an open-source Python package (https://github.com/YMa-lab/NovAST) that integrates seamlessly with existing analysis frameworks such as Scanpy and supports the analyses presented in this study.

## 2 Results

### 2.1 Overview of NovAST

NovAST is described in Methods and its method schematic is shown in Figure 1. Briefly, NovAST is an automated, reference-based annotation framework for spatial transcriptomics that jointly performs accurate cell type annotation and novel cell type discovery. The model operates through a three-stage pipeline. First, NovAST embeds both a labeled reference dataset and an unlabeled target dataset into a shared latent embedding space, where known cell type clusters are encouraged to be compact and well-separated using a constrained loss function that incorporates cluster-structure regularization terms (Figure 1b). Second, within this latent space, NovAST propagates reference labels to target cells using a graph-based label-spreading algorithm (Figure 1c). Third, cells that fall outside of the boundaries of existing reference clusters are flagged as candidate novel cell type populations, which are subsequently refined by subclustering to reveal true novel cell types (Figure 1d). Full methodological details are provided in the Methods section.

To evaluate its performance, we systematically benchmarked NovAST against three state-of-the-art (SOTA) methods, including Tangram [27], Spatial-ID [22], and STELLAR [25], which represent alignment-, label transfer-, and discovery-based cell type annotation strategies. We tested all methods on nine reference–target dataset pairs, creating controlled “novel cell type” scenarios by withholding one annotated cell type from the reference, thereby quantifying both the global annotation accuracy across all cell types and the capacity to recover the withheld populations as novel cell types. Across all evaluation settings, NovAST achieved superior or comparable accuracy relative to existing approaches and showed particularly strong gains when abundant cell types were withheld, indicating robustness to reference incompleteness. When applied directly to unmodified real datasets, including a human Xenium dataset of idiopathic pulmonary fibrosis (IPF) and a mouse MERFISH dataset of inflamed colon, NovAST identified previously uncharacterized clusters enriched for extracellular matrix remodeling and inflammatory programs consistent with disease pathology.

### 2.2 NovAST accurately annotates mouse MERFISH MOp dataset

To provide a consistent and interpretable benchmark across datasets, we designed four representative scenarios that reflect distinct challenges in novel cell type detection (Supplementary Note 1). In each scenario, one annotated cell type was withheld from the reference and treated as a pseudo-novel population in the target, simulating a previously unobserved cell type for evaluation. We selected: (i) the Most Abundant type, representing frequent and well-sampled populations; (ii) the Least Abundant type, representing rare cell types; (iii) the Most Separated type, representing cell types that are highly distinct in the embedding space; and (iv) the Most Mixed type, representing cell types intermixed with other populations. Together, these four scenarios capture the major challenges encountered in novel cell type detection and form a unified evaluation framework applied across all datasets in this study.

We first applied this benchmark to the mouse MERFISH primary motor cortex (MOp) dataset [12]. Annotating SRT data from the cerebral cortex remains challenging due to its complex laminar structure and extensive neuronal heterogeneity, encompassing numerous distinct subtypes [29]. Specifically, the dataset comprises 258 genes across 33 slices, with an average of 4,556 cells per slice and 20 annotated cell types, including inhibitory neuron subclasses (*Pvalb*, *Sst*, and *Lamp5*), excitatory neuron subclasses (L2/3 IT, L4/5 IT, L5 IT, L6 IT, L5 ET, L5/6 NP, L6 CT, and L6b), and non-neuronal classes (Astro, Endo, Micro, Oligo, OPC, Peri, PVM, SMC, and VLMC). We divided the slices into two balanced groups, assigning one as the reference and the other as the target. Using this setup, we benchmarked NovAST against Tangram, Spatial-ID, and STELLAR across the four evaluation scenarios, with each experiment repeated 10 times to assess variance and stability.

Across all four scenarios, NovAST outperformed Tangram and Spatial-ID under all evaluation metrics for cell type annotation (ARI, Accuracy, Macro F1, and Weighted F1; ones-sided Mann–Whitney U test, *p* < 0.0001; Figure 2a, Supplementary Figure 1). Compared with STELLAR, NovAST achieves significantly higher ARI in three settings (Most Abundant, Most Separated, and Most Mixed; *p* < 0.0001), while maintaining comparable performance in the Least Abundant case (Figure 2a). Results across additional evaluation metrics were consistent with these trends (Supplementary Figure 1). Collectively, these results highlight NovAST’s ability to robustly detect novel populations across a wide range of scenarios. Representative UMAP visualizations further illustrate this behavior. NovAST’s latent representation forms compact, well-separated clusters, preserving clear margins between the reference and novel populations (Figure 2b, Supplementary Figure 2). In the Most Abundant scenario (15.70% of total cells), the pseudo-novel L2/3 IT neurons are positioned distinctly from reference clusters, enabling clean separation that competing methods fail to achieve (Figure 2b, Supplementary Figure 3a). Other scenarios showed consistent behavior: in the Most Separated and Most Mixed settings, NovAST maintained clear boundaries between reference and novel populations, while in the Least Abundant case (1.22% of total cells), it preserved competitive accuracy despite the inherent challenge of detecting extremely rare cell populations (Supplementary Figures 2). By explicitly integrating the meta-consistency and min–max feature (MMF) regularization to simultaneously contract intra-cluster distances and enforce a margin between clusters (Methods), NovAST learns a latent space in which same-type cells form markedly more compact and well-separated clusters than those produced by baseline methods, a property reflected by its substantially lower Davies–Bouldin Index (DBI) (Supplementary Figure 3), which penalizes high intra-cluster dispersion and rewards large inter-cluster separation. This improved geometric structure, in turn, enables more reliable and accurate cell type annotation. Consistent with the embedding structure, confusion matrices revealed a strong one-to-one correspondence between predicted and ground-truth cell type labels, indicating consistently high accuracy across cell types and scenarios and, critically, enabling precise delineation of the novel clusters, highlighted in the red boxes (Figure 2c, Supplementary Figure 4). In the Most Abundant setting, NovAST correctly recovered the pseudo-novel L2/3 IT cluster, whereas competing methods misassigned these cells to nearby reference types (Figure 2c). In both the Most Separated and Most Mixed settings, NovAST correctly annotates the pseudo-novel clusters, whereas the competing methods fail to do so (Supplementary Figure 4).

**Figure 2:**
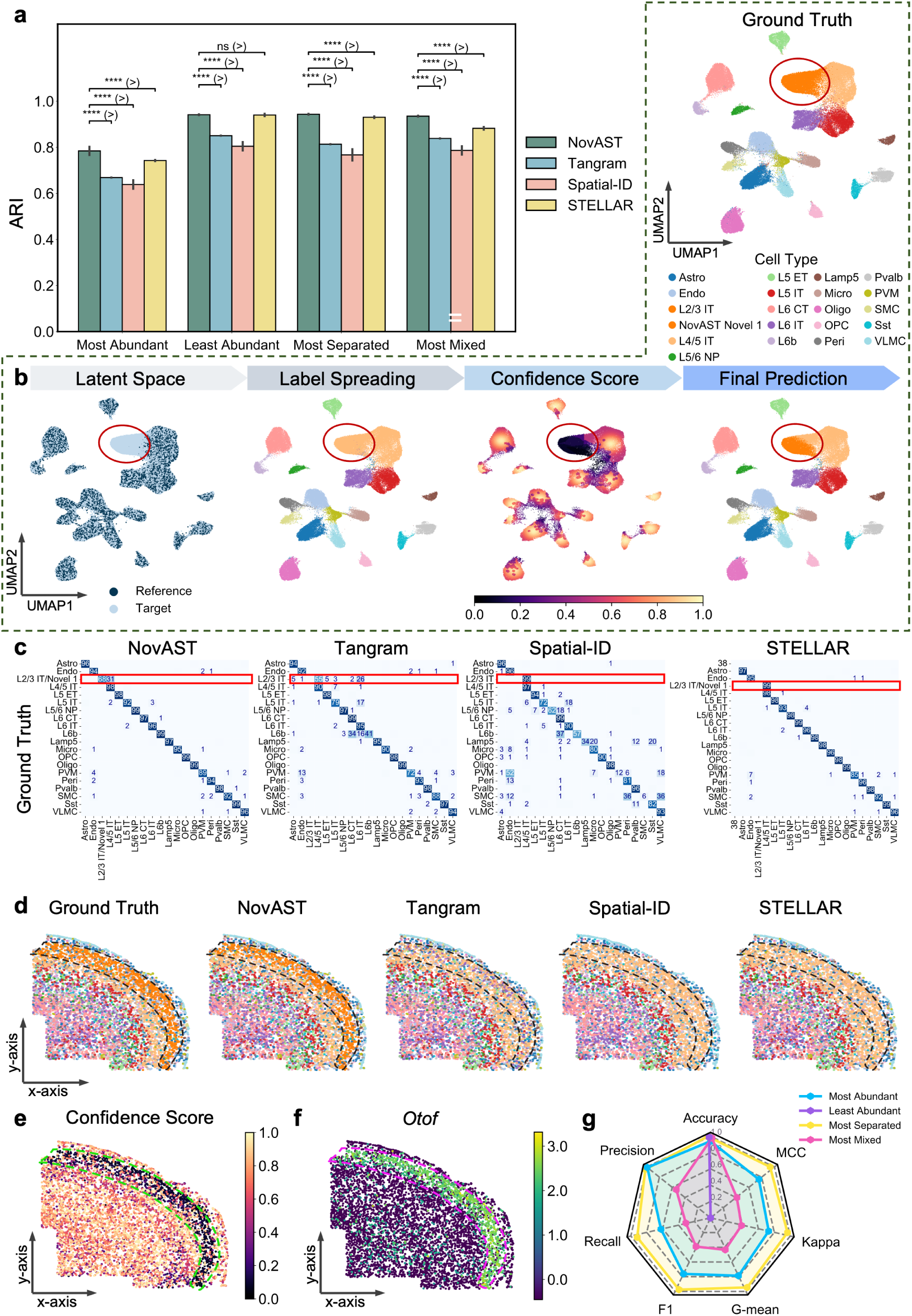
NovAST’s performance in the mouse MERFISH MOp. **(a)** ARI comparison of NovAST with three benchmarking methods (Tangram, Spatial-ID, and STELLAR) across different pseudo-novel cell type removal settings. See Methods for details on each setting. Error bars represent the standard deviation across 10 runs. Statistical significance was assessed using a one-sided Mann–Whitney U test under the alternative hypothesis that NovAST outperforms the competing method when NovAST has a higher sample mean, and vice versa; **** indicates p < 0.0001, and ‘ns’ denotes not significant. **(b)** Representative UMAP projections of the latent embedding learned by NovAST from a single run, with the Most Abundant cell type removed from the reference data. The main panel includes four horizontally arranged subpanels showing the latent space colored by reference versus target domains (titled Latent Space), label propagation results (titled Label Spreading), confidence scores ranging from 0 to 1 (titled Confidence Score), and the final predicted cell types (titled Final Prediction). An additional subpanel in the top right displays the ground truth cell type annotations for comparison (titled Ground Truth). These visualizations illustrate that NovAST produces a well-structured and representative embedding space that supports accurate label transfer and novel cell type detection. The red circle highlights the novel cell cluster in each subpanel. **(c)** Confusion matrices comparing predicted cell type labels to ground truth for NovAST and each benchmarking method for all slices. Each matrix is row-normalized based on the ground truth labels. Each entry shows the percentage of cells from the ground-truth class predicted as the label. For NovAST and STELLAR, predictions for the ‘Novel 1’ cluster were combined with the ground truth ‘L2/3 IT’ class to account for the intentionally removed reference cell type. STELLAR predicted an additional cluster ‘38’ which is not assigned to any existing cluster. A red box highlights the cell type removed from the reference and its corresponding predictions for each method. **(d)** Spatial visualization of predicted cell types from NovAST and benchmarking methods under the same setting as in (b), where the Most Abundant cell type was removed from the reference data. A single representative tissue slice (slice 71) is shown for each method. The color legend is shared with panel (c). Additional visualizations are provided in Supplementary Figures 6-7. Black dashed lines highlight the laminar region containing the novel cells. **(e)** Spatial visualization of the confidence scores computed in stage two of NovAST, shown on the same tissue slice as in panels (d). Green dashed lines highlight the laminar region containing the novel cells. **(f)** Spatial expression patterns of genes *Otof*, which are differentially expressed in the novel cell cluster identified by NovAST, shown for the same example as in panels (d) and (e). Pink dashed lines highlight the laminar region containing the novel cells. **(g)** Radar plot with seven axes (Accuracy, Precision, Recall, F1, G-mean, Cohen’s kappa, MCC). Four traces depict the settings Most Abundant, Least Abundant, Most Separated, and Most Mixed. Radial values are on a common scale ranging from 0 to 1; higher values are better.

To further assess NovAST’s ability to resolve novel cell types, we examined qualitative outputs, including a spatial map of cell types and marker gene expression patterns. The spatial distribution of NovAST-predicted cell type labels in the target data closely matches the ground truth, recapitulating the laminar organization of the cortex. Notably, NovAST uniquely identifies inhibitory neurons located in layers 2–3, whereas other methods misclassify these cells as L4/5 inhibitory neurons (Figure 2d, Supplementary Figures 5-6). The confidence scores, where lower values reflect a greater likelihood of novel-cell identity, further distinguish this population from neighboring cells (Figure 2e). This highlights NovAST’s unique ability to incorporate reliable uncertainty quantification into cell-type assignment. Moreover, NovAST uniquely identifies region-specific differentially expressed (DE) genes, such as *Otof*, which are not detected by competing methods, confirming its ability to capture biologically meaningful novel populations (Methods, Figure 2f, Supplementary Table 2).

Quantitative evaluations of novel type detection reinforce these findings (Methods). We further assessed overall robustness using seven complementary measures (i.e., Accuracy, Precision, Recall, F1, Geometric mean (G-mean), Kappa, and Matthews Correlation Coefficient (MCC)) (Methods). NovAST achieves the highest scores under the Most Separated setting, followed by the Most Abundant and Most Mixed settings, while its performance declines in the Least Abundant setting (Figure 2g). These results highlight NovAST’s ability to effectively uncover well-separated novel populations, maintain accurate annotation when novel populations are abundant or closely mixed with reference types, and remain conservative when novel cells are rare. Among all competing methods, STELLAR is the only method explicitly designed to identify novel cells. To ensure a fair evaluation, we tested its performance under multiple settings of the expected number of novel populations (Supplementary Figure 7). Although STELLAR achieved slightly higher scores when the expected number of novel populations was set to one, its performance was unstable across different parameter choices, reflecting strong sensitivity to this setting (Supplementary Figure 7a). Lower values of the number of expected cell types prevented the detection of true novel types. In contrast, higher values produced spurious false positives and further disrupted the annotation of known cell types (Supplementary Figures 7b-e and 8). In practice, this dependence poses challenges in real-world setting where the number of novel cell types is typically unknown. NovAST overcomes this limitation by eliminating user-defined parameters and leveraging a unified embedding and confidence-scoring framework, thereby enabling stable, reliable performance across datasets.

### 2.3 NovAST accurately annotates and discovers novel populations the human Xenium lung dataset

Building on its strong performance in the MERFISH dataset, we next evaluated NovAST on a Xenium dataset, which provides subcellular resolution and measures a targeted panel of 343 genes, offering a complementary evaluation on a different imaging-based SRT platform [30]. This dataset includes 30 samples from control and pulmonary fibrosis lungs. For benchmarking, we first focused on two control replicates (sample VUHD116A as reference and sample VUHD116B as target) and applied the same pseudo-novel cell type strategy as in the previous MERFISH experiments. In this dataset, the four evaluation scenarios are not independent but partially overlap across cell types. Specifically, the cell type withheld as the pseudo-novel population is the same in the Most Abundant and Most Mixed settings, and likewise for the Least Abundant and Most Separated cases. As a result, each removed population simultaneously exhibits two properties: for instance, being both abundant and highly intermixed, or rare and well separated. This natural coupling of abundance and separability increases the overall difficulty of evaluation, as methods must handle both factors concurrently rather than in isolation. Consequently, the Xenium dataset provides a particularly stringent and biologically realistic test for novel cell type detection. In line with the MERFISH results, NovAST consistently outperforms all three competing methods across all scenarios and evaluation metrics (*p* < 0.0001; Figure 3a; Supplementary Figure 9). Representative UMAP projections of the latent embedding learned by NovAST under the Most Abundant and Most Mixed settings illustrate how the model adapts to challenging scenarios in which abundant cell types are highly intermixed with others (Figure 3b). In these settings, the embedding forms a pyramid-like organization, with reference clusters occupying the vertices and the pseudo-novel population positioned in the intervening space. This structure arises from NovAST’s embedding objective, which simultaneously enforces compact intra-cluster representation and inter-cluster separation. Consistent with this behavior, NovAST achieves markedly lower DBI values than competing methods (Supplementary Figure 10). As a result, the model learns a representation that preserves continuity across related populations while maintaining sufficient distinctiveness to isolate the novel cluster, enabling informative uncertainty estimates that, in turn, support reliable novel-cell identification (Figure 3b). A similar pattern emerges in the Least Abundant and Most Separated settings, demonstrating NovAST’s consistent performance across diverse and challenging annotation scenarios (Supplementary Figure 11a). Confusion matrices further confirm this behavior, showing that NovAST achieves performance comparable to other methods on shared cell types (outside the red box) but uniquely succeeds in correctly identifying the novel population (inside the red box), thereby strengthening its overall accuracy (Figure 3c, Supplementary Figure 11b). Moreover, the spatial distribution of the predicted cell type labels (Supplementary Figures 11c & 12) shows that NovAST is the only method to correctly identify both endothelial and epithelial cells, consistent with the ground-truth annotations across all settings.

**Figure 3:**
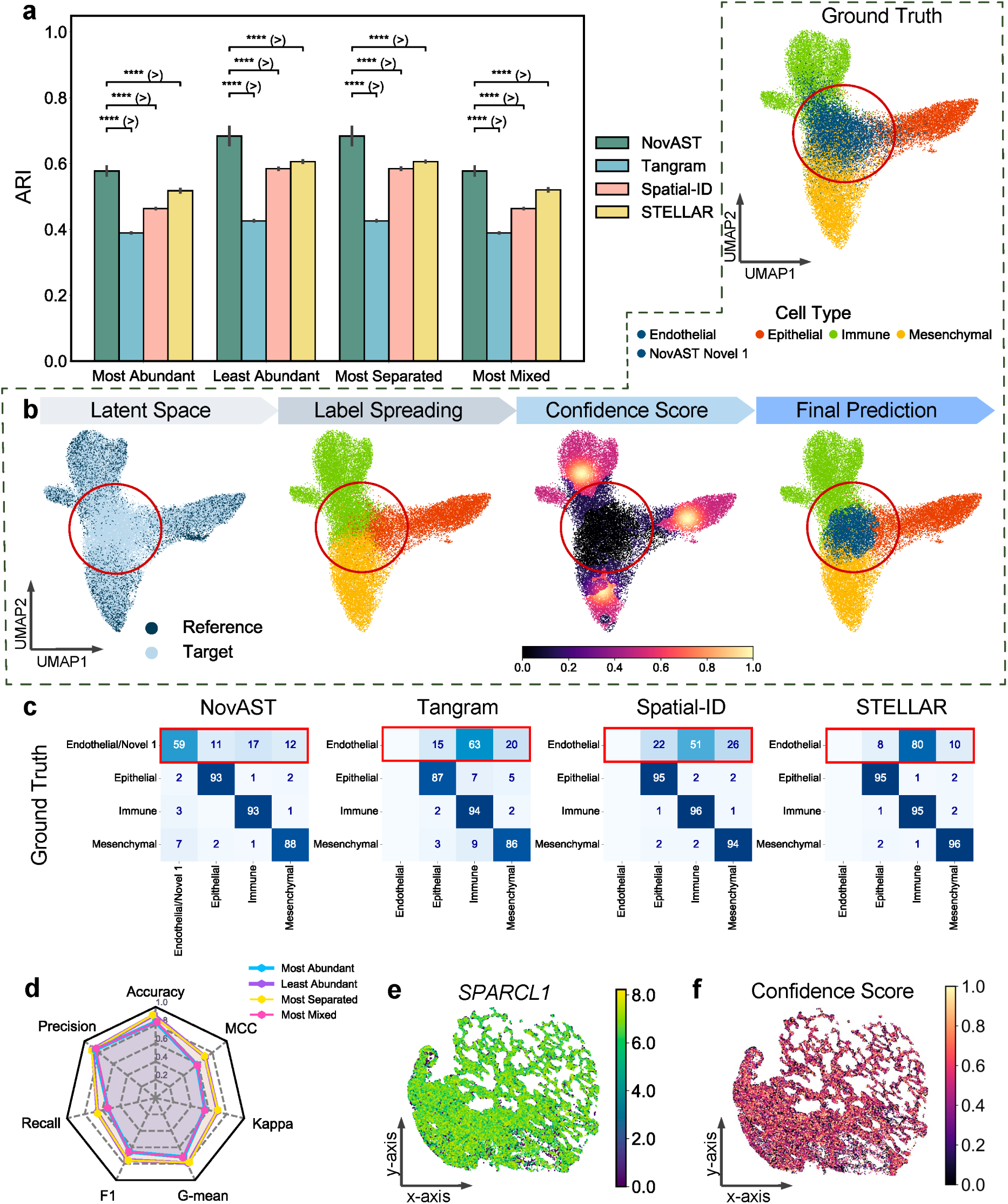
NovAST’s performance in the human Xenium lung. **(a)** ARI comparison of NovAST with three benchmarking methods (Tangram, Spatial-ID, and STELLAR) across different pseudo-novel cell type removal settings. See Methods for details on each setting. Error bars represent the standard deviation across 10 runs. Statistical significance was assessed using a one-sided Mann–Whitney U test under the alternative hypothesis that NovAST outperforms the competing method when NovAST has a higher sample mean, and vice versa; **** indicates p < 0.0001. **(b)** Representative UMAP projections of the latent embedding learned by NovAST from a single run, corresponding to both the Most Abundant and Most Mixed settings, in which Endothelial cells were removed from the reference data. The main panel includes four horizontally arranged subpanels showing the latent space colored by reference versus target domains (titled Latent Space), label propagation results (titled Label Spreading), confidence scores ranging from 0 to 1 (titled Confidence Score), and the final predicted cell types (titled Final Prediction). An additional subpanel in the top right displays the ground truth cell type annotations for comparison (titled Ground Truth). These visualizations illustrate that NovAST produces a well-structured and representative embedding space that supports accurate label transfer and novel cell type detection. The red circle highlights the novel cell cluster in each subpanel. **(c)** Confusion matrices comparing predicted cell type labels to ground truth for NovAST and each benchmarking method for all slices. Each matrix is row-normalized based on the ground truth labels. Each entry shows the percentage of cells from the ground-truth class predicted as the label. For NovAST, predictions for the ‘Novel 1’ cluster were combined with the ground truth ‘Endothelial’ class to account for the intentionally removed reference cell type. A red box highlights the cell type removed from the reference and its corresponding predictions for each method. **(d)** Radar plot with seven axes (Accuracy, Precision, Recall, F1, G-mean, Cohen’s kappa, MCC). Four traces depict the settings Most Abundant, Least Abundant, Most Separated, and Most Mixed. Radial values are on a common scale ranging from 0 to 1; higher values are better. **(e)** Spatial expression patterns of genes *SPARCL1*, which are differentially expressed in the novel cell cluster identified by NovAST, shown for the same example as in panels (b) and (c). **(f)** Spatial visualization of the confidence scores computed in stage two of NovAST, shown on the same example as in panels (b), (c), and (e).

Beyond annotation accuracy, NovAST also identifies biologically meaningful novel cell populations. Specifically, NovAST achieves consistently high performance across seven quantitative metrics (Figure 3d) and uniquely detects the endothelial cell type marker gene *SPARCL1* (Figure 3e, Supplementary Table 3) as a DE gene, with its spatial distribution concordant with the confidence scores generated by NovAST (Figure 3f). In addition, NovAST outperforms STELLAR under default settings or when the number of expected novel populations (novel seed parameter) was set to zero or one, where STELLAR identifies few to no novel cells (Supplementary Figure 13). Although STELLAR attains slightly higher scores across seven metrics when this parameter is increased, this improvement is accompanied by pronounced over-segmentation. The model partitions the data into an excessive number of small subclusters that diverge from established biological structure. This fragmentation does not isolate the novel population alone; instead, it disrupts the integrity of other well-defined cell types, leading to widespread misannotation and reduced overall coherence. By contrast, NovAST preserves stable and biologically interpretable clusters without the need for parameter tuning, reflecting its generalizability and fully automated design.

### 2.4 NovAST maintains superior accuracy on the human MERFISH developmental cortex when novel cell types encompass multiple related subtypes

In previous analyses, we evaluated model performance under settings in which either fine-grained subtype labels were used for the MERFISH mouse cortex or coarse-level labels were applied to the Xenium human lung, with only a single cell type removed in each case. To further increase difficulty, we next examined a recent MERFISH dataset of the human developmental cortex with highly resolved subtype annotations [14]. Two replicates at gestational week 34 from the primary cortical area of the parietal cortex (samples UMB5900-BA40a and UMB5900-BA40b) are designated as reference and target datasets, comprising 191,819 and 165,520 cells, respectively. Here, instead of removing a single subtype, we excluded entire coarse-level groups encompassing multiple related subtypes (Supplementary Note 3, Supplementary Figure 14).

This pseudo-novel strategy represents a more stringent test by simulating realistic scenarios in which entire families of related cell types are missing from the reference due to tissue preparation or sampling discrepancies. Consistent with the previous two datasets, NovAST outperforms Tangram and Spatial-ID across all four scenarios and evaluation metrics (p-values < 0.0001; Figure 4a, Supplementary Figure 15). Compared with STELLAR, NovAST achieved higher ARI in three settings (Most Abundant, Least Abundant, and Most Mixed) and comparable results across the remaining metrics (Figure 4a, Supplementary Figure 15). Together, these results underscore NovAST’s consistent and robust performance across increasingly challenging conditions.

**Figure 4:**
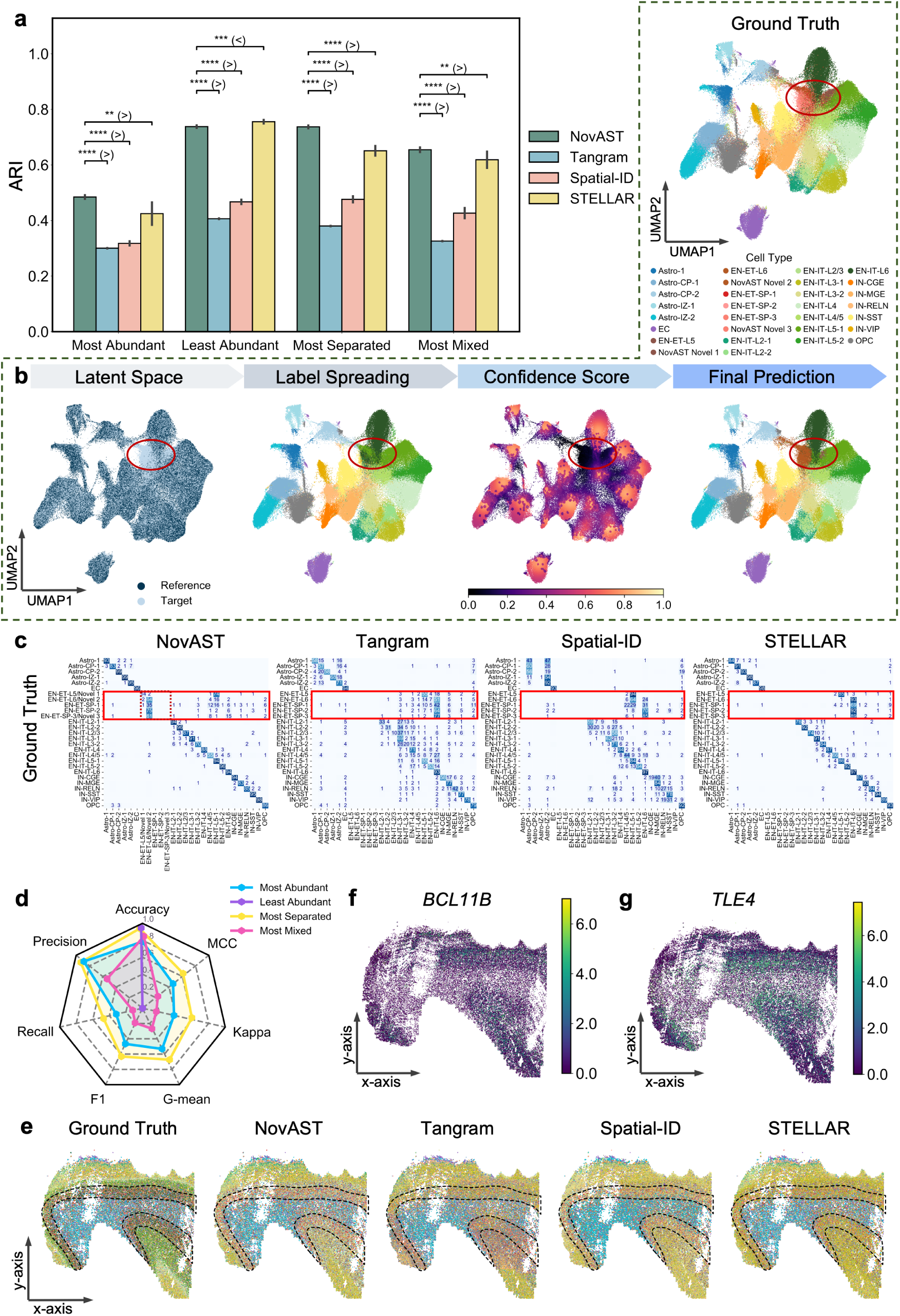
NovAST’s performance in the human MERFISH developmental cortex. **(a)** ARI comparison of NovAST with three benchmarking methods (Tangram, Spatial-ID, and STELLAR) across different pseudo-novel cell type removal settings. See Methods for details on each setting. Error bars represent the standard deviation across 10 runs. Statistical significance was assessed using a one-sided Mann–Whitney U test under the alternative hypothesis that NovAST outperforms the competing method when NovAST has a higher sample mean, and vice versa; **** indicates p < 0.0001, *** indicates p < 0.001, and ** indicates p < 0.01. **(b)** Representative UMAP projections of the latent embedding learned by NovAST from a single run, with the Most Separated cell type removed from the reference data. The main panel includes four horizontally arranged subpanels showing the latent space colored by reference versus target domains (titled Latent Space), label propagation results (titled Label Spreading), confidence scores ranging from 0 to 1 (titled Confidence Score), and the final predicted cell types (titled Final Prediction). An additional subpanel in the top right displays the ground truth cell type annotations for comparison (titled Ground Truth). These visualizations illustrate that NovAST produces a well-structured and representative embedding space that supports accurate label transfer and novel cell type detection. The red circle highlights the novel cell cluster in each subpanel. **(c)** Confusion matrices comparing predicted cell type labels to ground truth for NovAST and each benchmarking method for all slices. Each matrix is row-normalized based on the ground truth labels. Each entry shows the percentage of cells from the ground-truth class predicted as the label. For NovAST, predictions of novel clusters were combined with the ground truth class with which they most closely matched to account for the intentionally removed reference cell types. A red box highlights these cell types removed from the reference and their corresponding predictions for each method. For NovAST, an additional dashed box marks the diagonal entries of the removed cell type, illustrating that the novel cells identified by NovAST correspond to the true novel population. **(d)** Radar plot with seven axes (Accuracy, Precision, Recall, F1, G-mean, Cohen’s kappa, MCC). Four traces depict the settings Most Abundant, Least Abundant, Most Separated, and Most Mixed. Radial values are on a common scale ranging from 0 to 1; higher values are better. **(e)** Spatial visualization of predicted cell types from NovAST and benchmarking methods under the same setting as in (b), where the most separated cell type was removed from the reference data. The color legend is shared with panel (b). The black dashed line highlights the region where novel cells are predominantly aggregated. **(f)** Spatial expression patterns of genes *BCL11B*, which are differentially expressed in the novel cell cluster identified by NovAST, shown for the same example as in panel (b). **(g)** Spatial expression patterns of genes *TLE4*, which are differentially expressed in the novel cell cluster identified by NovAST, shown for the same example as in panel (b).

Representative UMAP visualizations further illustrate how NovAST achieves this improvement (Figure 4b, Supplementary Figure 16). The latent embedding learned by NovAST organizes reference and target cells into compact, well-separated manifolds, with novel populations forming distinct regions that remain proximal yet separable from related reference types (Supplementary Figure 17). During label propagation, this structure allows NovAST to accurately spread annotations while maintaining sharp cluster boundaries. The subsequent confidence map highlights low-confidence regions at the edges of novel populations, preventing erroneous assignments. The final predictions show that NovAST successfully recovers novel clusters aligned with ground-truth identities, demonstrating that its embedding and confidence scoring jointly drive its superior annotation accuracy.

Confusion matrices further validate these observations, showing that NovAST achieves markedly cleaner and more diagonal alignment between predicted and true cell type labels than competing methods (Figure 4c, Supplementary Figure 18). For non-novel cell types (outside the red box), predictions are tightly concentrated along the diagonal, indicating high annotation accuracy, whereas Tangram, Spatial-ID, and STELLAR exhibit broader dispersion and cross-class mixing. Within the red-boxed regions, corresponding to pseudo-novel populations, NovAST uniquely recovers clear and compact novel clusters that align with their ground-truth subtypes. In the Most Separated setting, two distinct novel groups are identified: Novel 1 largely overlaps with EN-ET-L5, whereas Novel 2 spans multiple related subtypes, including EN-ET-L6, EN-ET-SP-1, EN-ET-SP-2, and EN-ET-SP-3 (Figure 4c). For quantitative calculations, Novel 2 is assigned to the dominant overlapping ground-truth type (EN-ET-L6), a conservative choice that may slightly underestimate performance. Similar patterns are observed in the Most Abundant and Most Mixed settings (Supplementary Figure 17a,c & 18a,c), where NovAST delineates cohesive novel populations that other methods merge into existing classes. By contrast, no novel group is detected in the Least Abundant setting (Supplementary Figure 17b & 18b), reflecting NovAST’s conservative detection strategy that favors precision over false-positive discovery. Together, these results demonstrate that NovAST reliably identifies dominant novel populations across complex settings while maintaining stringent control against spurious detections (Figure 4d). In contrast, STELLAR exhibits variability across novel-seed settings, detecting few or none novel cells when the novel seed parameter is small (Supplementary Figures 19-20).

Importantly, the novel cells identified by NovAST preserve the laminar organization of the human cortex. The spatial maps reveal that the identified novel clusters occupy appropriate cortical layers, consistent with the ground truth (Figure 4e, Supplementary Figure 21). DE analysis of the novel cells further reveals two marker genes, *BCL11B* and *TLE4* (Methods, Figure 4f-g). These genes, which are known markers for layer 5 and the subplate, respectively [14], are only detectable through NovAST’s novel cell identification, highlighting a capability absent in the competing methods (Supplementary Table 4).

### 2.5 Quantitative evaluation on additional datasets

We further extended the quantitative evaluation of NovAST to six additional SRT datasets spanning multiple imaging-based platforms, including MERFISH, Xenium, and CosMx SMI [15, 31, 32, 16] (see Methods). Across nine benchmarks, NovAST consistently outperforms competing methods in accuracy and weighted F1 under the Most Abundant setting, the most challenging scenario across all datasets tested (Figure 5b,d). The performance gains were statistically significant relative to Tangram, Spatial-ID, and STELLAR (accuracy: *p* = [0.0019, 0.0039, 0.027]; weighted F1: *p* = [0.0020, 0.0020, 0.0059]). For the other two metrics (ARI and macro F1), NovAST significantly outperforms Tangram and Spatial-ID (all p-values < 0.05) (Figure 5a,c) and achieved similar performance with STELLAR (*p* = [0.545, 0.917]) (Figure 5a,c). Results across the other three evaluation settings remained consistent (Supplementary Figure 22). These results demonstrate that NovAST generalizes robustly across datasets and platforms. Its consistently high accuracy and weighted F1 highlight its robustness to class imbalance and its reliable annotation of existing cell types. In detecting novel populations, NovAST maintains high precision and overall accuracy while conservatively limiting false positives, yielding balanced performance across complementary metrics such as F1, G-mean, Kappa, and MCC (Supplementary Figure 23).

**Figure 5:**
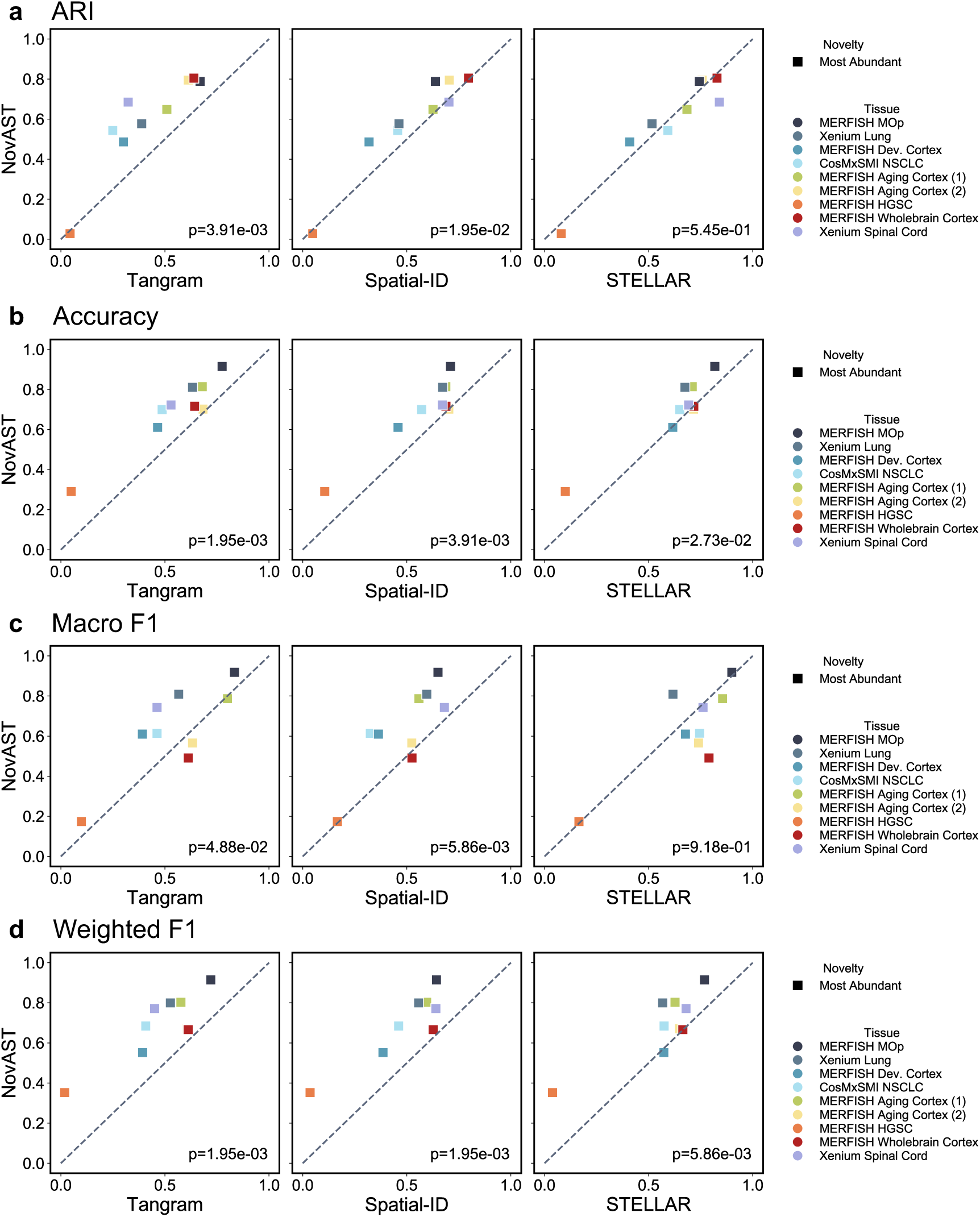
NovAST’s performance on six additional datasets. Subpanels **(a–d)** correspond to four evaluation metrics: **(a)** ARI, **(b)** Accuracy, **(c)** Macro F1, and **(d)** Weighted F1. Each subpanel contains three scatter plots, where the x-axis shows the performance of competing methods and the y-axis shows NovAST performance. Each dot represents one of nine datasets. Reported *P* values are from one-sided Wilcoxon signed-rank tests, assessing whether NovAST significantly outperforms competing methods.

### 2.6 NovAST revealed disease-associated immune patterns in human Xenium idiopathic pulmonary fibrosis dataset

Given NovAST’s consistently strong performance in benchmarking analyses, we next applied it to real SRT datasets for biological discovery, focusing on the identification of naturally occurring novel cell populations rather than pseudo-novel types. Because competing methods failed to recover novel cell types in prior evaluations, NovAST served as the primary analytical framework for downstream discovery. A common and biologically relevant setting involves comparing a healthy reference with a diseased sample, where most cell types are shared, but disease-associated populations may emerge as novel. We hypothesized that NovAST could transfer shared labels for conserved cell types and identify previously uncharacterized disease-specific states as novel cell types. To test this, we analyzed 10x Xenium SRT data from lungs affected by idiopathic pulmonary fibrosis (IPF) [30], one of the most severe interstitial lung diseases [33] characterized by aberrant epithelial injury, persistent myofibroblast activation, and excessive extracellular matrix deposition [34]. The dataset included one healthy donor (VUHD116, two replicates as reference) and one IPF donor (VUILD107, one slice as target), each assayed with a 343-gene panel, comprising 41,519 and 68,413 cells, respectively.

Consistent with previous results, NovAST produces a well-integrated latent embedding in which reference and target cells are coherently aligned, while preserving clear boundaries in major cell type clusters and revealing a distinct novel cluster adjacent to immune populations (Supplementary Figure 24a). Specifically, two clusters (i.e., Novel 1 and Novel 2) were identified near immune cells, with Novel 2 showing partial overlap with the immune cell population. Hierarchical clustering of their expression profiles confirms their close relationships with immune lineages (Supplementary Figure 24b).

To define the molecular programs underlying the novel clusters, we first performed DE analysis between IPF and healthy lungs (Methods). This analysis identified 261 significantly upregulated genes in the IPF sample, with all top 20 markers upregulated in disease. Mapping these IPF-associated genes onto the NovAST-derived clusters revealed distinct enrichments: five genes (*JCHAIN, CD79A, HERPUD1, PIM2, ITM2C*) were highly expressed in Novel 1, whereas six genes (*LUM, PTGDS, SFRP2, CXCR4, TRAC, CD44*) were highly enriched in Novel 2 (Figure 6a). Cluster-specific DE analysis further confirms these patterns (Methods, Figure 6b): four of the top ten marker genes for each cluster overlapped with the globally upregulated IPF genes (Novel 1: *JCHAIN, HERPUD1, PIM2, ITM2C*; Novel 2: *LUM, CXCR4, TRAC, CD44*), indicating that each novel cluster represents a transcriptional state closely aligned with IPF-associated molecular programs.

**Figure 6:**
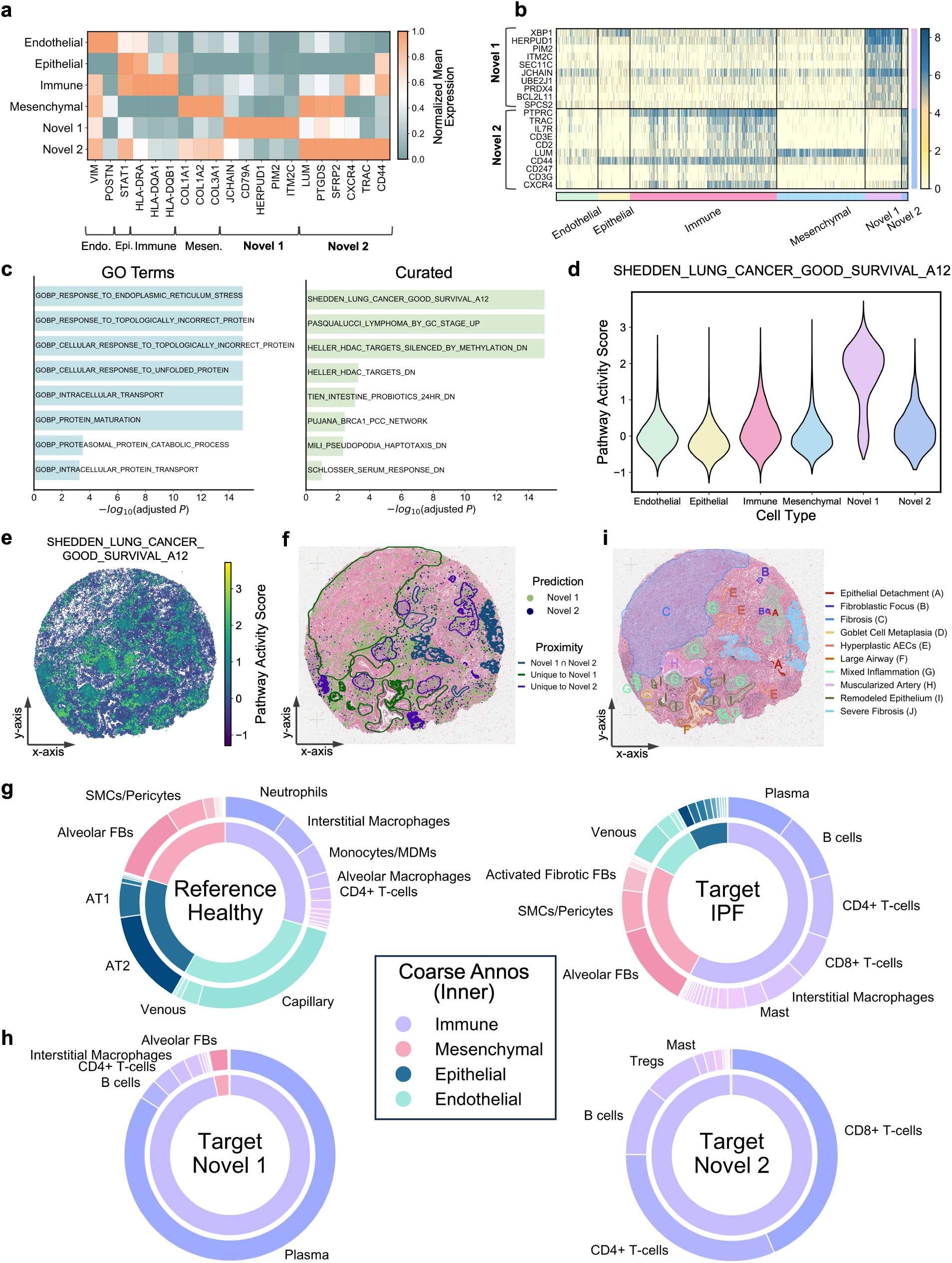
NovAST identifies disease-associated immune cells in the human Xenium IPF dataset. **(a)** Matrix plots showing the scaled average expression of marker genes across predicted cell types (rescaled to the 0–1 range per gene across cell types). Shown genes represent the top 20 markers identified by differential expression analysis (DEA) between the reference and target datasets, and are ordered by cell type according to their predominant expression. **(b)** Heatmap of gene expression clustered by predicted cell types, showing the top 10 differentially expressed (DE) genes for each of the NovAST-predicted ‘Novel 1’ and ‘Novel 2’ clusters. Rows correspond to DE genes and columns to predicted cell types, with values representing log-normalized expression. **(c)** Gene set enrichment analysis of DE genes from the Novel 1 cluster. The left panel shows enrichment against GO terms, and the right panel shows enrichment against curated datasets from the MSigDB database. Bars show the top-ranked gene sets ordered by –log_10_(adjusted *P*), with gene set names labeled on the bars. **(d)** Violin plots showing gene set activity scores for the top-ranked MSigDB pathway (`SHEDDEN_LUNG_CANCER_GOOD_SURVIVAL_A12`) across predicted cell types. Scores were computed using Scanpy’s `score_genes` function, which estimates gene set activity as the mean expression of the pathway genes relative to that of randomly selected control genes matched for average expression. **(e)** Spatial map of gene set activity scores for `SHEDDEN_LUNG_CANCER_GOOD_SURVIVAL_A12`. **(f)** Spatial plots showing the localization of ‘Novel 1’ and ‘Novel 2’ cells on the hematoxylin and eosin (H&E) image. The contour lines mark the tissue regions in which Novel 1, Novel 2, or both populations exhibit spatial proximity, with the corresponding histopathology annotations provided in panel (i). **(g)** Two-layer donut plots for reference (left) and target (right) dataset. The inner ring shows the proportion of coarse-level annotations in [30], and the outer ring indicates the proportions of fine-grained annotations, with most abundant subtypes labeled. Panels (g) and (h) share the same color legend for the coarse-level annotations. **(h)** Donut plots in the same format as (g), showing the cell type proportions of cells annotated as ‘Novel 1’ (left) and ‘Novel 2’ (right), respectively, by NovAST. **(i)** Histopathology annotations from [30] overlaid on the hematoxylin and eosin (H&E) image.

We next examined these genes in more detail. Specifically, Novel 1 exhibited a distinct expression signature characteristic of plasma cell–enriched compartments. Its top ten DE genes include *XBP1* (linked to mucus dysregulation [35]), *HERPUD1* and *SEC11C* (ER stress [36, 37]), *PIM2* (upregulated in fibrotic lesions [38]), *ITM2C* (association with plasmablast [39]), and *JCHAIN* (plasma cell enrichment [40]) (Figure 6b). Notably, *JCHAIN*, *PIM2*, and *ITM2C* have been previously identified as marker genes of plasma cells [30]. Additional markers such as *PRDX4* and *BCL2L11* point to redox imbalance and reactive oxygen species (ROS) regulation, both implicated in IPF progression [41, 42, 43]. These results highlight pathways central to IPF pathogenesis, including mucus dysregulation, ER stress, immune activation, and oxidative stress. Gene set enrichment analysis (GSEA) reinforces these findings. Novel 1 DE genes are enriched for Gene Ontology terms related to the unfolded protein response (UPR) and endoplasmic reticulum (ER) stress pathways (Figure 6c, left), and for proteostasis-related pathways such as protein maturation, intracellular transport, and proteasomal degradation. The Novel 1 markers were also enriched in a gene set signature `SHEDDEN_LUNG_CANCER_GOOD_SURVIVAL_A12` (Methods, Figure 6c, right), a lung-cancer–derived immune program in which tumor-infiltrating B and plasma cells are prognostically relevant [44, 45]. Its high activity in Novel 1 (Figure 6d) therefore suggests that fibrotic plasma-cell–like states may engage an immune-activation program reminiscent of early lung cancer, pointing to shared immunological mechanisms across cancer and fibrosis. In addition, its spatial pattern closely matched the Novel 1 region (Spearman Correlation= 0.45, *`p` ≪* 0.001, Figure 6e-f). The top three DE genes showed similar spatial enrichment (Spearman Correlations= [0.33, 0.54, 0.49], *`p` ≪* 0.001, Supplementary Figure 24c). These findings further show that Novel cluster 1 is as a plasma-cell–like population with pronounced ER-stress and immune-activation features.

Novel 2 exhibited an expression signature characteristic of T cells (Figure 6b). Its top DE genes include *PTPRC*, which regulates T-cell activation [46]; *TRAC*, a core component of T-cell receptor (TCR) [47]. Other genes included *IL7R*, which contributes to T cell homeostasis, and several parts of the T cell receptor complex (*CD3E, CD247, CD3G*) [48]. Genes such as *CD2* and *CXCR4* are involved in T cell adhesion [49] and migration [50] within tissues. Together, these markers suggest that Novel 2 represents a T cell–rich population that is likely part of the chronic immune activity seen in IPF [51].

We then examined whether these expression patterns corresponded to changes in cell composition across conditions. NovAST’s annotations captured strong differences between healthy and IPF lungs (Figure 6g-h). At the coarse main type level, the IPF sample contained a much larger immune compartment (57% versus 29%), with smaller proportions of epithelial, endothelial, and mesenchymal cells (Figure 6g, inner rings). At the finer subtype level, immune cells showed striking expansion in IPF. Plasma cells increased from 0.3% to 10.4%, B cells from 0.1% to 10.0%, CD4 T cells from 2.0% to 9.8%, and CD8 T cells from 0.7% to 6.9% (Figure 6g, right verses left, outer rings). These enriched subtypes correspond directly to the two novel clusters detected by NovAST: Novel 1 is dominated by plasma cells (84.2%), while Novel 2 consists primarily of CD8^+^ T cells (43.4%), CD4^+^ T cells (31.6%), and B cells (10.7%) (Figure 6h). These results demonstrate that NovAST captures disease-enriched immune populations that are absent in the healthy reference.

Spatial analysis provided further support. Both novel populations localized to immunerich regions (Figure 6f). These areas likely represent tertiary lymphoid structures, which contain dense groups of B cells, T cells, and plasma cells [52] (Figure 6i). Novel 1 cells appeared near muscularized arteries, large airways, goblet cell metaplasia, fibrosis, severe fibrosis, remodeled epithelium, hyperplastic alveolar epithelial cells (AECs) (*`p` <* 0.001 for Mann-Whitney U-tests, detailed values see Supplementary Table 5). These regions are associated with vascular and airway remodeling and epithelial injury. Novel 2 cells appeared in areas of mixed inflammation, epithelial detachment, severe fibrosis, fibroblastic focus, and hyperplastic AECs regions (*`p` <* 0.001, detailed values see Supplementary Table 6). These compartments are known to support chronic immune activity in IPF. The spatial patterns match observations from recent Visium studies reporting central B and T cell regions with surrounding plasma cells in IPF [53]. This spatial agreement with independent data supports that NovAST is capturing true IPF-associated immune niches.

### 2.7 NovAST discovered inflammation-associated fibroblasts and immune cells in mouse MERFISH inflamed colon data

Finally, we applied NovAST to a mouse MERFISH colon dataset of dextran sulfate sodium (DSS)-induced colitis to investigate how acute epithelial injury and immune infiltration remodel the intestinal mucosa [54]. For our analysis, Day 0 slices serve as the healthy reference and Day 9 slices at peak inflammation served as the target. The inflamed colons exhibit hallmark pathology, including crypt loss, epithelial flattening with inflammation-associated epithelial populations, and immune infiltration by monocytes, neutrophils, DC (Fcn1+), and B cells [54]. NovAST identifies two novel clusters across slices, of which Novel 1 represents the dominant population, whereas Novel 2 appeared at extremely low frequency (0.002%, fewer than five cells per slice; Supplementary Figure 25). We therefore focused downstream analyses on Novel 1.

To define the molecular identity of Novel 1, we performed DE and GSEA analyses. Comparisons between reference and target slices reveal strong upregulation of *Il1b, Sbno2, CXCL10, Mmp3, Mmp9, Acod1*, with elevated expression specifically within Novel 1 cells (Figure 7a). DE analysis of Novel 1 versus all other cells confirms these markers and highlights several overlapped DE genes (Figure 7b), including *Il1b, Acod1, Mmp9*. Notably, these genes are known markers for disease-associated cell types: *Il1b, Acod1, Mmp9* are markers for neutrophils, which are frontline innate cells rapidly mobilized to initiate inflammation [54, 55], and *Il1b* is also highly expressed in Macrophage (Cxcl10+), a macrophage subset enriched during immune activation [54, 56]. GSEA further highlighted enrichment of immune response pathways, including B cell– and leukocyte-mediated immunity, alongside extracellular matrix remodeling programs such as the NABA core matrisome (Supplementary Figure 26). These gene set signatures are consistent with the inflammatory remodeling of the colonic mucosa in the target slices, where immune infiltration and matrix reorganization represent key hallmarks of DSS-induced colitis [54, 57].

**Figure 7:**
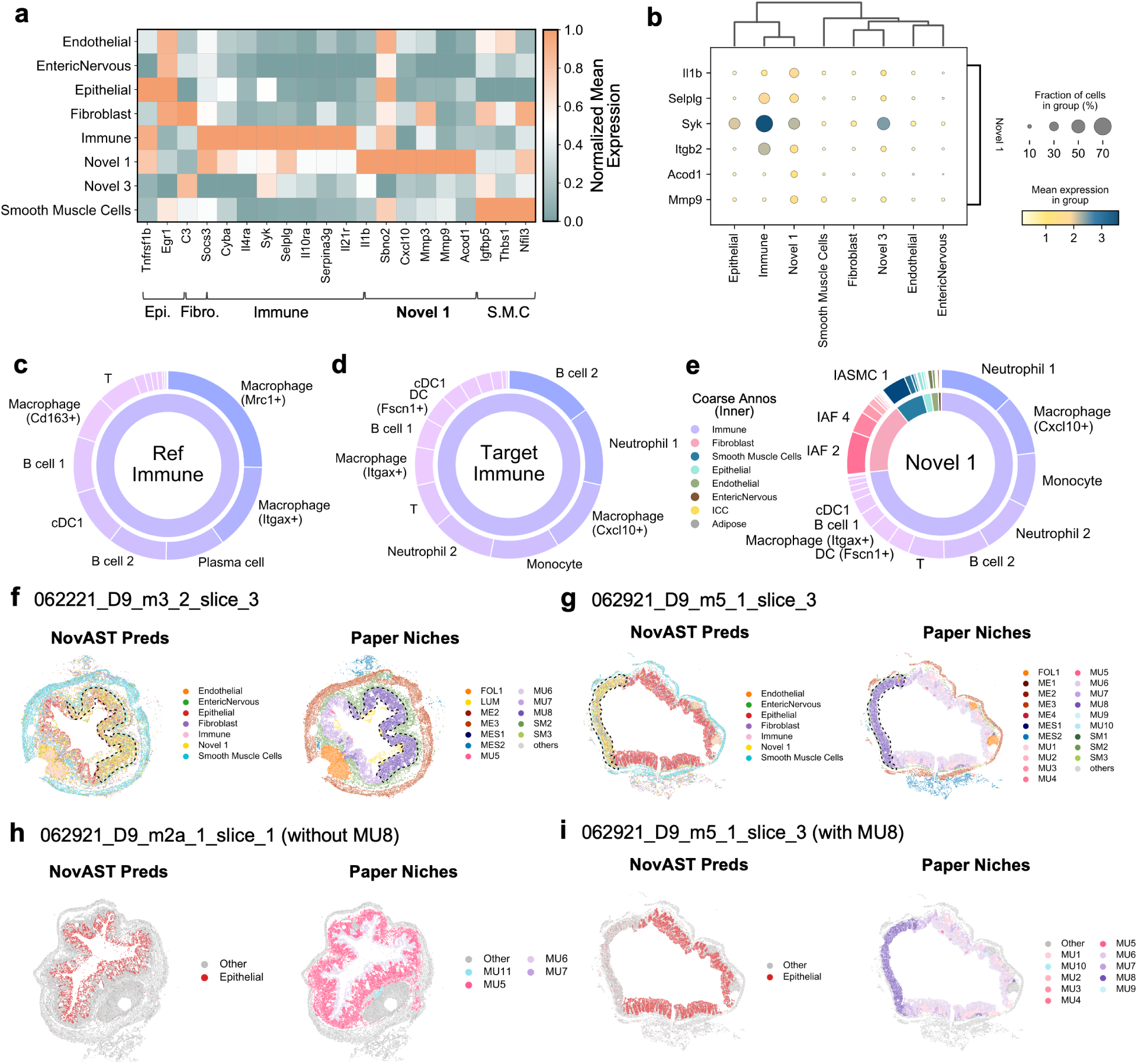
NovAST detects inflammation-related cells in the mouse MERFISH inflamed colon dataset. **(a)** Matrix plots showing the scaled average expression of marker genes across predicted cell types (rescaled to the 0–1 range per gene across cell types). Shown genes represent the top 20 markers identified by differential expression analysis (DEA) between the reference and target datasets, and are ordered by cell type according to their predominant expression. **(b)** Dot plot of top differentially expressed (DE) genes for NovAST-predicted ‘Novel 1’ cluster versus all other cell types. Columns correspond to predicted cell types and rows to the top six DE genes. Dot color represents the mean expression level, and dot size indicates the fraction of cells within each group expressing the gene. **(c-d)** Two-layer donut plots showing subtype proportions at the outer ring for immune cells in the reference **(a)** and target **(b)**. The most abundant subtypes are labeled. **(e)** Two-layer donut plots for the Novel 1 cluster identified by NovAST. The inner ring shows the proportions of coarse-level annotations, while the outer ring indicates the proportions of fine-grained annotations, with the most abundant subtypes labeled. Both annotation levels are provided in [54]. The coarse-level color legend is shown on the left. **(f-g)** Spatial plots showing the cell type predictions from NovAST (left), and the niches identified in [54] (right) for slices **(f)** `062921_D9_m3_2_slice_3` and **(g)** `062921_D9_m5_1_slice_3`. Black dashed lines highlight the region for MU8 where Novel 1 cells aggregated. **(h-i)** Spatial plots showing highlighting the epithelial cells predicted from NovAST (left), and the mucosal niches in [54] (right) for slices **(g)** `062921_D9_m2a_1_slice_1` and **(i)** `062921_D9_m5_1_slice_3`.

To verify these transcriptional findings, we next evaluated cell type composition and spatial context. Consistent with the original study [54], NovAST captures clear compositional differences between reference and target in both immune and fibroblast compartments (Figure 7c-d, Supplementary Figure 27). In the inflamed target sample, immune populations such as B cell 2, Neutrophil 1, Macrophage (Cxcl10+), Monocyte, and Neutrophil 2 are strongly enriched but nearly absent in the healthy reference (Figure 7c-d). These subtypes correspond directly to the composition of Novel 1 (Figure 7e), which also encompasses inflammation-associated fibroblast subsets (IAF2–4) uniquely present in the inflamed tissue (Supplementary Figure 27). Spatial neighborhood analysis provides further validation. In the original study [54], each cell was assigned a mucosal niche (MU) identity based on its local neighborhood composition, revealing inflammation-associated remodeling: target slices contained unique mucosal environments dominated by MU6, MU7, MU8, and LUM. Remarkably, Novel 1 cells identified by NovAST localize almost exclusively within these inflamed niches, particularly MU8 (Figure 7f-g; Supplementary Figure 28-29), confirming that the novel populations detected by NovAST arise from reference–target discrepancies in cell type composition. Additionally, the original study [54] notes that MU8 is distinguished by a nearly complete loss of epithelial layers, and NovAST’s predictions mirror this pattern: epithelial cells are correctly annotated only in MU6 and MU7 (Figure 7h-i). Even in mixed regions where MU6, MU7, and MU8 co-occur, epithelial annotations remain restricted to MU6–MU7, with none erroneously assigned to MU8 (Figure 7i). Together, these analyses demonstrate NovAST’s potential to reliably identify condition-specific cell populations and highlight their biological significance in tissue remodeling and inflammation.

## 3 Discussion

In this study, we present NovAST, an automated framework for both reference-based annotation and data-driven discovery of novel cell populations in spatial transcriptomics. Through systematic benchmarking using pseudo-novel removal and diverse reference–target pairs, NovAST consistently outperformed existing methods across nine datasets spanning multiple platforms. A key advantage of the framework is that it does not rely on user-defined parameters. This contrasts with methods such as STELLAR, whose performance depends strongly on preset expectations about the number of novel populations.

Despite these advantages, NovAST does not achieve perfect sensitivity in all scenarios. To minimize false positives, NovAST incorporates conservative design choices: clustering with Gaussian mixture models under a mean confidence threshold and an ensemble reassignment scheme that reclassifies cells that are not consistently identified as novel. While these strategies enhance precision, they reduce sensitivity, particularly for rare populations that lie close to existing clusters. The trade-off is most apparent in complex latent spaces where cluster boundaries are subtle. Future extensions may include hierarchical alignment strategies that first align broad cell classes before resolving finer subtypes, thereby better balancing precision and sensitivity in heterogeneous tissues.

NovAST also relies on UMAP-derived distances when computing confidence scores. UMAP does not fully preserve high-dimensional relationships, and distortions can arise [58, 59]. Nonetheless, we observed that UMAP-based distances yielded more stable and accurate confidence scores than Euclidean distances in the latent or PCA spaces (Supplementary Figure 36b). We attribute this improvement to UMAP’s ability to preserve both local and global neighborhood structures, which aligns with NovAST’s focus on cluster separability. Moreover, NovAST evaluates relative rather than absolute distances by comparing centroid–centroid separation with within-cluster compactness. This reliance on relative structure makes the framework less sensitive to global distortions in the embedding space and more effective at distinguishing true novel populations from boundary noise.

Together, these results establish NovAST as a robust and interpretable framework for cell type annotation and novel population discovery in spatial transcriptomics. By combining automated implementation, precision-oriented design, and broad applicability across multiple platforms, NovAST addresses key challenges in reference-based annotation. Future work will extend NovAST toward hierarchical modeling, integration with multimodal data, and application to increasingly complex tissues and disease contexts.

## 4 Methods

### 4.1 Overview of NovAST

NovAST (**Nov**el cell type discovery and **A**nnotation in **S**patial **T**ranscriptomics) is an automated deep learning framework that unifies reference-guided annotation with discovery of novel cell type populations (Figure 1). The framework operates in three stages. Specifically, in the first stage, an autoencoder constrained to learn a cluster-aware latent representation jointly embeds the cells from the reference and target datasets into a shared latent space, aligning target cells to known reference populations while preserving separability among distinctive cell type clusters through regularization that enforces compact intra-cluster structure and enlarged inter-cluster distances. In the second stage, cell type labels are propagated from the reference to the target using a graph-based label spreading algorithm that operates on the joint embedding space, enabling accurate annotation transfer. In the third stage, NovAST detects potential novel cell type populations by quantifying confidence scores and sub-clustering target cells with low assignment confidence scores. These confidence scores serve as an uncertainty-aware measure of assignment reliability, enabling NovAST to distinguish well-supported annotations from cells that deviate from known reference manifolds. This three-stage design allows NovAST to simultaneously annotate known cell types and uncover novel ones. Reference-aligned target cells are reliably classified with high confidence, while cells poorly explained by the reference are adaptively modeled as novel cell type populations. Unlike existing methods, NovAST does not require users to specify parameters such as the expected number of novel cell types. Instead, it automatically infers whether no novel cell types are present, or whether novel cells form one or several distinct clusters.

#### 4.1.1 Constrained autoencoder for representation learning and alignment

The first stage of NovAST learns a shared latent space that integrates reference and target datasets while preserving cell type cluster structure. Given a labeled reference SRT dataset *`X`*^R^ *∈* ℝ^*n*_ref_×p^ with cell type labels 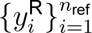 and unlabeled target SRT dataset *`X`*^T^ *∈* ℝ^*n*_tgt_×p^, NovAST jointly embeds both datasets into a shared latent space. Here, *`p`* denotes the number of overlapping genes and *`n`_ref_ `, n`_tgt_* denotes the number of reference and target cells, respectively. Both reference and target datasets are generated from imaging-based platforms (e.g., MERFISH, Xenium, and CosMx SMI), where transcriptomic profiles are aggregated at single-cell resolution.

##### Autoencoder architecture and reconstruction objective

NovAST first trains an autoencoder with cluster-structure regularization on *`X`*^R^*`, X`*^T^ to learn low-dimensional latent representations

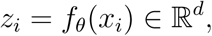

where the encoder *`f`_θ_* and decoder *`g`_ϕ_* each contain two fully connected layers with 128 hidden units, and the latent dimension is set to *`d`* = 64 by default. The decoder reconstructs the input expression profile from its latent embedding as

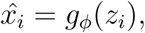

where *`x̂`_i_* denotes the reconstructed gene expression vector for cell *`i`*. The model is trained to minimize the reconstruction error between the original and reconstructed profiles using a mean-squared error loss:

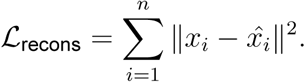

This reconstruction objective preserves global data structure across reference and target cells, ensuring that both datasets are encoded into a unified low-dimensional space. However, it does not guarantee that cells of the same type form compact clusters or that distinct types are clearly separated in the latent space (Supplementary Figure 30-32). To address this, NovAST incorporates two additional regularization terms applied to the reference embeddings *`Z`*^R^ to encourage within-type compactness and between-type separation.

##### Min–Max Feature (MMF) regularization

First, we add the Min-Max Feature (MMF) regularization, which promotes intra–cluster compactness while enforcing inter–cluster separation [60] among cell types. Intuitively, this loss encourages reference cells of the same type to aggregate closely in the latent space while pushing apart cells from different types, thereby enhancing cluster distinctiveness. Specifically, the MMF loss is defined as:

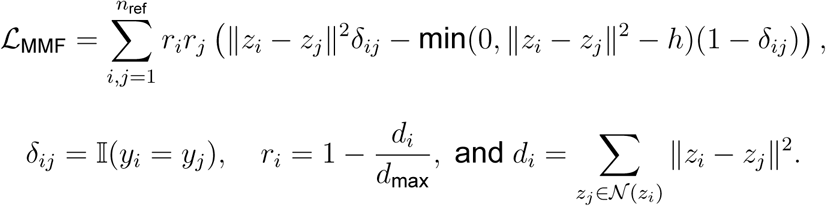

Here, ‖ · ‖ denotes the Euclidean norm; *`y`_i_* is the cell type label for reference cell *`i`*, taking values *`k`* ∈ {1*`, …, C`*} that index the known cell type classes; the indicator function *`δ`_ij_* equals to 1 if cells *`i`* and *`j`* share the same label and 0 otherwise; the term *`d`_i_* represents the sum of squared Euclidean distances from *`z`_i_* to its nearest neighbors in the latent space, using 200 neighbors by default; *`r`_i_* is a normalized local density score that assigns higher weights to cells in dense neighborhoods, thus contributing more to the regularization; and *`h`* is a predefined margin that enforces separation between different cell types, setting to 2.0 by default.

##### Meta-consistency regularization

Second, NovAST introduces a novel loss term, meta-consistency regularization (*L*_meta-cons_), which further enhances the organization of the reference embeddings by promoting directional consistency among cells of the same cell type. While MMF regularization enforces Euclidean proximity, the meta-consistency term aligns the orientations of latent vectors for cells with the same label, thereby encouraging coherent cluster geometry. For each reference cell, it computes cosine similarities to all other reference cells, aggregates the similarities by cell type, applies a softmax across classes, and minimizes the cross-entropy with the ground-truth cell type labels. The loss is defined as:

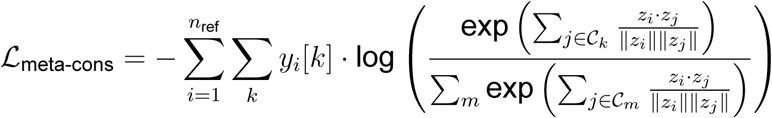

where *`C`_k_* ⊆ {1*`, …, n`*_ref_} denotes the set of reference cells with class label *`k`*, *`z`_i_* is the latent embedding of reference cell *`i`*, and *`y`_i_*[*`k`*] is the one-hot encoded annotated label indicating whether cell *`i`* belongs to class *`k`*. This design encourages cells of the same type to share similar embedding directions, reinforcing within-type homogeneity, while maintaining angular distinction across cell types. By combining MMF and meta-consistency regularizations, the model achieves spatial compactness through MMF and directional alignment through meta-consistency, producing well-separated and biologically coherent clusters that generalize effectively to target datasets.

##### Training strategy and design justification

Lastly, since reference and target cells share the same encoder, target cells are naturally mapped into nearby regions in the latent space where reference cell clusters exhibit similar expression profiles. Although the MMF and meta-consistency losses are applied only to reference cells, they indirectly guide the structure of the target embeddings, promoting alignment while preserving cell type boundaries. The overall training objective of the constrained autoencoder combines the reconstruction loss with these two regularization terms, formulated as:

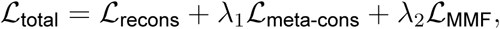

where *λ*_1_ and *λ*_2_ are weighting hyper-parameters, set by default to 1.4 and 0.1, respectively.

We train the stage-one autoencoder using the Adam optimizer with a learning rate of 1 *×* 10*^−^*^4^, weight decay of 5 *×* 10*^−^*^2^, and 50 training epochs. The batch size is set by default to 500, but can be adjusted to accommodate the sample size and GPU memory constraints. All datasets are processed using these default parameters. Ablation studies demonstrate that each component of the total loss, including reconstruction, MMF, and meta-consistency, contributes critically to model performance, and the selected hyperparameters yield optimal or near-optimal results across datasets (Supplementary Figure 33-34, Supplementary Note 2). Specifically, evaluation using the Davies–Bouldin Index (DBI) confirms that the complete model architecture produces compact and well-separated clusters, reflecting improved intra-cluster cohesion and inter-cluster distinction (Supplementary Figure 35, Supplementary Note 2). Furthermore, fully connected encoder–decoder layers were adopted instead of graph attention networks (GATs) based on spatial coordinates, as ablation analyses revealed that the fully connected design yields superior performance, stability, and generalization across diverse datasets (Supplementary Figure 33d, Supplementary Note 2).

#### 4.1.2 Graph-based label spreading for annotation transfer

The second stage of NovAST transfers cell type labels from the reference to the target dataset within the learned latent space. Here, reference embeddings *`Z`^R^* are labeled, while target embeddings *`Z`^T^* are initially unlabeled. Because Stage 1 aligns both datasets in a shared latent representation, known cell types form mixed clusters of reference and target cells, providing a natural setting for label propagation, whereby annotations from labeled reference cells diffuse through the latent geometry and are assigned to nearby unlabeled target cells. We adopt a graph-based label spreading framework [61], which assumes that nearby cells in the latent space are likely to share the same type. Specifically, we construct a sparse affinity matrix *`W`* using a *`k`*-nearest neighbors (k-NN) graph on all embeddings, where each cell is connected only to its *`k`*-nearest neighbors in the latent space, where *`k`* = 100 by default. The affinity matrix *`W`* is symmetrically normalized as *`S`* = *`D`^−^*^1/2^*`WD`^−^*^1/2^, where *`D`* is the diagonal degree matrix. Label spreading is then performed by solving the fixed-point equation:

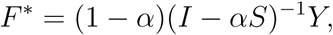

where *`Y`* encodes one-hot labels for reference cells and zeros for target cells, and *`α`* in (0, 1) controls the trade-off between label fidelity and smoothness across the graph. Each target cell is assigned the label corresponding to the maximum value in its row of *`F` ^∗^*. To accommodate large SRT datasets, we implement an efficient approximate neighbor search using hierarchical navigable small world (HNSW) graphs via the `hnswlib` package [62], replacing the exact k-NN graph when the number of cells exceeds 100, 000. This substantially reduces runtime and memory requirements while maintaining comparable accuracy (Supplementary Figure 36). Under these optimized conditions, the Stage 2 hyperparameters achieve near-optimal performance across all datasets in the ablation study (Supplementary Figure 37, Supplementary Note 2).

By leveraging both local neighborhood relationships and the global manifold structure, Stage 2 achieves robust label transfer, propagating reference annotations to target cells while respecting the geometry learned in Stage 1. This step establishes the foundation for distinguishing well-aligned known types from candidate novel populations.

#### 4.1.3 Confidence scoring-based detection and clustering of novel cells

The third stage of NovAST aims to identify target cells that may not correspond to any known reference cell type. While Stage 2 propagates labels from reference to target cells, some assignments are inherently uncertain. For example, cells located far from all reference clusters in the latent space, or cells embedded in neighborhoods dominated by mixed cell type labels, may not be well explained by any known cell type. Such cells represent natural candidates for novel populations.

To systematically identify these cells, NovAST assigns each target cell a composite confidence score that integrates two complementary measures: (1) *centroid proximity*, which quantifies how closely a cell aligns with known reference clusters, and (2) *neighborhood label homogeneity*, which assesses whether its local neighborhood is consistently aligned with a single reference label. Cells that are simultaneously distant from all reference centroids and surrounded by heterogeneous neighbors are deemed low-confidence and are flagged as potential novel cells. Specifically, these two components are calculated as follows:

##### Centroid proximity

Latent embeddings **z***_i_* are projected into two dimensions using UMAP [63] to obtain 2D UMAP coordinates **z̃**_*i*_ ∈ ℝ^2^. For each reference cell type *`j`*, a centroid *C_j_* is defined as the mean of its 2D embeddings

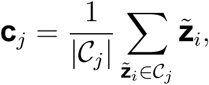

where *C_j_* is the set of reference cells of cell type *`j`*. For each target cell *`i`*, the Euclidean distance to centroid *`j`* is computed as *`d`_i,j_* = ‖ **z̃***_i_−***c***_j_*‖, and converted into a similarity weight using a Gaussian kernel (*`σ`* = 1.0):

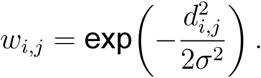

Weights are then summed across all reference types to obtain a single proximity score:

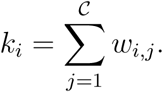

Because the raw values of *k_i_* depend on dataset size and scale, the proximity scores are normalized to the range [0,1]:

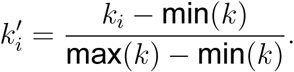

Thus, 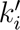 reflects the relative proximity of cell *i* to known centroids, where higher values indicate closer alignment with one or more reference cell type, while lower values indicate weak similarity to all known cell types.

##### Neighborhood label homogeneity

For each target cell *`i`*, NovAST identifies its *`k`*-nearest reference neighbors in the UMAP space (default *`k`* = 15 for datasets with fewer than 100,000 cells and *`k`* = 30 otherwise). Let *`T`_i_* = {*`t`*_1_*`, …, t`_k_*} denote the labels of these neighbors, and *`U`_i_* = unique(*`T`_i_*) the set of distinct reference labels. The homogeneity score is then defined as the inverse of the number of unique labels:

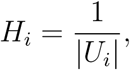

which approaches 1 when all of a target cell’s neighbors share a single label and decreases toward 0 as the neighborhood becomes more heterogeneous. As with proximity, we normalize homogeneity scores across all cells for comparability:

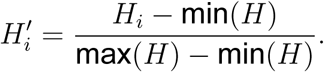

Thus, 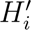 captures the degree of local label consistency, where higher values indicate coherent neighborhoods aligned with one reference type, whereas lower values indicate mixed or ambiguous neighborhoods.

##### Composite confidence score and clustering

The overall confidence score for target cell *i* is then computed as the average of the two normalized components:

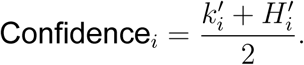

Here, 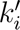 reflects how well a cell aligns with known cell type centroids, while 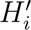 captures the local consistency of its neighborhood labels. Cells with high confidence scores are therefore both close to a known reference cluster and embedded within a homogeneous neighborhood, indicating reliable annotation. Conversely, cells with low confidence scores are distant from all reference clusters or situated in label-heterogeneous regions, suggesting potential novelty.

To delineate novel populations, NovAST first applies the Gaussian Mixture Model (GMM) clustering to the distribution of confidence scores. Minority components with mean values below a predefined threshold (default 0.12) are designated as low-confidence groups. Within these groups, we then apply Density-Based Spatial Clustering of Applications with Noise (DBSCAN) [64] to detect locally dense regions, which are interpreted as candidate novel populations. Finally, to mitigate spurious detections and ensure reproducibility, NovAST adopts an ensemble consensus strategy: the full pipeline is repeated across multiple random seeds, and a cell is retained as novel only if consistently identified in at least half of the runs (default 10 repetitions). This conservative design minimizes spurious detections and ensures that the identified novel populations represent coherent, reproducible biological signals rather than isolated low-confidence outliers.

#### 4.2 Compared baseline methods

We benchmarked with three baseline methods: Tangram, Spatial-ID, and STELLAR. Tangram [27] performs cell-to-spot alignment between reference scRNA-seq data to SRT datasets, subsequently transferring the reference cell type labels to the target data. We run Tangram following their GitHub tutorial (https://github.com/broadinstitute/Tangram). However, because Tangram learns a mapping matrix between reference and target cells via non-convex optimization and lacks a multi-threaded GPU implementation, it cannot efficiently handle large datasets due to both memory constraints and slow computational speed. To address this limitation in large-sample datasets that Tangram cannot process as a whole, we first determine whether the data originate from a single slice or multiple slices. If multiple slices are present, each slice naturally forms its own chunk. If the dataset consists of a single slice, we randomly partition it into appropriately sized chunks, testing a hierarchical set of partition numbers (10, 20, 50, 100, 500, 1,000) to balance computational efficiency and spatial resolution. We then annotate cells for each reference-target subset pair and average the model performance across subsets. Spatial-ID [22] is a deep learning framework that cell type labels from scRNA-seq to SRT datasets by generating pseudo-labels from the reference and learning spatial embeddings of the target data through a variational graph autoencoder (VGAE). We run it following their GitHub tutorial at https://github.com/STOmics/SpatialID. STELLAR [25] is a geometric deep learning framework designed to annotate a target SRT dataset given a reference SRT dataset, with the capability to identify novel cell type clusters when the hyperparameter `num-seed-class` (the expected number of novel clusters) is set greater than zero. Notably, the model architecture is flexible with respect to this parameter, such that the number of predicted novel clusters does not necessarily match the specified value. We followed their GitHub tutorial default settings (https://github.com/snap-stanford/stellar), and varied the number of novel cell types to be 1, 5, and 9. However, we observed that STELLAR failed on large datasets due to the brute-force calculation of pairwise distance matrices (*`O`*(*`n`*^2^) complexity) in its nearest neighbor search. To address this scalability issue, we replaced the distance computation with scikit-learn’s `NearestNeighbors` (with `algorithm=“auto”`), which adaptively selects efficient tree-based algorithms, without altering the remainder of STELLAR’s workflow.

#### 4.3 Data collection and preprocessing

We evaluated NovAST on eleven reference-target dataset pairs of SRT data spanning MERFISH, Xenium, and CosMx SMI technologies, encompassing cortex, lung, spinal cord, and colon tissues from both human and mouse samples, as summarized in Supplementary Table 1. We obtained the gene count matrices at the single-cell level, together with the corresponding spatial coordinates and the cell type annotations provided by the original study as ground truth. For each pair, we maintained the genes that are measured in both reference and target datasets. We then applied a prepossessing pipeline for each dataset individually: the counts were normalized to adjust for library size, the expression values were log-transformed with a pseudo-count one to stabilize the variance, and gene expression was subsequently standardized.

#### 4.4 Benchmarking metrics on cell type annotation performance

We evaluated NovAST and baseline methods using four complementary metrics, including **Accuracy**, **Adjusted Rand Index (ARI)**, **Macro F1**, and **Weighted F1**. When novel clusters are predicted, we first construct a confusion matrix between the pseudo-novel cell types and the predicted novel clusters, and assign each pseudo-novel type to the best-matching predicted cluster. Specifically, **Accuracy** measures the proportion of correctly classified cells, and **Adjusted Rand Index (ARI)** [65] quantifies the similarity between predicted clusters and the ground truth cell types. In addition, the F1 score balances precision and recall for each class. Because multiple classes are present, we aggregated the F1 scores of each class in two ways. The **Macro F1** [66] is the unweighted average of per-class F1 scores, and Macro F1 emphasizes performance on minority classes by giving all classes the same weight. In contrast, the **Weighted F1** weights each class’ F1 score by its number of true samples, so reflecting overall performance while accounting for class imbalance by giving more influence to larger classes. All four metrics were computed using `scikit-learn` [67], and their mathematical formulations are provided in Supplementary Note 3.

To further assess the detection of truly novel cells, we treated novel cell identification as a binary classification problem (novel vs. non-novel) and evaluated seven performance metrics: **Accuracy**, **Precision**, **Recall**, **F1-score**, **Geometric Mean (G-mean)**, **Kappa**, and the **Matthews Correlation Coefficient (MCC)**. Detailed definitions of these metrics are also provided in Supplementary Note 3.

Additionally, **Davies–Bouldin Index (DBI)** scores are used to evaluate the compactness and separation of the embedding spaces learned by NovAST and the baseline methods (i.e., Spatial-ID and STELLAR), with metric details provided in Supplementary Note 3.

#### 4.5 Differential gene analysis and gene set enrichment analysis

To characterize molecular differences uncovered by NovAST, We performed two types of differential expression (DE) analyses: (1) between cells from reference and target datasets (Figure 6a, Figure 7a), and (2) between each novel cell cluster and all remaining cell clusters within the target dataset (Figure 6b, Figure 7b. Both analyses were conducted using the `rank_genes_groups` function in Scanpy with the Wilcoxon rank-sum test, applied to log-normalized expression values. For each comparison, genes were ranked by the test statistics, and we selected the top *n* significant genes (with adjusted p-values < 0.05) for subsequent plotting.

We next compared NovAST with competing methods in identifying marker genes specific to novel clusters (Figure 2f, Figure 3e, Figure 4f-g). For NovAST, DE analysis was performed between the novel cluster and all other cells. For each competing method, we identified the predicted cluster showing the largest overlap with the true novel cells—typically an existing known type—and performed the same DEA contrast. We then compared the top 20 significant DE genes from these analyses and identified those unique to NovAST.

Finally, we performed gene set enrichment analysis (GSEA) on NovAST-identified novel clusters using the `gseapy` [68] implementation of the `prerank` function. Genes were ranked by log fold change from the DEA comparing novel clusters to all other cells. Enrichment was assessed using gene sets from the Molecular Signatures Database (MSigDB; https://www.gsea-msigdb.org/gsea/msigdb/index.jsp), including both the Gene Ontology (GO) collection and the curated collection (Reactome, KEGG, and other pathway resources) for human and mouse. Gene set activity was quantified using `scanpy.tl.score_genes`, which computes a cell-level enrichment score by comparing the average expression of the genes in the set to a matched set of control genes with similar expression levels. Scores were computed on log-normalized expression values.

## 5 Data availability

A summary of evaluated datasets are included in Supplementary Table 1. The public datasets analyzed in this study are available from the following sources: the MERFISH mouse primary motor cortex (MOp) dataset from the Brain Image Library (https://doi.brainimagelibrary.org/doi/10.35077/g.21); the human Xenium lung (pulmonary fibrosis, PF) dataset from GEO under accession GSE250346 (https://www.ncbi.nlm.nih.gov/geo/query/acc.cgi?acc=GSE250346); the human MERFISH developmental cortex dataset from Zenodo (https://doi.org/10.5281/zenodo.14422018); the CosMx SMI human non-small cell lung cancer (NSCLC) dataset from Zenodo (https://zenodo.org/records/15240431); the MERFISH mouse cortex aging study (24–90 weeks) from the CellxGene portal (https://cellxgene.cziscience.com/collections/31937775-0602-4e52-a799-b6acdd2bac2e); the MERFISH tubo-ovarian high-grade serous carcinoma (HGSC) dataset from the Single Cell Portal (https://singlecell.broadinstitute.org/single_cell/study/SCP2650/hgsc-spatial-cohort-discovery-dataset); the MERFISH whole mouse brain dataset from CellxGene (https://cellxgene.cziscience.com/collections/0cca8620-8dee-45d0-aef5-23f032a5cf09); the Xenium human spinal cord dataset from Zenodo (https://doi.org/10.5281/zenodo.8037425); and the mouse MERFISH inflamed colon dataset from Dryad (https://doi.org/10.5061/dryad.rjdfn2zh3).

## 6 Code availability

NovAST is implemented in Python and is available on GitHub: https://github.com/YMa-lab/NovAST.

## 7 Acknowledgments

We gratefully acknowledge Winston Li for contributing the initial formulation of the meta-consistency loss.

## 9 Author Contributions Statement

Y.M. conceived the idea, and designed the study. Y.Z and Y.M. designed the benchmarking and real data experiments in the manuscript. Y.Z led the method development, with supervision from Y.M. and R.S.. Y.Z. implemented the NovAST software, performed real data analysis, benchmarking experiments, and prepared the figures. Y.Z, and Y.M. wrote the manuscript, with input from R.S..

## References

[1] Changde Cheng, Wenan Chen, Hongjian Jin, and Xiang Chen. A review of single-cell rna-seq annotation, integration, and cell–cell communication. Cells, 12(15), 2023.

[2] Tianhao Li, Zixuan Wang, Yuhang Liu, Sihan He, Quan Zou, and Yongqing Zhang. An overview of computational methods in single-cell transcriptomic cell type annotation. Briefings in Bioinformatics, 26(3):bbaf207, 05 2025.

[3] Yuhan Hao, Tim Stuart, Madeline H Kowalski, Saket Choudhary, Paul Hoffman, Austin Hartman, Avi Srivastava, Gesmira Molla, Shaista Madad, Carlos Fernandez-Granda, et al. Dictionary learning for integrative, multimodal and scalable single-cell analysis. Nature biotechnology, 42(2):293–304, 2024.

[4] Dylan M Cable, Evan Murray, Vignesh Shanmugam, Simon Zhang, Luli S Zou, Michael Diao, Haiqi Chen, Evan Z Macosko, Rafael A Irizarry, and Fei Chen. Cell type-specific inference of differential expression in spatial transcriptomics. Nature methods, 19(9):1076–1087, 2022.

[5] Ying Ma, Shiquan Sun, Xuequn Shang, Evan T Keller, Mengjie Chen, and Xiang Zhou. Integrative differential expression and gene set enrichment analysis using summary statistics for scrna-seq studies. Nature communications, 11(1):1585, 2020.

[6] Lulu Shang, Peijun Wu, and Xiang Zhou. Statistical identification of cell type-specific spatially variable genes in spatial transcriptomics. Nature communications, 16(1):1059, 2025.

[7] Luz Garcia-Alonso, Louis-François Handfield, Kenny Roberts, Konstantina Nikolakopoulou, Ridma C Fernando, Lucy Gardner, Benjamin Woodhams, Anna Arutyunyan, Krzysztof Polanski, Regina Hoo, et al. Mapping the temporal and spatial dynamics of the human endometrium in vivo and in vitro. Nature genetics, 53(12):1698–1711, 2021.

[8] Suoqin Jin, Christian F Guerrero-Juarez, Lihua Zhang, Ivan Chang, Raul Ramos, Chen-Hsiang Kuan, Peggy Myung, Maksim V Plikus, and Qing Nie. Inference and analysis of cell-cell communication using cellchat. Nature communications, 12(1):1088, 2021.

[9] Volker Bergen, Marius Lange, Stefan Peidli, F Alexander Wolf, and Fabian J Theis. Generalizing rna velocity to transient cell states through dynamical modeling. Nature biotechnology, 38(12):1408–1414, 2020.

[10] Satija Lab. Seurat guided clustering tutorial of 3k pbmcs. https://satijalab.org/seurat/articles/pbmc3k_tutorial.html, 2025. Accessed: 2025-09-01.

[11] Scanpy developers. Scanpy tutorial: Clustering and manual cell type annotation. https://scanpy.readthedocs.io/en/stable/tutorials/basics/clustering.html#manual-cell-type-annotation, 2025. Accessed: 2025-09-01.

[12] Meng Zhang, Stephen W. Eichhorn, Brian Zingg, Zizhen Yao, Kaelan Cotter, Hongkui Zeng, Hongwei Dong, and Xiaowei Zhuang. Spatially resolved cell atlas of the mouse primary motor cortex by merfish. Nature, 598(7879):137–143, 2021.

[13] Rongxin Fang, Chenglong Xia, Jennie L. Close, Meng Zhang, Jiang He, Zhengkai Huang, Aaron R. Halpern, Brian Long, Jeremy A. Miller, Ed S. Lein, and Xiaowei Zhuang. Conservation and divergence of cortical cell organization in human and mouse revealed by merfish. Science, 377(6601):56–62, 2022.

[14] Xuyu Qian, Kyle Coleman, Shunzhou Jiang, Andrea J Kriz, Jack H Marciano, Chunyu Luo, Chunhui Cai, Monica Devi Manam, Emre Caglayan, Abbe Lai, et al. Spatial transcriptomics reveals human cortical layer and area specification. Nature, pages 1–11, 2025.

[15] Tancredi Massimo Pentimalli, Simon Schallenberg, Daniel León-Periñán, Ivano Legnini, Ilan Theurillat, Gwendolin Thomas, Anastasiya Boltengagen, Sonja Fritzsche, Jose Nimo, Lukas Ruff, et al. Combining spatial transcriptomics and ecm imaging in 3d for mapping cellular interactions in the tumor microenvironment. Cell systems, 16(5), 2025.

[16] Petra Kukanja, Christoffer M Langseth, Leslie A Rubio Rodríguez-Kirby, Eneritz Agirre, Chao Zheng, Amitha Raman, Chika Yokota, Christophe Avenel, Katarina Tiklova, Andre O Guerreiro-Cacais, et al. Cellular architecture of evolving neuroinflammatory lesions and multiple sclerosis pathology. Cell, 187(8):1990–2009, 2024.

[17] Monika Krzak, Yordan Raykov, Alexis Boukouvalas, Luisa Cutillo, and Claudia Angelini. Benchmark and parameter sensitivity analysis of single-cell rna sequencing clustering methods. Frontiers in Genetics, 10:1253, 2019.

[18] Felix Raimundo, Celine Vallot, and Jean-Philippe Vert. Tuning parameters of dimensionality reduction methods for single-cell rna-seq analysis. Genome Biology, 21:212, 2020.

[19] Xiao Liang, Lijie Cao, Hao Chen, Lidan Wang, Yangyun Wang, Lijuan Fu, Xiaqin Tan, Enxiang Chen, Yubin Ding, and Jing Tang. A critical assessment of clustering algorithms to improve cell clustering and identification in single-cell transcriptome study. Briefings in Bioinformatics, 25(1):bbad497, 01 2024.

[20] Vladimir Yu Kiselev, Andrew Yiu, and Martin Hemberg. scmap: projection of single-cell rna-seq data across data sets. Nature methods, 15(5):359–362, 2018.

[21] Dvir Aran, Agnieszka P Looney, Leqian Liu, Esther Wu, Valerie Fong, Austin Hsu, Suzanna Chak, Ram P Naikawadi, Paul J Wolters, Adam R Abate, et al. Reference-based analysis of lung single-cell sequencing reveals a transitional profibrotic macrophage. Nature immunology, 20(2):163–172, 2019.

[22] Rongbo Shen, Lin Liu, Zihan Wu, Ying Zhang, Zhiyuan Yuan, Junfu Guo, Fan Yang, Chao Zhang, Bichao Chen, Wanwan Feng, et al. Spatial-id: a cell typing method for spatially resolved transcriptomics via transfer learning and spatial embedding. Nature communications, 13(1):7640, 2022.

[23] Jinming Cheng, Xinyi Jin, Gordon K Smyth, and Yunshun Chen. Benchmarking cell type annotation methods for 10x xenium spatial transcriptomics data. BMC bioinformatics, 26(1):22, 2025.

[24] Quyuan Tao, Yiheng Xu, Youzhe He, Ting Luo, Xiaoming Li, and Lei Han. Benchmarking mapping algorithms for cell-type annotating in mouse brain by integrating single-nucleus rna-seq and stereo-seq data. Briefings in Bioinformatics, 25(4):bbae250, 05 2024.

[25] Maria Brbić, Kaidi Cao, John W Hickey, Yuqi Tan, Michael P Snyder, Garry P Nolan, and Jure Leskovec. Annotation of spatially resolved single-cell data with stellar. Nature Methods, 19(11):1411–1418, 2022.

[26] Yuyao Zhai, Liang Chen, and Minghua Deng. scbol: a universal cell type identification framework for single-cell and spatial transcriptomics data. Briefings in Bioinformatics, 25(3):bbae188, 04 2024.

[27] Tommaso Biancalani, Gabriele Scalia, Lorenzo Buffoni, Raghav Avasthi, Ziqing Lu, Aman Sanger, Neriman Tokcan, Charles R Vanderburg, Åsa Segerstolpe, Meng Zhang, et al. Deep learning and alignment of spatially resolved single-cell transcriptomes with tangram. Nature methods, 18(11):1352–1362, 2021.

[28] Kevin H Wong, Natalia Andrade Rodriguez, and Nikki Traylor-Knowles. Exploring the unknown: How can we improve single-cell rnaseq cell type annotations in non-model organisms? Integrative and Comparative Biology, 64(5):1291–1299, 07 2024.

[29] Namyoung Jung and Tae-Kyung Kim. Spatial transcriptomics in neuroscience. Experimental & molecular medicine, 55(10):2105–2115, 2023.

[30] Annika Vannan, Ruqian Lyu, Arianna L Williams, Nicholas M Negretti, Evan D Mee, Joseph Hirsh, Samuel Hirsh, Niran Hadad, David S Nichols, Carla L Calvi, et al. Spatial transcriptomics identifies molecular niche dysregulation associated with distal lung remodeling in pulmonary fibrosis. Nature genetics, 57(3):647–658, 2025.

[31] William E Allen, Timothy R Blosser, Zuri A Sullivan, Catherine Dulac, and Xiaowei Zhuang. Molecular and spatial signatures of mouse brain aging at single-cell resolution. Cell, 186(1):194–208, 2023.

[32] Christine Yiwen Yeh, Karmen Aguirre, Olivia Laveroni, Subin Kim, Aihui Wang, Brooke Liang, Xiaoming Zhang, Lucy M Han, Raeline Valbuena, Michael C Bassik, et al. Mapping spatial organization and genetic cell-state regulators to target immune evasion in ovarian cancer. Nature immunology, 25(10):1943–1958, 2024.

[33] Shaney L Barratt, Andrew Creamer, Conal Hayton, and Nazia Chaudhuri. Idiopathic pulmonary fibrosis (ipf): an overview. Journal of clinical medicine, 7(8):201, 2018.

[34] Qianru Mei, Zhe Liu, He Zuo, Zhenhua Yang, and Jing Qu. Idiopathic pulmonary fibrosis: an update on pathogenesis. Frontiers in pharmacology, 12:797292, 2022.

[35] Gang Chen, Carla MP Ribeiro, Ling Sun, Kenichi Okuda, Takafumi Kato, Rodney C Gilmore, Mary B Martino, Hong Dang, Aiman Abzhanova, Jennifer M Lin, et al. Xbp1s regulates muc5b in a promoter variant–dependent pathway in idiopathic pulmonary fibrosis airway epithelia. American journal of respiratory and critical care medicine, 200(2):220–234, 2019.

[36] Koichi Kokame, Kishan Lal Agarwala, Hisao Kato, and Toshiyuki Miyata. Herp, a new ubiquitin-like membrane protein induced by endoplasmic reticulum stress*. Journal of Biological Chemistry, 275(42):32846–32853, 2000.

[37] A Manuel Liaci, Barbara Steigenberger, Paulo Cesar Telles de Souza, Sem Tamara, Mariska Gröllers-Mulderij, Patrick Ogrissek, Siewert J Marrink, Richard A Scheltema, and Friedrich Förster. Structure of the human signal peptidase complex reveals the determinants for signal peptide cleavage. Molecular cell, 81(19):3934–3948, 2021.

[38] Hisao Higo, Kadoaki Ohashi, Shuta Tomida, Sachi Okawa, Hiromasa Yamamoto, Seiichiro Sugimoto, Satoru Senoo, Go Makimoto, Kiichiro Ninomiya, Takamasa Nakasuka, et al. Identification of targetable kinases in idiopathic pulmonary fibrosis. Respiratory research, 23(1):20, 2022.

[39] Sudhir Ghandikota and Anil G. Jegga. From gene networks to therapeutics: A causal inference and deep learning approach for drug discovery. Pharmaceuticals, 18(9), 2025.

[40] Guiquan Jia, Thirumalai R Ramalingam, Jason Vander Heiden, Xia Gao, Daryle DePianto, Katrina B Morshead, Zora Modrusan, Nandhini Ramamoorthi, Paul Wolters, Celia Lin, et al. An interleukin 6 responsive plasma cell signature is associated with disease progression in systemic sclerosis interstitial lung disease. Iscience, 26(11), 2023.

[41] Tetsuya Hanaka, Takashi Kido, Shingo Noguchi, Sohsuke Yamada, Hirotsugu Noguchi, Xin Guo, Aya Nawata, Ke-Yong Wang, Keishi Oda, Tsutomu Takaki, et al. The overexpression of peroxiredoxin-4 affects the progression of idiopathic pulmonary fibrosis. BMC Pulmonary Medicine, 19(1):265, 2019.

[42] Weiwei Zhu, Chunquan Liu, Chunting Tan, and Jie Zhang. Predictive biomarkers of disease progression in idiopathic pulmonary fibrosis. Heliyon, 10(1):e23543, 2024.

[43] Judith Hagenbuchner, Andrey Kuznetsov, Martin Hermann, Barbara Hausott, Petra Obexer, and Michael J. Ausserlechner. Foxo3-induced reactive oxygen species are regulated by bcl2l11 (bim) and sesn3. Journal of Cell Science, 125(5):1191–1203, 03 2012.

[44] Si-si Wang, Wei Liu, Dalam Ly, Hao Xu, Limei Qu, and Li Zhang. Tumor-infiltrating b cells: their role and application in anti-tumor immunity in lung cancer. Cellular & molecular immunology, 16(1):6–18, 2019.

[45] Dapeng Hao, Guangchun Han, Ansam Sinjab, Lorena Isabel Gomez-Bolanos, Rossana Lazcano, Alejandra Serrano, Sharia D Hernandez, Enyu Dai, Xuanye Cao, Jian Hu, et al. The single-cell immunogenomic landscape of b and plasma cells in early-stage lung adenocarcinoma. Cancer discovery, 12(11):2626–2645, 2022.

[46] Maryam Ahmed Al Barashdi, Ahlam Ali, Mary Frances McMullin, and Ken Mills. Protein tyrosine phosphatase receptor type c (ptprc or cd45). Journal of Clinical Pathology, 74(9):548–552, 2021.

[47] National Center for Biotechnology Information (NCBI). Trac: T cell receptor alpha constant [homo sapiens (human)]. https://www.ncbi.nlm.nih.gov/gene?Db=gene&Cmd=DetailsSearch&Term=28755, 2025. Retrieved October 16, 2025, from the NCBI Gene Database.

[48] Matthew E Call and Kai W Wucherpfennig. The t cell receptor: critical role of the membrane environment in receptor assembly and function. Annu. Rev. Immunol., 23(1):101–125, 2005.

[49] Bin Li, Yan Lu, Ming-Chao Zhong, Jin Qian, Rui Li, Dominique Davidson, Zhenghai Tang, Kaiwen Zhu, Jérémy Argenty, Anne Gonzalez de Peredo, Bernard Malissen, Romain Roncagalli, and André Veillette. Cis interactions between cd2 and its ligands on t cells are required for t cell activation. Science Immunology, 7(74):eabn6373, 2022.

[50] Rita Lucia Contento, Barbara Molon, Cedric Boularan, Tullio Pozzan, Santos Manes, Stefano Marullo, and Antonella Viola. Cxcr4–ccr5: A couple modulating t cell functions. Proceedings of the National Academy of Sciences, 105(29):10101–10106, 2008.

[51] Xiaobo Sun, Xinwen Zhang, Yuhan He, Xueting Du, Qian Cai, and Zhihong Liu. Cd4+t and cd8+t cells profile in lung inflammation and fibrosis: targets and potential therapeutic drugs. *Frontiers in Immunology*, Volume 16–2025, 2025.

[52] Catherine Sautès-Fridman, Florent Petitprez, Julien Calderaro, and Wolf Herman Fridman. Tertiary lymphoid structures in the era of cancer immunotherapy. Nature Reviews Cancer, 19(6):307–325, 2019.

[53] Zhi Yang, Gaoyuan Cao, Xiaosheng Tan, Sarah Orfanos, Joseph Jude, Gaetan Barbet, Steven S. An, Dianhua Jiang, Reynold A. Panettieri, and Qi Yang. Distinct mural cells and fibroblasts promote pathogenic plasma cell accumulation in idiopathic pulmonary fibrosis. European Respiratory Journal, 65(6), 2025.

[54] Paolo Cadinu, Kisha N. Sivanathan, Aditya Misra, Rosalind J. Xu, Davide Mangani, Evan Yang, Joseph M. Rone, Katherine Tooley, Yoon-Chul Kye, Lloyd Bod, Ludwig Geistlinger, Tyrone Lee, Randall T. Mertens, Noriaki Ono, Gang Wang, Liliana Sanmarco, Francisco J. Quintana, Ana C. Anderson, Vijay K. Kuchroo, Jeffrey R. Moffitt, and Roni Nowarski. Charting the cellular biogeography in colitis reveals fibroblast trajectories and coordinated spatial remodeling. Cell, 187(7):2010–2028, April 2024.

[55] Harry L. Malech, Frank R. DeLeo, and Mark T. Quinn. *T*he Role of Neutrophils in the Immune System: An Overview, pages 3–10. Humana Press, Totowa, NJ, 2014.

[56] Cheng Peng, Guangping Tu, Jiale Wang, Yilin Wang, Peng Wu, Li Yu, Zhiqiang Li, and Xiao Yu. Mlkl signaling regulates macrophage polarization in acute pancreatitis through cxcl10. Cell Death & Disease, 14(2):155, 2023.

[57] Chiara Ippolito, Rocchina Colucci, Cristina Segnani, Mariella Errede, Francesco Girolamo, Daniela Virgintino, Amelio Dolfi, Erika Tirotta, Piero Buccianti, Giulio Di Candio, Daniela Campani, Maura Castagna, Gabrio Bassotti, Vincenzo Villanacci, Corrado Blandizzi, and Nunzia Bernardini. Fibrotic and vascular remodelling of colonic wall in patients with active ulcerative colitis. Journal of Crohn’s and Colitis, 10(10):1194–1204, 03 2016.

[58] Sebastian Damrich and Fred A Hamprecht. On umap’s true loss function. Advances in Neural Information Processing Systems, 34:5798–5809, 2021.

[59] Hyeon Jeon, Jeongin Park, Sungbok Shin, and Jinwook Seo. Stop misusing t-sne and umap for visual analytics. arXiv preprint arXiv:2506.08725, 2025.

[60] Weiwei Shi, Yihong Gong, Chris Ding, Zhiheng Ma, Xiaoyu Tao, and Nanning Zheng. Transductive semi-supervised deep learning using min-max features. In Vittorio Ferrari, Martial Hebert, Cristian Sminchisescu, and Yair Weiss, editors, Computer Vision – ECCV 2018, pages 311–327, Cham, 2018. Springer International Publishing.

[61] Dengyong Zhou, Olivier Bousquet, Thomas Navin Lal, Jason Weston, and Bernhard Schölkopf. Learning with local and global consistency. In Advances in neural information processing systems, volume 16, pages 321–328, 2003.

[62] Yu A Malkov and Dmitry A Yashunin. Efficient and robust approximate nearest neighbor search using hierarchical navigable small world graphs. IEEE transactions on pattern analysis and machine intelligence, 42(4):824–836, 2018.

[63] Leland McInnes, John Healy, and James Melville. Umap: uniform manifold approximation and projection for dimension reduction. arxiv. arXiv preprint arXiv:1802.03426, 10, 2018.

[64] Martin Ester, Hans-Peter Kriegel, Jörg Sander, Xiaowei Xu, et al. A density-based algorithm for discovering clusters in large spatial databases with noise. In *kdd*, volume 96, pages 226–231, 1996.

[65] Lawrence Hubert and Phipps Arabie. Comparing partitions. Journal of classification, 2(1):193–218, 1985.

66. Juri Opitz and Sebastian Burst. Macro f1 and macro f1. arXiv preprint arXiv:1911.03347, 2019.

[67] Fabian Pedregosa, Gaël Varoquaux, Alexandre Gramfort, Vincent Michel, Bertrand Thirion, Olivier Grisel, Mathieu Blondel, Peter Prettenhofer, Ron Weiss, Vincent Dubourg, Jake Vanderplas, Alexandre Passos, David Cournapeau, Matthieu Brucher, Matthieu Perrot, and Édouard Duchesnay. Scikit-learn: Machine learning in Python. Journal of Machine Learning Research, 12:2825–2830, 2011.

[68] Zhuoqing Fang, Xinyuan Liu, and Gary Peltz. Gseapy: a comprehensive package for performing gene set enrichment analysis in python. Bioinformatics, 39(1):btac757, 11 2022.

