## Supplementary Materials for "NovAST: a deep learning framework for automated label transfer and novel cell type discovery in spatial transcriptomics"

**Supplementary Figures** **8**

**Supplementary Tables** **50**

### Supplementary Notes

#### Supplementary Note 1: Design of pseudo-novel evaluation scenarios

To rigorously evaluate NovAST, we designed a benchmarking strategy that systematically varied the difficulty of novel cell type detection. Using a mouse MERFISH primary motor cortex (MOp) dataset [1], we constructed pseudo-novel settings by withholding one cell type at a time from the reference and assessing whether it could be rediscovered as novel in the target. Performance was averaged over 10 replicates (mean F1) (Methods). We then investigated two factors that could influence detection difficulty: (1) cell-type abundance in the target dataset and (2) separability, defined as the mean centroid distance of that cell type to all others in the UMAP embedding. Both factors showed some linear associations with performance (Supplementary Figure 38a-b; $R^2 = 0.12$ for frequency and $R^2 = 0.18$ for distance), characterized by consistently lower performance for rare cell types and improved accuracy with greater spatial separation. Guided by this, we evaluated four representative scenarios, (1) Most Abundant, (2) Least Abundant, (3) Most Separated, and (4) Most Mixed, to cover a broad range of frequency–separation combinations.

#### Supplementary Note 2: Details in ablation studies and parameter sensitivity analysis

We systematically evaluated the contribution of major model components and parameter choices to NovAST's overall performance using three datasets: mouse MERFISH MOp, human Xenium lung, and human MERFISH developmental cortex under all four pseudo-novel scenarios (Most Abundant, Least Abundant, Most Separated, and Most Mixed). Model performance is assessed using four metrics: ARI, Accuracy, Macro F1, and Weighted F1 (Supplementary Figures 33-37).

Supplementary Figure 33a-c quantify the importance of each loss component by individually removing the reconstruction, meta-consistency, and min–max feature (MMF), and comparing performance to NovAST trained with the full constrained autoencoder. Across datasets and evaluation metrics, removal of any individual loss term leads to a noticeable decline in performance in at least one dataset, indicating that all three objectives contribute synergistically to embedding quality and accurate identification of novel cell populations. Supplementary Figures 30-32 further visualize the embedding space, showing that the full model architecture produces more compact intra-type aggregation and improved inter-type separation across cell populations. The Davies–Bouldin Index (DBI) (Supplementary Note 4) quantitatively supports these observations across all pseudo-novel scenarios (Supplementary Figure 35), based on pairwise distances computed from UMAP coordinates, which are subsequently used for neighborhood construction in novel cell identification. In addition, Supplementary Figure 33d evaluates the impact of incorporating spatial information by comparing models using graph attention layers (GAT) on a $k$-nearest neighbor (kNN) graph derived from spatial coordinates versus fully connected layers without spatial input. Incorporating spatial graphs led to a significant decline in NovAST's performance, likely due to the over-smoothing effect of message passing in GATs (Supplementary Figure 39). Therefore, we adopted the fully connected architecture without explicit spatial information as the default configuration for subsequent analyses.

We next examined how key training parameters influence overall performance, including those used during the training of the constrained autoencoder (Supplementary Figure 34) and during the novel cell type identification stage (Supplementary Figure 37). For the training of the constrained autoencoder, we systematically varied the latent dimension size, number of hidden layers, margin parameter for the MMF loss, and the number of nearest neighbors ($k$) used within MMF. Performance remains stable across a broad range of latent dimensions (32-256) (Supplementary Figure 34a) and exhibits modest variation with network depths, with the default two-layer architecture providing the best overall balance in performance across datasets (Supplementary Figure 34b). MMF-related parameters exhibits more pronounced effects: overly large margins led to over-smoothing and a decline in performance (Supplementary Figure 34c), whereas the number of neighbors ( $k$) has only a minor influence, with a slight preference for larger values, likely due to the large sample sizes of SRT datasets (Supplementary Figure 34d). Optimal parameters are selected based on balanced performance across datasets.

During the novel cell type identification stage, we systematically varied the number of $k$-nearest neighbors for neighborhood label homogeneity estimation, the Gaussian kernel band width for centroid proximity calculation, the number of GMM components for novel cluster detection, and the minimum samples parameter in DBSCAN used for sub-clustering refinement. Each parameter is tested once for a range of candidate values. Overall, performance remains stable within moderate parameter ranges. Specifically, NovAST achieves optimal performance at $50 - 100$ $k$-nearest neighbors (Supplementary Figure 37a), $1.0 - 2.0$ band widths (Supplementary Figure 37b), $6 - 9$ GMM components (Supplementary Figure 37c), and $100 - 200$ minimum samples for DBSCAN (Supplementary Figure 37d). These analyses demonstrate that NovAST's performance is robust to parameter variation and does not rely on narrow hyperparameter tuning, highlighting its generalizability across datasets.

We further evaluated the sensitivity of NovAST to graph construction and coordinate representations. Specifically, we compared an approximate graph generated using the `hnswlib` package [2] with an exact $k$-nearest neighbor (KNN) graph during label spreading, and assessed different coordinate spaces (UMAP, PCA, and the original latent space) for neighbor generation in the confidence score calculation (Supplementary Figure 36). The approximate graph achieves comparable performance to the exact graph while substantially reducing computational time (Supplementary Figure 36a). Additionally, although using UMAP coordinates yield a modest improvement in ARI for the human Xenium lung dataset, NovAST demonstrates overall stable performance across neighborhood constructed from different coordinate spaces (Supplementary Figure 36b).

##### Supplementary Note 3: Details of pseudo-novel cell removal from the human MERFISH developmental cortex dataset

In the human MERFISH developmental cortex dataset, we evaluated a more challenging scenario in which multiple related sub-cell types belonging to the same coarse-level lineage were removed, while the coarse-level lineage itself continued to be selected according to the abundance or spatial separation criteria of each pseudo-novel setting. For example, in the Most Separated setting, the extratelencephalic excitatory neuron (EN-ET) group was removed (Figure 14c, left), including EN-ET-L5, EN-ET-L6, and three subplate-specific ET subtypes (EN-ET-SP-1, EN-ET-SP-2, EN-ET-SP-3) (Figure 14c, right). Similarly, intratelencephalic excitatory neurons (EN-IT-1), glial and endothelial cells (ECs), and inhibitory neurons (INs) were excluded for the Most Abundant, Least Abundant, and Most Mixed settings, respectively (Figure 14a,b,d).

**Supplementary Note 4: Details of evaluation metrics**

In our benchmarking analysis, we evaluated NovAST and benchmarking methods using four metrics for overall cell type annotation, which are **Accuracy**, **Adjusted Rand Index (ARI)**, **Macro F1**, and **weighted F1**, and seven additional metrics assessing performance in novel cell identification, including **Accuracy**, **Precision**, **Recall**, **F1-score**, **Geometric Mean (G-mean)**, **Kappa**, and **Matthews Correlation Coefficient (MCC)**. The **Davies–Bouldin Index (DBI)** is used to quantitatively evaluate cluster structure in comparison with benchmarking methods and in the ablation studies, demonstrating that NovAST's superior performance arises from its markedly more compact within-cluster structure and greater between-cluster separation. Detailed formulations of all metrics are provided below.

The four overall cell type annotation metrics are:

- **Accuracy** is calculated as

$$\text{Accuracy} = \frac{1}{N}\sum_{i=1}^{N}\mathbb{1}(\hat{y}_i = y_i),$$

  where $N$ is the total number of cells, $y_i$ is the ground truth label, and $\hat{y}_i$ is the predicted label. Accuracy ranges from 0 to 1, with higher values indicating better performance.

- **Adjusted Rand Index (ARI)** [3] is calculated as

$$\text{ARI} = \frac{\sum_{ij}\binom{n_{ij}}{2} - \left[\sum_i \binom{a_i}{2}\sum_j \binom{b_j}{2}\right]/\binom{n}{2}}{\frac{1}{2}\left[\sum_i \binom{a_i}{2} + \sum_j \binom{b_j}{2}\right] - \left[\sum_i \binom{a_i}{2}\sum_j \binom{b_j}{2}\right]/\binom{n}{2}},$$

  where $n_{ij}$ is the number of samples in both true cluster $i$ and predicted cluster $j$, $a_i = \sum_j n_{ij}$, $b_j = \sum_i n_{ij}$, and $n = \sum_{ij} n_{ij}$. ARI ranges from $-1$ to $1$, where $1$ indicates perfect agreement and higher values denote better clustering consistency.

- **Macro F1** [4] is the unweighted average of per-class F1 scores such that

$$\text{Macro F1} = \frac{1}{\mathcal{C}}\sum_{k=1}^{\mathcal{C}}\text{F1}_k,$$

  where $k$ is the class label and $\mathcal{C}$ is the total number of classes. Specifically, the F1 score is calculated as

$$\text{F1} = \frac{2 \cdot \text{Precision} \cdot \text{Recall}}{\text{Precision} + \text{Recall}},$$

where
$$\text{Precision} = \frac{\text{TP}}{\text{TP} + \text{FP}}, \quad \text{Recall} = \frac{\text{TP}}{\text{TP} + \text{FN}}.$$

Here, TP, TN, FP, and FN denote the number of true positives, true negatives, false positives, and false negatives, respectively. Macro F1 ranges from $0$ to $1$, with higher values indicating better balanced performance across classes.

- **Weighted F1** weights each class F1 score by its number of true samples $n_k$,

$$\text{Weighted F1} = \frac{1}{N} \sum_{k=1}^{\mathcal{C}} n_k \cdot \text{F1}_k, \quad N = \sum_{k=1}^{\mathcal{C}} n_k.$$

  Weighted F1 ranges from $0$ to $1$, with higher values reflecting better overall classification performance while accounting for class imbalance.

The seven metrics for evaluation of novel cell identification are:

- **Accuracy** measures the overall proportion of correctly classified samples:

$$\text{Accuracy} = \frac{\text{TP} + \text{TN}}{\text{TP} + \text{TN} + \text{FP} + \text{FN}}.$$

  Accuracy reflects overall correctness but ignores class imbalance, as the majority class dominates its value.

- **Precision** quantifies the proportion of predicted novel cells that are truly novel:

$$\text{Precision} = \frac{\text{TP}}{\text{TP} + \text{FP}}.$$

  High precision means that most predicted novel cells are truly novel, which indicates that predictions of novel cells are reliable.

- **Recall** (or Sensitivity) quantifies the proportion of true novel cells that were correctly identified:

$$\text{Recall} = \frac{\text{TP}}{\text{TP} + \text{FN}}.$$

  High recall means that most true novel cells are captured, which reflects comprehensive detection of novel cells.

- **F1** is the harmonic mean of precision and recall:

$$\text{F1} = \frac{2 \cdot \text{Precision} \cdot \text{Recall}}{\text{Precision} + \text{Recall}}.$$

  F1 score balances precision and recall, and is informative in our setting since the classes are imbalanced.

- **G-mean** is the geometric mean of recall for novel cells (sensitivity) and recall for non-novel cells (specificity):

$$\text{G-mean} = \sqrt{\frac{\text{TP}}{\text{TP} + \text{FN}} \cdot \frac{\text{TN}}{\text{TN} + \text{FP}}},$$

  which balances performance between the two classes, as it is maximized only when both recalls are high, making it well suited for imbalanced classification.

- **Kappa** (short for Cohen's Kappa) [5] measures agreement between predictions and ground truth, adjusted for agreement expected by chance:

$$\kappa = \frac{p_o - p_e}{1 - p_e},$$

  where $p_o = \frac{\text{TP+TN}}{N}$ is the observed accuracy and $p_e = \frac{\text{(TP+FP)(TP+FN)+(FN+TN)(FP+TN)}}{N^2}$ is the expected accuracy by random chance. Equivalently, in terms of the confusion matrix entries,

$$\kappa = \frac{2 \cdot (\text{TP} \cdot \text{TN} - \text{FP} \cdot \text{FN})}{(\text{TP} + \text{FP}) \cdot (\text{FP} + \text{TN}) + (\text{TP} + \text{FN}) \cdot (\text{FN} + \text{TN})}.$$

  Kappa therefore quantifies chance-corrected agreement, making it more robust than raw accuracy in imbalanced settings.

- **MCC** (short for Matthews Correlation Coefficient) [6] evaluates the correlation between predicted and true labels:

$$\text{MCC} = \frac{\text{TP} \cdot \text{TN} - \text{FP} \cdot \text{FN}}{\sqrt{(\text{TP} + \text{FP}) \cdot (\text{TP} + \text{FN}) \cdot (\text{TN} + \text{FP}) \cdot (\text{TN} + \text{FN})}},$$

  which returns a value between $-1$ and $1$, where $1$ indicates perfect prediction, $0$ corresponds to random prediction, and $-1$ indicates total disagreement. MCC provides a balanced and informative measure even under severe class imbalance, as it incorporates all four terms of the confusion matrix.

The Davies-Bouldin Index (DBI) [7] quantifies cluster quality by favoring clusters that are both compact within clusters and well separated between clusters. It is defined as

$$\text{DBI} = \frac{1}{k} \sum_{i=1}^{k} \text{max}(\frac{\Delta(X_i) + \Delta(X_j)}{\delta(X_i, X_j)})$$

for $k$ clusters, where $\Delta(X_i)$ measures the intra-cluster distances of cluster $i$ and $\delta(X_i, X_j)$ measures the inter-cluster distances between the centroids of clusters $i$ and $j$. Lower DBI values indicate tighter within-cluster cohesion and greater separation between clusters.

#### Supplementary Note 5: Broader adaptation to diverse datasets

To further support the broad applicability of NovAST across diverse datasets and sequencing modalities, including integration with single-cell RNA-seq references, we implement additional preprocessing procedures in the NovAST Python package. Specifically, when both datasets contain more than 5,000 genes, the top 5,000 highly variable genes (HVGs) are selected from each dataset using Scanpy's `highly_variable_genes` function with `"seurat_v3"` mode, and the intersection of the two HVG sets is used for analysis. To mitigate NovAST's sensitivity to imbalanced cell numbers, particularly when the reference dataset contained substantially more cells, as often occurs when transferring from a single-cell atlas to a spatial transcriptomics slice, an automatic subsampling step is applied. This step reduces the reference set to match the target's size while preserving cell type proportions and is triggered when the reference contains at least three-fold more cells than the target.

#### Supplementary Figures

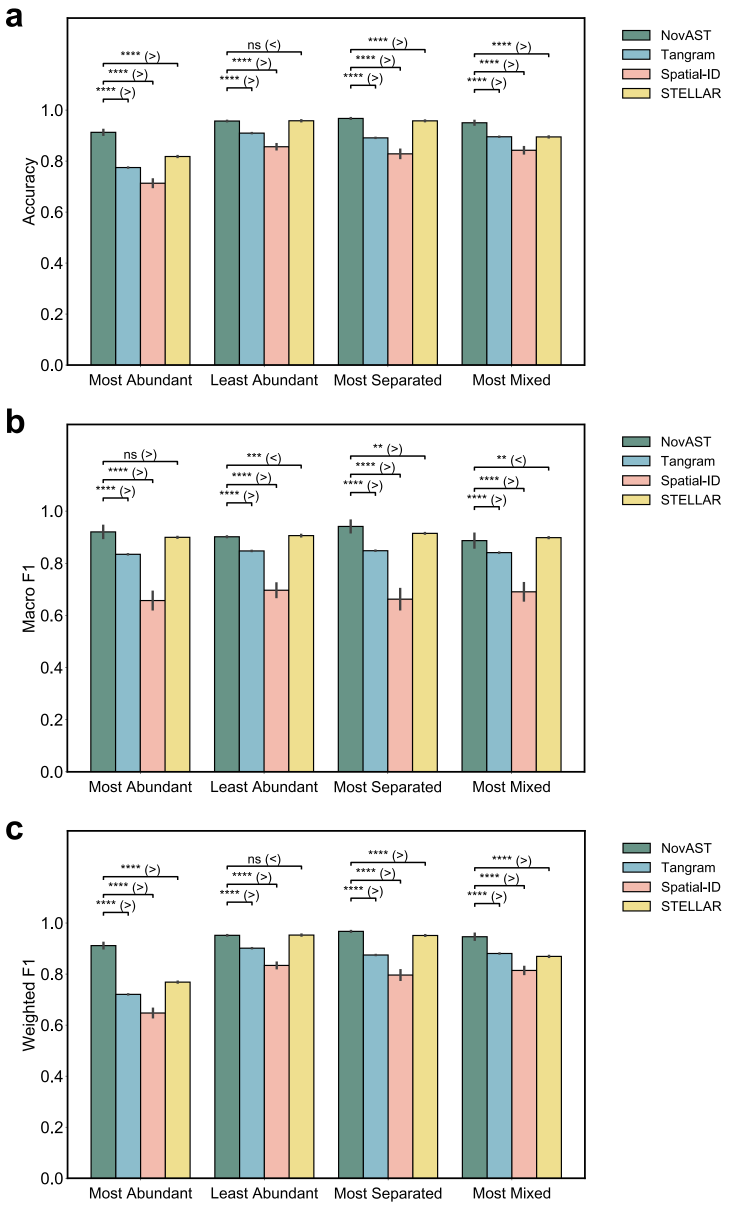

**Supplementary Figure 1**: **Accuracy and F1 scores comparisons of NovAST with benchmarking methods in the mouse MERFISH MOp dataset**

Comparison of NovAST with Tangram, Spatial-ID, and STELLAR across pseudo-novel cell type removal settings, evaluated using **(a)** Accuracy, **(b)** Macro F1, and **(c)** Weighted F1. Error bars denote the standard deviation across 10 runs. Statistical significance was assessed using a one-sided Mann–Whitney U test under the alternative hypothesis that NovAST outperforms the competing method when NovAST has a higher sample mean, and vice versa; **** indicates $p < 0.0001$, *** indicates $p < 0.001$, ** indicates $p < 0.01$, and 'ns' denotes not significant. See Methods for details on the pseudo-novel settings.

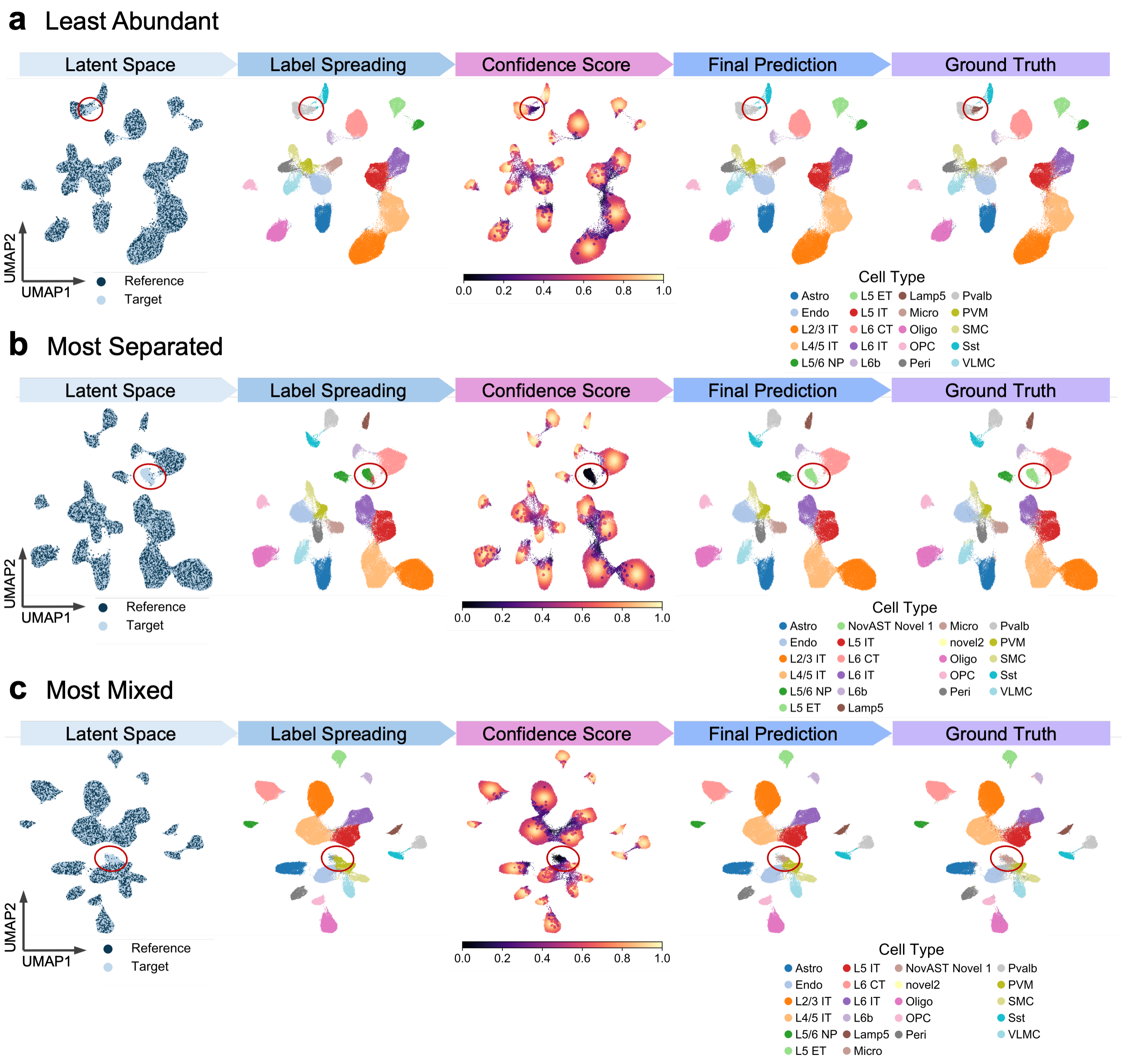

**Supplementary Figure 2**: **Representative UMAP projections of NovAST latent embeddings across pseudo-novel settings in the mouse MERFISH MOp dataset**
Representative UMAP projections of the latent embedding learned by NovAST under three additional pseudo-novel settings: **(a)** Least Abundant, **(b)** Most Separated, and **(c)** Most Mixed. For each setting, five subpanels are shown in a row, corresponding to the latent space colored by reference versus target domains (Latent Space), label propagation results (Label Spreading), confidence scores ranging from 0 to 1 (Confidence Score), the final predicted cell types (Final Prediction), and the ground truth annotations (Ground Truth). These visualizations demonstrate that NovAST consistently produces structured and informative embedding across different settings, enabling accurate label transfer and robust novel cell type detection. The red circle highlights the novel cell cluster in each subpanel.

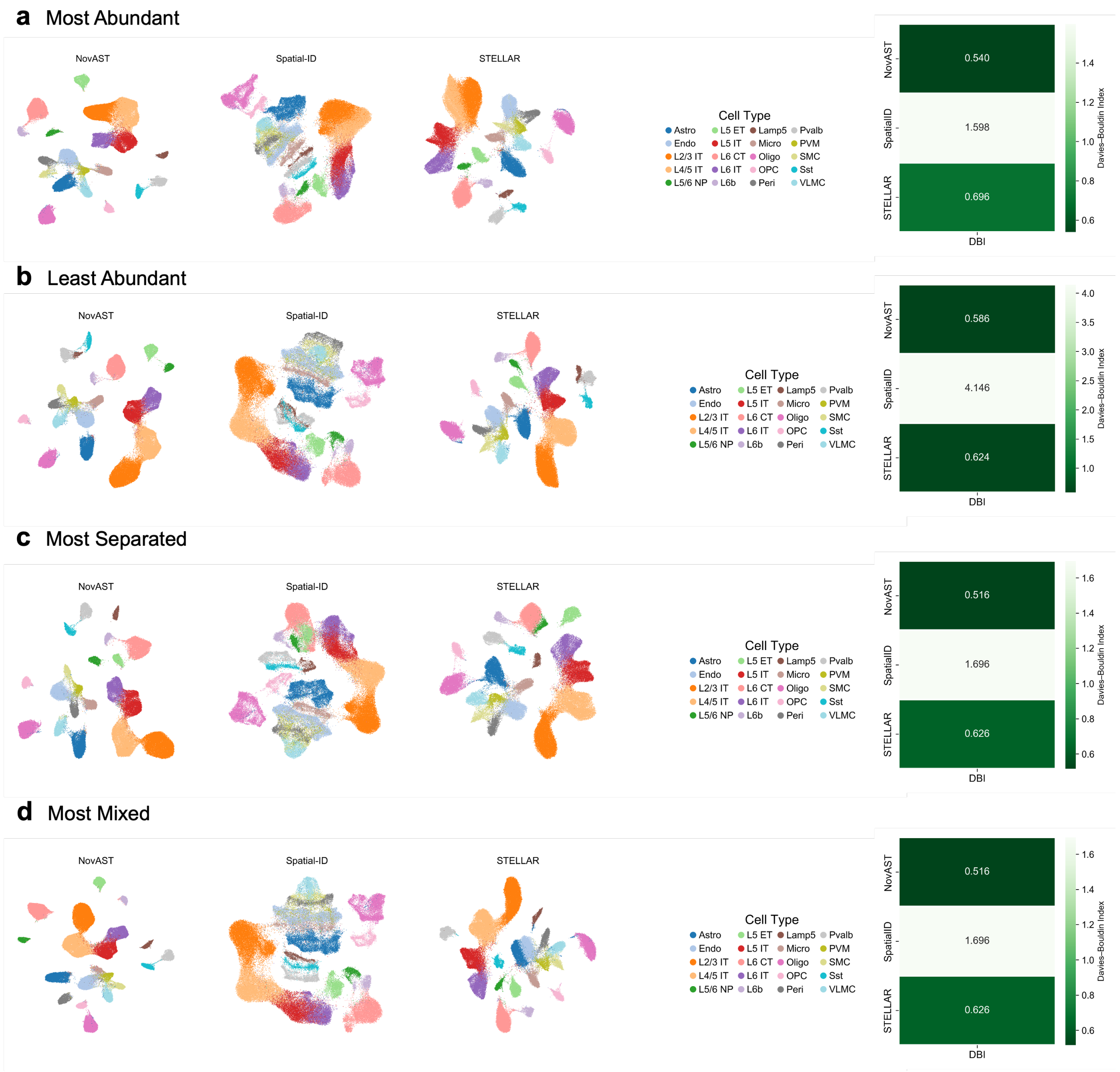

**Supplementary Figure 3**: **Representative UMAP projections of latent embeddings produced by NovAST and benchmarking methods across pseudo-novel settings in the mouse MERFISH MOp dataset**

Representative UMAP projections of the latent embedding learned by NovAST and benchmarking methods under all pseudo-novel settings: **(a)** Most Abundant, **(a)**Least Abundant, **(b)** Most Separated, and **(c)** Most Mixed. For each setting, five subpanels are shown in a row, corresponding to the latent space of NovAST, Spatial-ID, STELLAR, followed by the cell type legend and the Davies–Bouldin Index (DBI) comparison across the three methods. Tangram is not included here because it does not produce a latent embedding suitable for UMAP visualization.

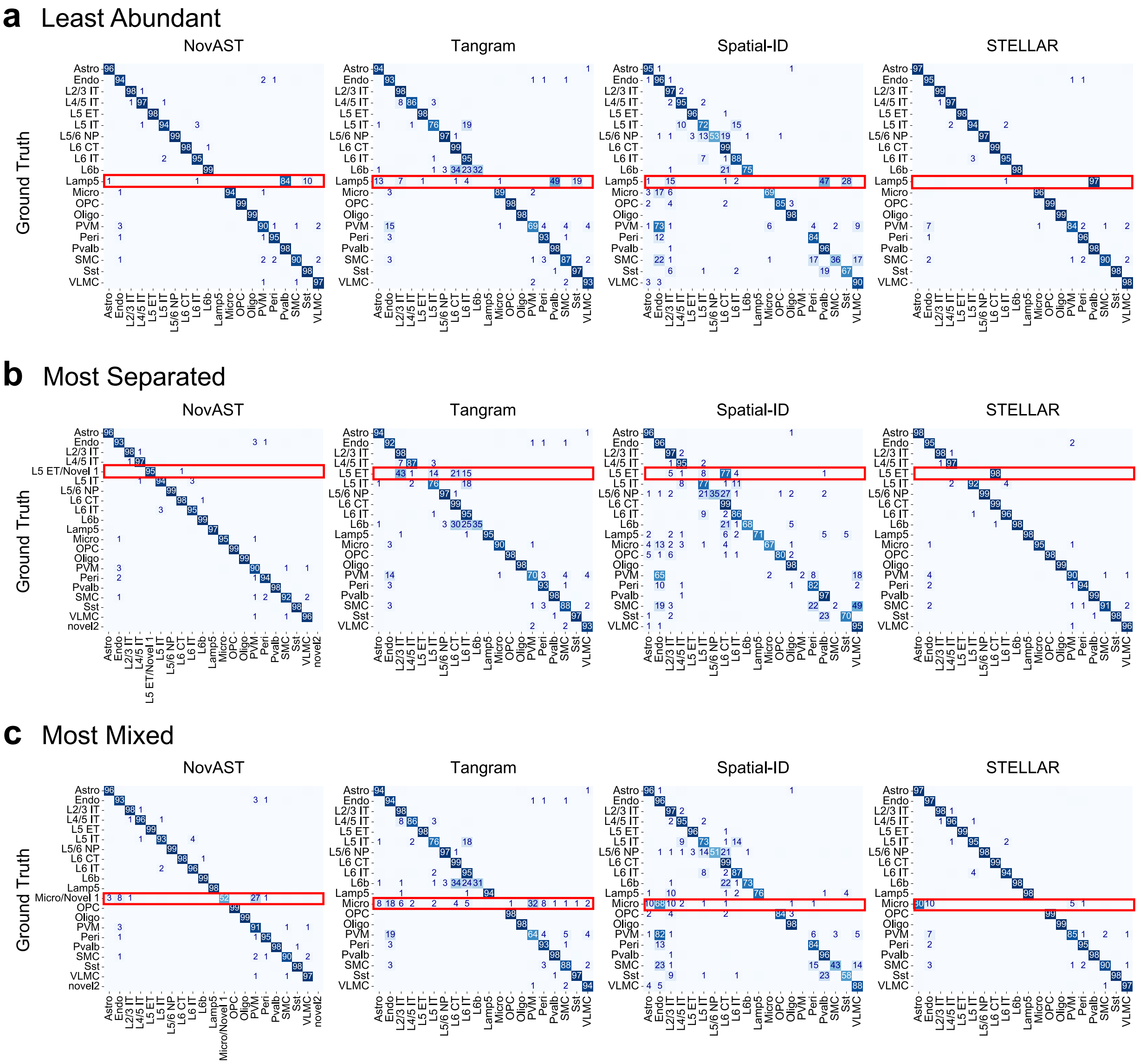

**Supplementary Figure 4**: **Confusion matrices comparing cell type predictions across all slices in the mouse MERFISH MOp dataset**

Confusion matrices compare predicted cell type labels with ground truth for NovAST and benchmarking methods. Each matrix is row-normalized by the ground truth labels, with entries indicating the percentage of cells from each true class assigned to each predicted label. Subfigures show results for the **(a)** Least Abundant, **(b)** Most Separated, and **(c)** Most Mixed settings. Predictions of the novel cluster were merged with the ground truth to account for the intentionally removed reference cell type. A red box highlights the cell type removed from the reference and its corresponding predictions for each method, which are Lamp5, L5 ET, and Micro for **(a-c)**, respectively.

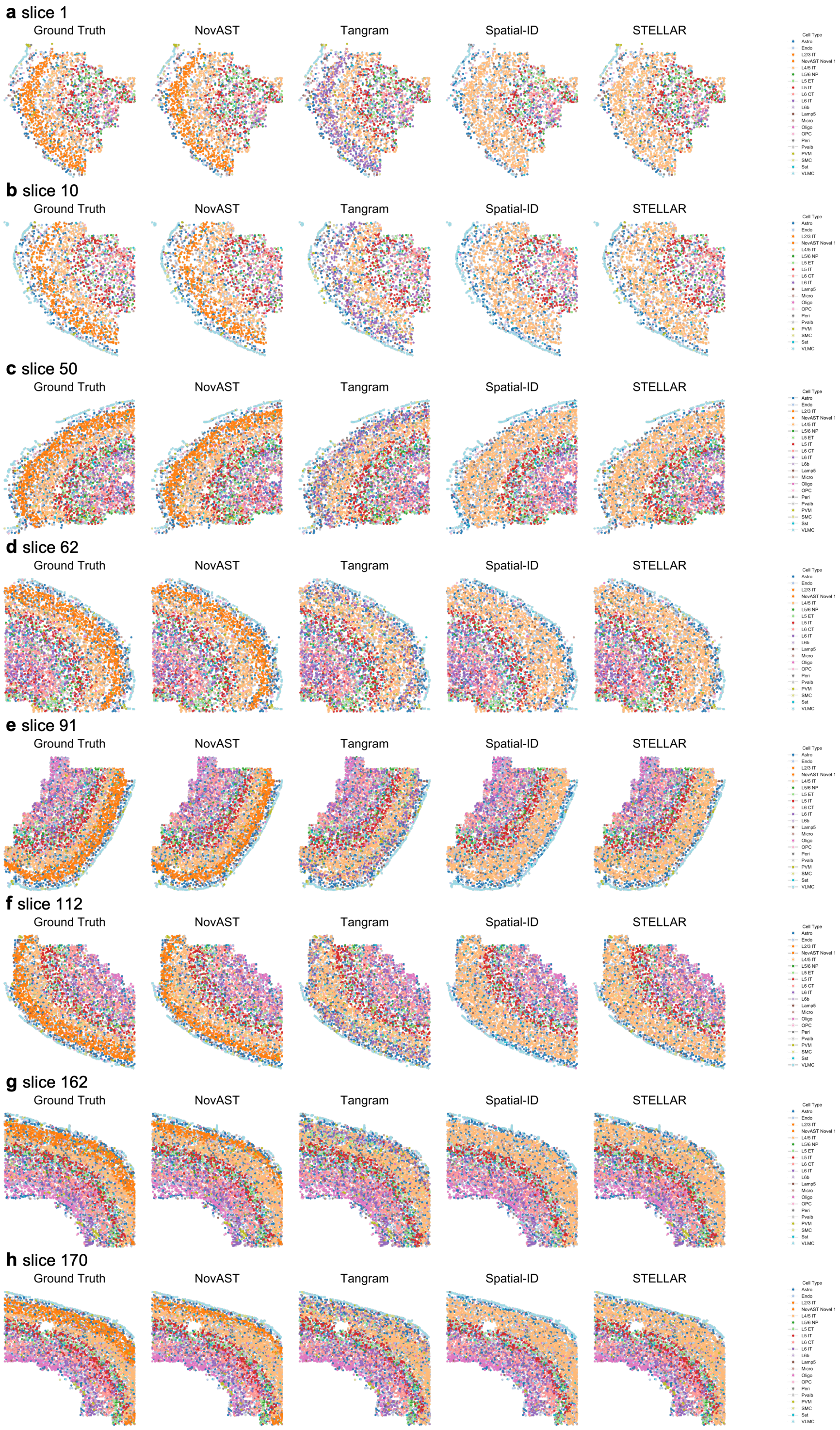

**Supplementary Figure 5**: **Spatial visualization of predicted cell types across additional tissue slices in the mouse MERFISH MOp dataset**
Spatial maps of cell type predictions from NovAST and benchmarking methods under the same setting as in Figure 2d, where the most abundant cell type was removed from the reference data. Eight slices presented here, with rows corresponding to the ground truth annotation and predictions from NovAST and competing methods.

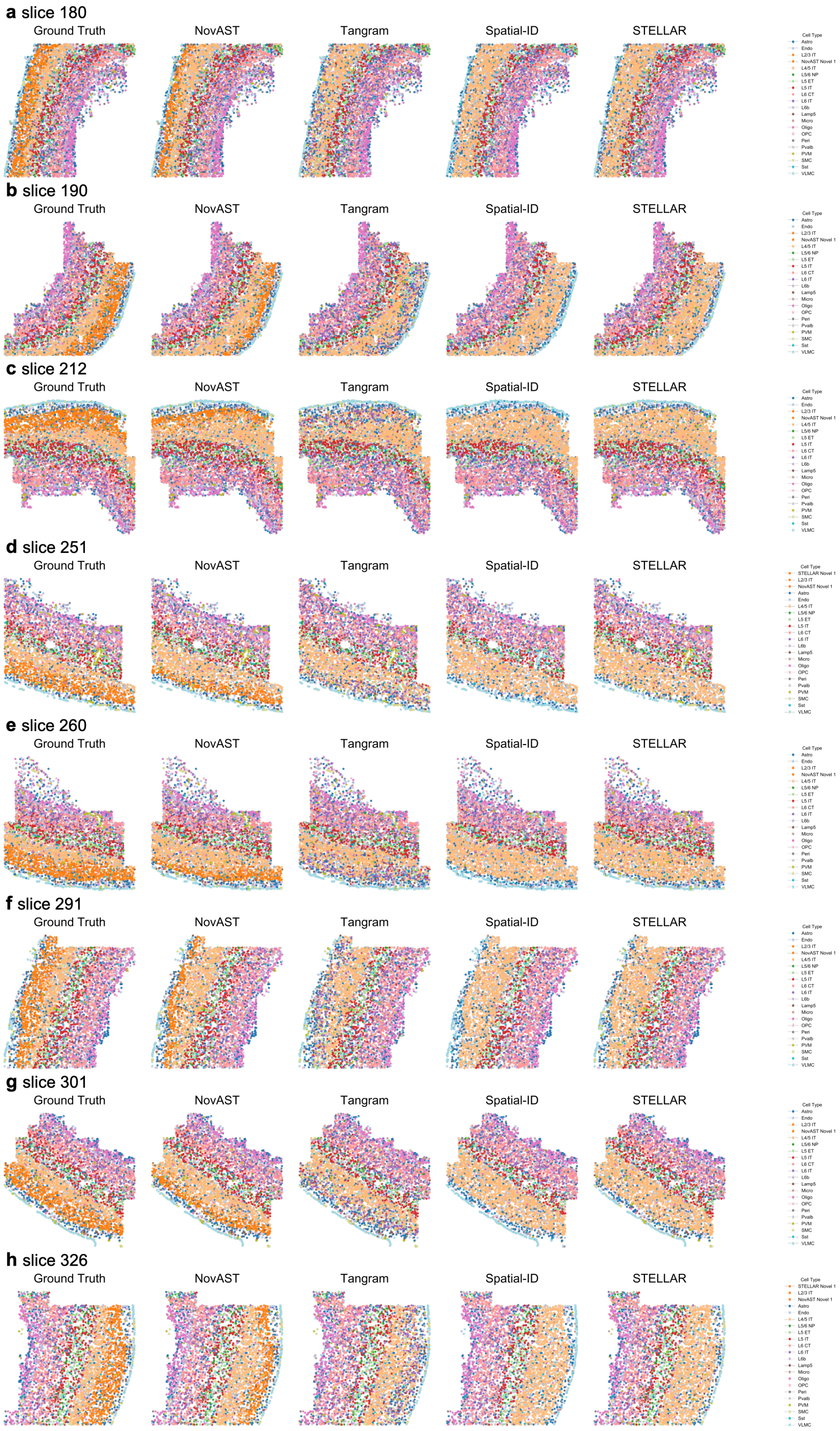

**Supplementary Figure 6: Spatial visualization of predicted cell types across additional tissue slices in the mouse MERFISH MOp dataset (continued on Supplementary Figure 5)**

Spatial maps of cell type predictions from NovAST and benchmarking methods under the same setting as in Figure 2d, where the most abundant cell type was removed from the reference data. Eight slices presented here, with rows corresponding to the ground truth annotation and predictions from NovAST and competing methods.

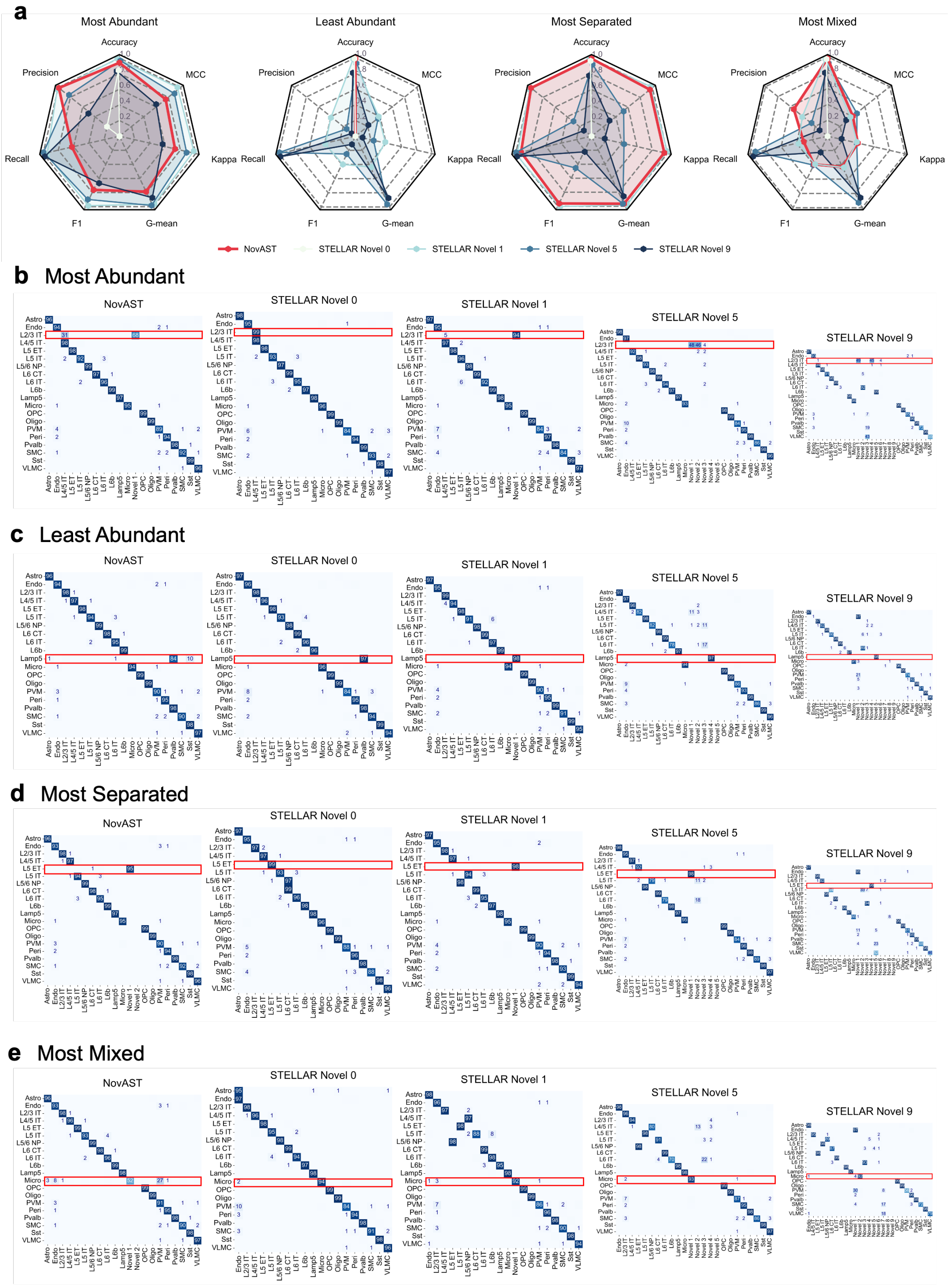

**Supplementary Figure 7: Comparative evaluation of novel cell identification between NovAST and STELLAR in the mouse MERFISH MOp dataset (caption continued next page)**

**(a)** Four radar plots with seven axes (Accuracy, Precision, Recall, F1, G-mean, Cohen's kappa, MCC) and five traces summarize the performance of NovAST and STELLAR, where STELLAR is evaluated under four settings of its novel seed parameter (expected number of novel cell types): 0 (default), 1, 5, and 9. Each panel corresponds to one of the four settings: Most Abundant, Least Abundant, Most Separated, and Most Mixed. Radial values are on a common scale ranging from 0 to 1; higher values are better. **(b)–(e)** Confusion matrices compare predicted cell type labels with ground truth for NovAST and STELLAR under novel seeds of 0, 1, 5, and 9. Each matrix is row-normalized by the ground truth labels, with entries indicating the percentage of cells from each true class assigned to each predicted label. Subfigures show results for the **(b)** Most Abundant, **(c)** Least Abundant, **(d)** Most Separated, and **(e)** Most Mixed settings. A red box highlights the cell types removed from the reference and its corresponding predictions for each method.

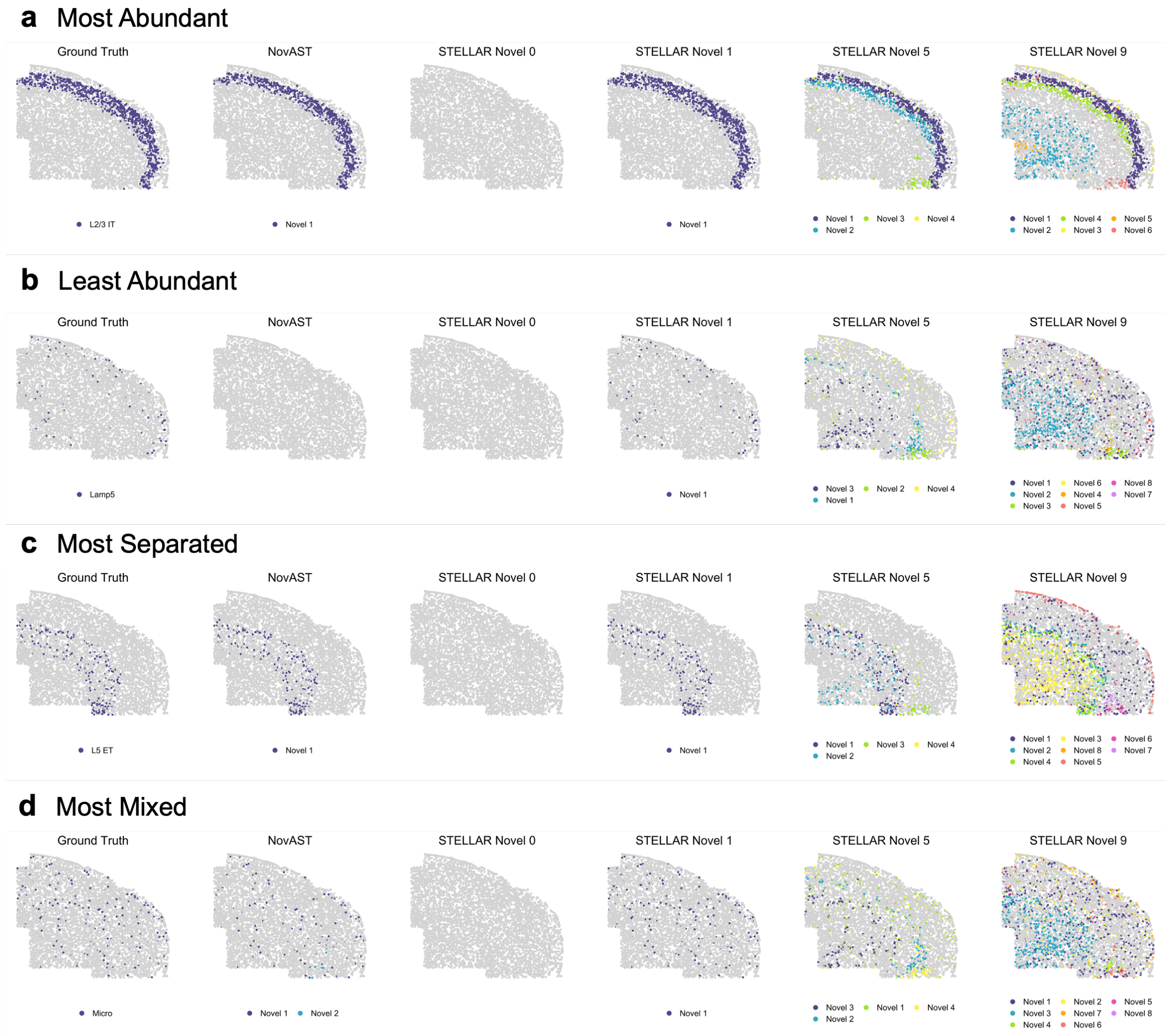

**Supplementary Figure 8**: **Representative spatial maps of novel cell identification by NovAST and STELLAR in the mouse MERFISH MOp dataset**

**(a)–(d)** Spatial maps for the four pseudo-novel scenarios: **(a)** Most Abundant, **(b)** Least Abundant, **(c)** Most Separated, and **(d)** Most Mixed. Each scenario includes six panels of the same tissue slice (slice 71), showing the ground-truth novel cells and the novel cell predictions from NovAST and STELLAR (with novel seed parameters of 0, 1, 5, and 9). Novel cells are shown in colors, and all other cells are shown in grey.

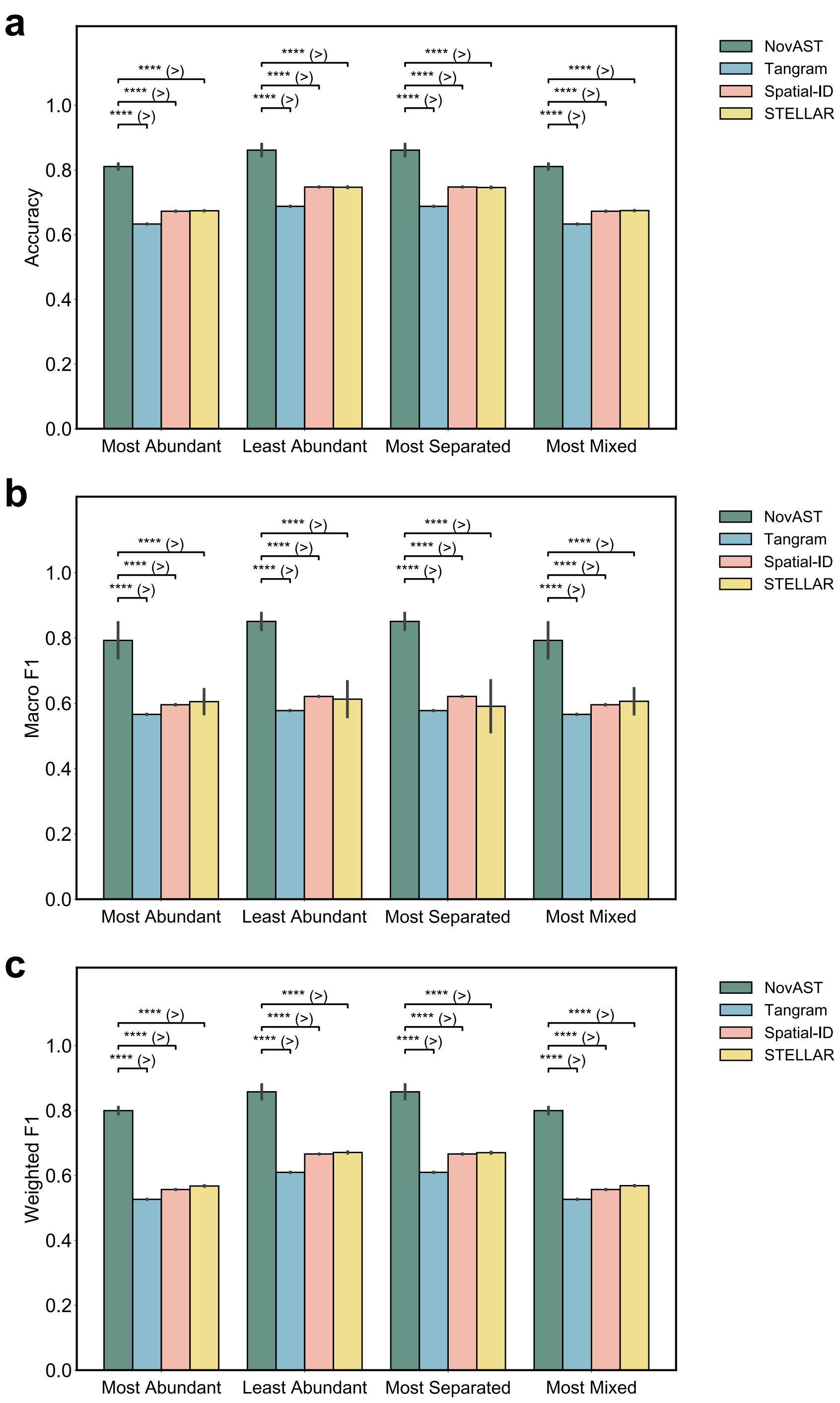

**Supplementary Figure 9**: **Accuracy and F1 scores comparisons of NovAST with benchmarking methods in the human Xenium lung**

Comparison of NovAST with Tangram, Spatial-ID, and STELLAR across pseudo-novel cell type removal settings, evaluated using **(a)** Accuracy, **(b)** Macro F1, and **(c)** Weighted F1. Error bars denote the standard deviation across 10 runs. Statistical significance was assessed using a one-sided Mann–Whitney U test under the alternative hypothesis that NovAST outperforms the competing method when NovAST has a higher sample mean, and vice versa; **** indicates $p < 0.0001$. See Methods for details on the pseudo-novel settings.

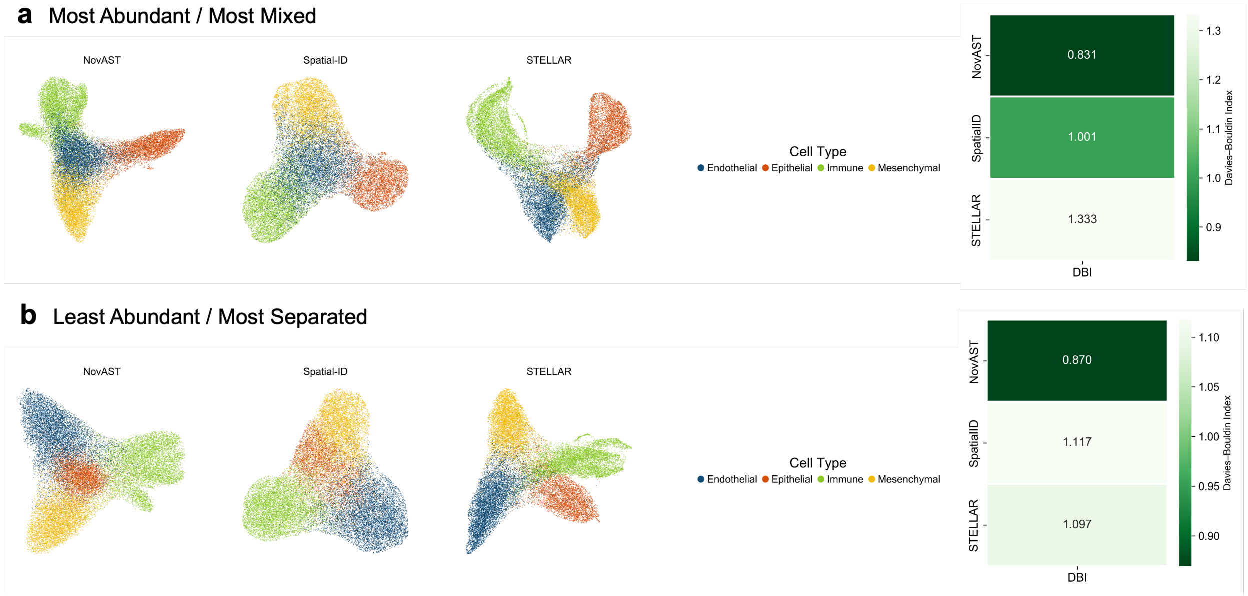

**Supplementary Figure 10**: **Representative UMAP projections of latent embeddings produced by NovAST and benchmarking methods across pseudo-novel settings in the human Xenium lung dataset**

Representative UMAP projections of the latent embedding learned by NovAST and benchmarking methods under all four pseudo-novel settings: **(a)** Most Abundant, **(a)**Least Abundant, **(b)** Most Separated, and **(c)** Most Mixed. For each setting, five subpanels are shown in a row, corresponding to the latent space of NovAST, Spatial-ID, STELLAR, followed by the cell type legend and the Davies–Bouldin Index (DBI) comparison across the three methods. Tangram is not included here because it does not produce a latent embedding suitable for UMAP visualization.

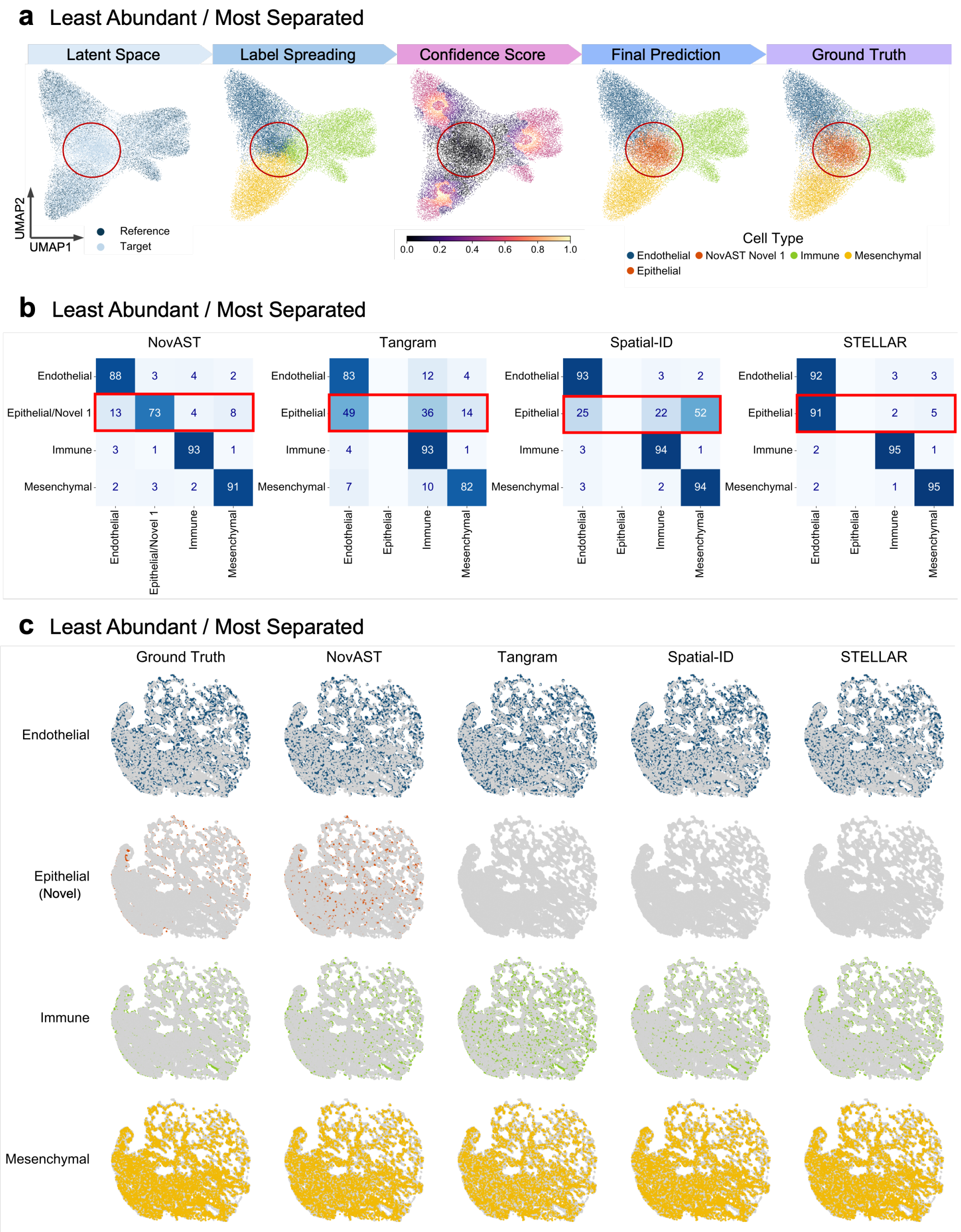

**Supplementary Figure 11: NovAST's performance in the human Xenium lung dataset under the Least Abundant / Most Separated setting.**
**(a)** UMAP projections, **(b)** confusion matrices, and **(c)** spatial visualizations are shown under the Least Abundant / Most Separated setting with epithelial cells removed from the reference data. A red circle in **(a)** highlights the novel cell cluster, and a red box in **(b)** marks the removed reference cell type. In **(b)**, Novel 1' predictions were merged with the ground truth Epithelial class for NovAST. Spatial maps in **(c)** showing individual cell types (rows) across ground truth and model predictions by NovAST, Tangram, Spatial-ID, and STELLAR (columns). Epithelial cells constitute the true novel population in this dataset. Colored dots indicate cells of the corresponding type, while all other cells are shown in gray.

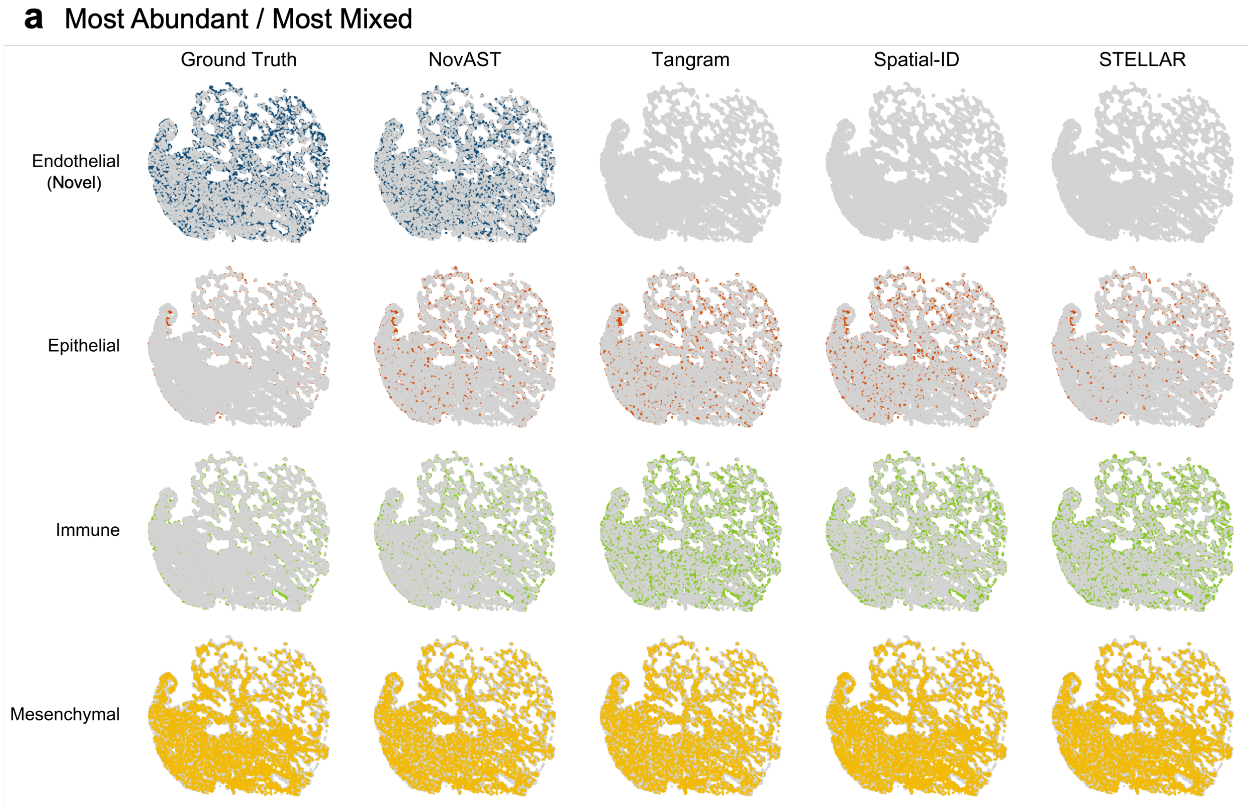

**Supplementary Figure 12**: **Spatial visualization in the human Xenium lung dataset under the Most Abundant / Most Mixed setting.**
**(a)** Spatial maps showing individual cell types (rows) across ground truth and model predictions by NovAST, Tangram, Spatial-ID, and STELLAR (columns). Endothelial cells constitute the true novel population in this dataset. Colored dots indicate cells of the corresponding type, while all other cells are shown in gray.

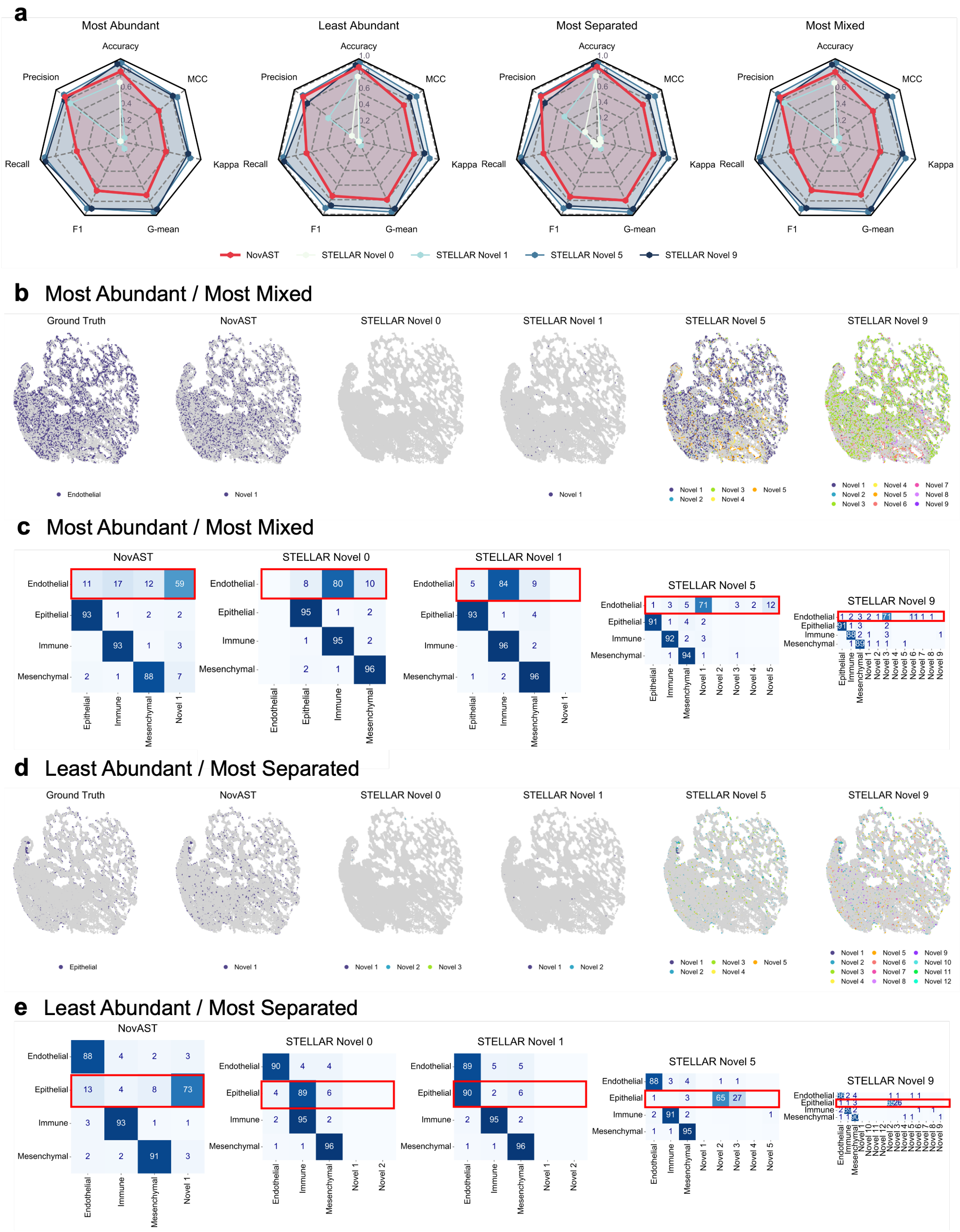

**Supplementary Figure 13: Comparative evaluation of novel cell identification between NovAST and STELLAR in the human Xenium lung dataset (caption continued next page)**

**(a)** Four radar plots with seven axes (Accuracy, Precision, Recall, F1, G-mean, Cohen's kappa, MCC) and five traces summarize the performance of NovAST and STELLAR, where STELLAR is evaluated under four settings of its novel seed parameter (expected number of novel cell types): 0 (default), 1, 5, and 9. Each panel corresponds to one of the four settings: Most Abundant, Least Abundant, Most Separated, and Most Mixed. Radial values are on a common scale ranging from 0 to 1; higher values are better. **(b-e)** Representative spatial maps and confusion matrices under the four pseudo-novel scenarios: **(b-c)** Most Abundant / Most Mixed, **(d-e)** Most Separated / Most Mixed. Representative spatial maps **(b,d)** contains six spatial panels, highlighting the ground-truth novel cells and the novel cell predictions from NovAST and STELLAR (with novel seed parameters of 0, 1, 5, and 9). Novel cells are shown in colors, and all other cells are shown in grey. Confusion matrices **(c,e)** comparing predicted and ground-truth cell-type labels for NovAST and STELLAR (with novel seed parameters of 0, 1, 5, and 9). Each matrix is row-normalized by the ground truth labels, with entries indicating the percentage of cells from each true class assigned to each predicted label. Red boxes highlight the cell types removed from the reference and its corresponding predictions for each method.

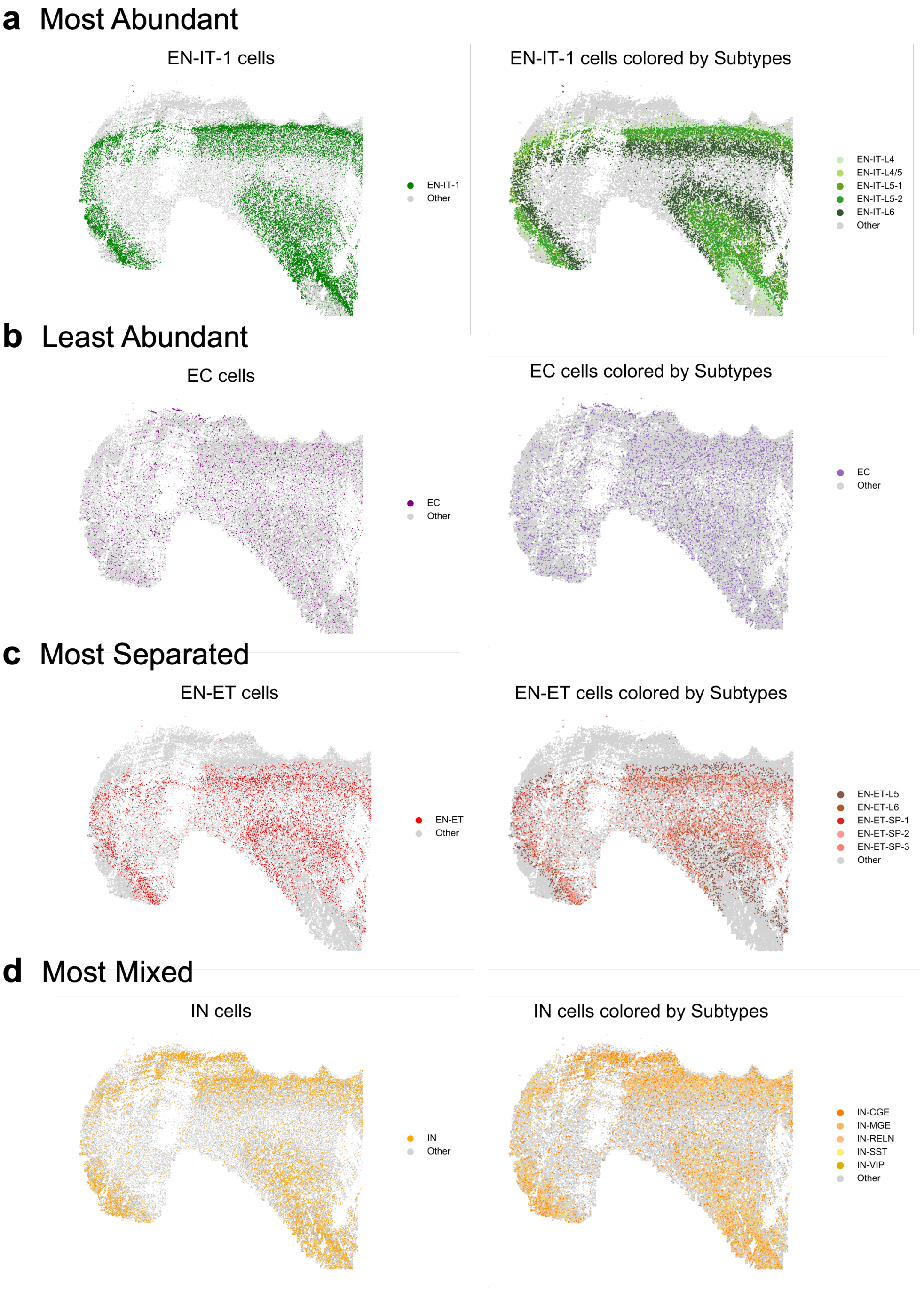

**Supplementary Figure 14**: **Spatial visualizations of pseudo-novel cell types across different settings in the human MERFISH developmental cortex dataset**
Spatial distributions of the cell types removed from the reference dataset and treated as novel in the target dataset under the **(a)** Most Abundant, **(b)** Least Abundant, **(c)** Most Separated, and **(d)** Most Mixed settings. For each subpanel, the left plot shows coarse-level labels and the right plot shows fine-grained labels, where the removed cell types are highlighted in color and all others are shown in grey. In each case, one coarse-level cell type is removed, which corresponds to one or more fine-grained subtypes.

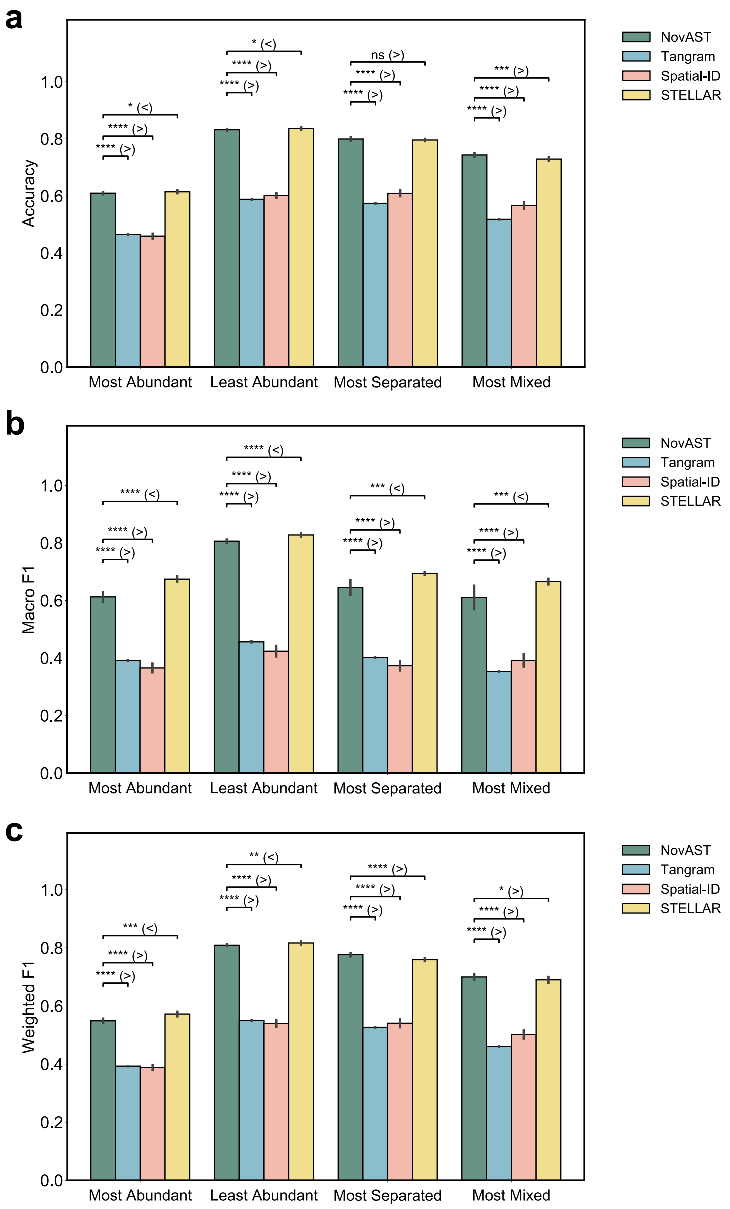

**Supplementary Figure 15**: **Accuracy and F1 scores comparisons of NovAST with benchmarking methods in the human MERFISH developmental cortex dataset**
Comparison of NovAST with Tangram, Spatial-ID, and STELLAR across pseudo-novel cell type removal settings, evaluated using **(a)** Accuracy, **(b)** Macro F1, and **(c)** Weighted F1. Error bars denote the standard deviation across 10 runs. Statistical significance was assessed using a one-sided Mann–Whitney U test under the alternative hypothesis that NovAST outperforms the competing method when NovAST has a higher sample mean, and vice versa; **** indicates $p < 0.0001$, *** indicates $p < 0.001$, ** indicates $p < 0.01$, * indicates $p < 0.05$, and 'ns' denotes not significant. See Methods for details on the pseudo-novel settings.

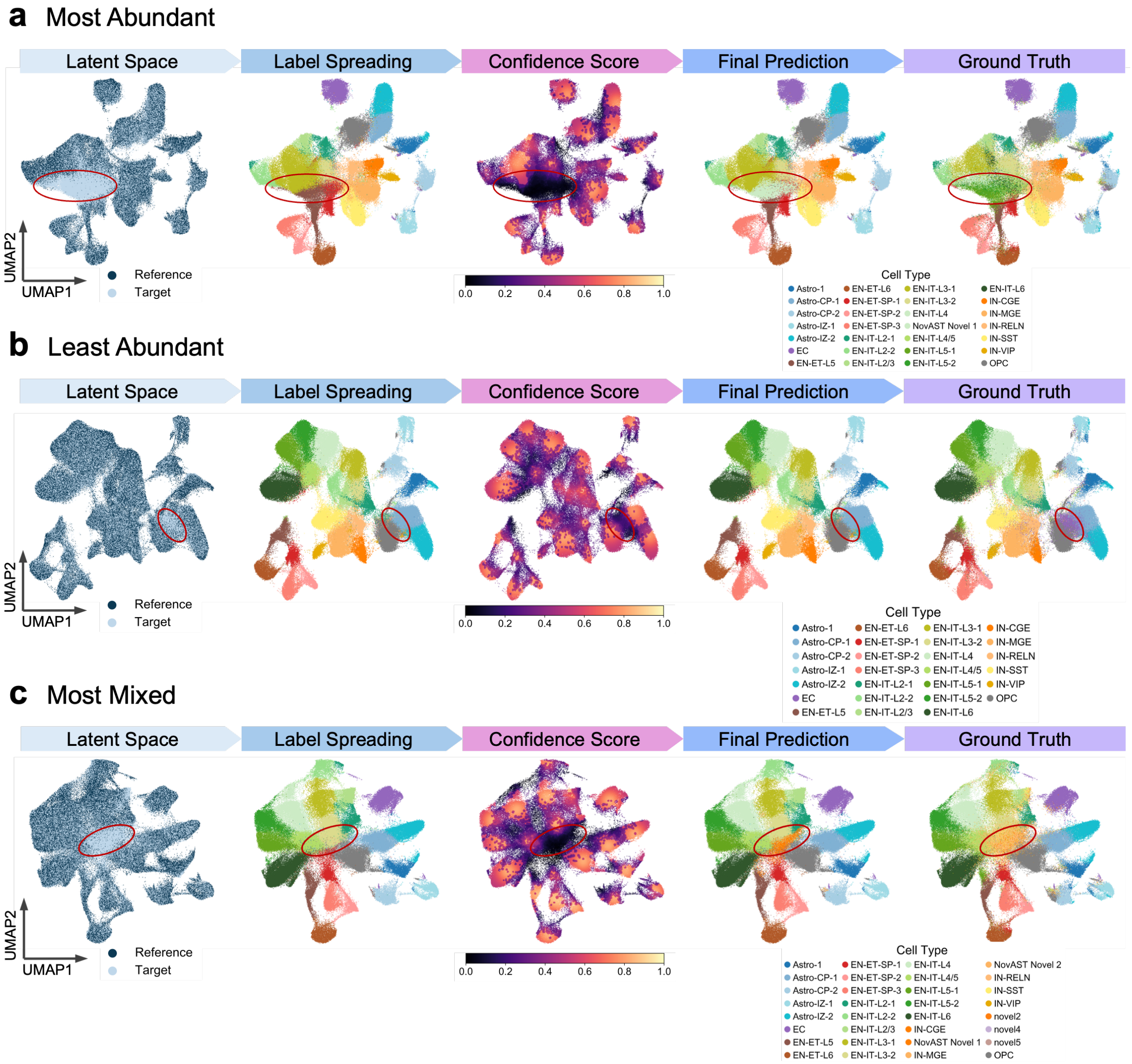

**Supplementary Figure 16**: **Representative UMAP projections of NovAST latent embeddings across pseudo-novel settings in the human MERFISH developmental cortex dataset**

Representative UMAP projections of the latent embedding learned by NovAST under three additional pseudo-novel settings: **(a)** Most Abundant, **(b)** Least Abundant, and **(c)** Most Mixed. For each setting, five subpanels are shown in a row, corresponding to the latent space colored by reference versus target domains (Latent Space), label propagation results (Label Spreading), confidence scores ranging from 0 to 1 (Confidence Score), the final predicted cell types (Final Prediction), and the ground truth annotations (Ground Truth). These visualizations demonstrate that NovAST consistently produces structured and informative embeddings across different settings, enabling accurate label transfer and robust novel cell type detection. The red circle highlights the novel cell cluster in each subpanel.

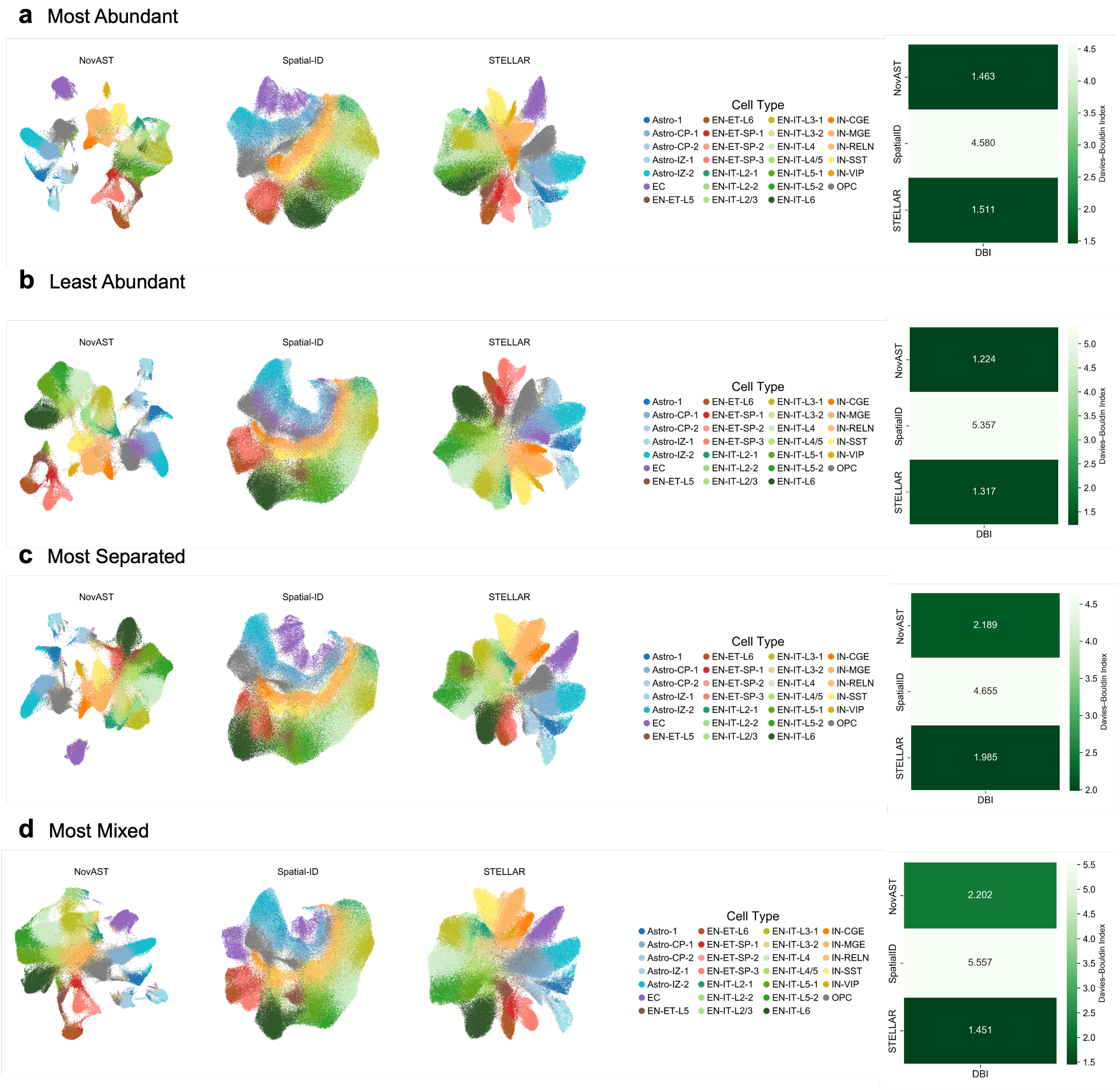

**Supplementary Figure 17**: **Representative UMAP projections of latent embeddings produced by NovAST and benchmarking methods across pseudo-novel settings in the human MERFISH developmental cortex dataset**

Representative UMAP projections of the latent embedding learned by NovAST and benchmarking methods under all pseudo-novel settings: **(a)** Most Abundant, **(a)**Least Abundant, **(b)** Most Separated, and **(c)** Most Mixed. For each setting, five subpanels are shown in a row, corresponding to the latent space of NovAST, Spatial-ID, STELLAR, followed by the cell type legend and the Davies–Bouldin Index (DBI) comparison across the three methods. Tangram is not included here because it does not produce a latent embedding suitable for UMAP visualization.

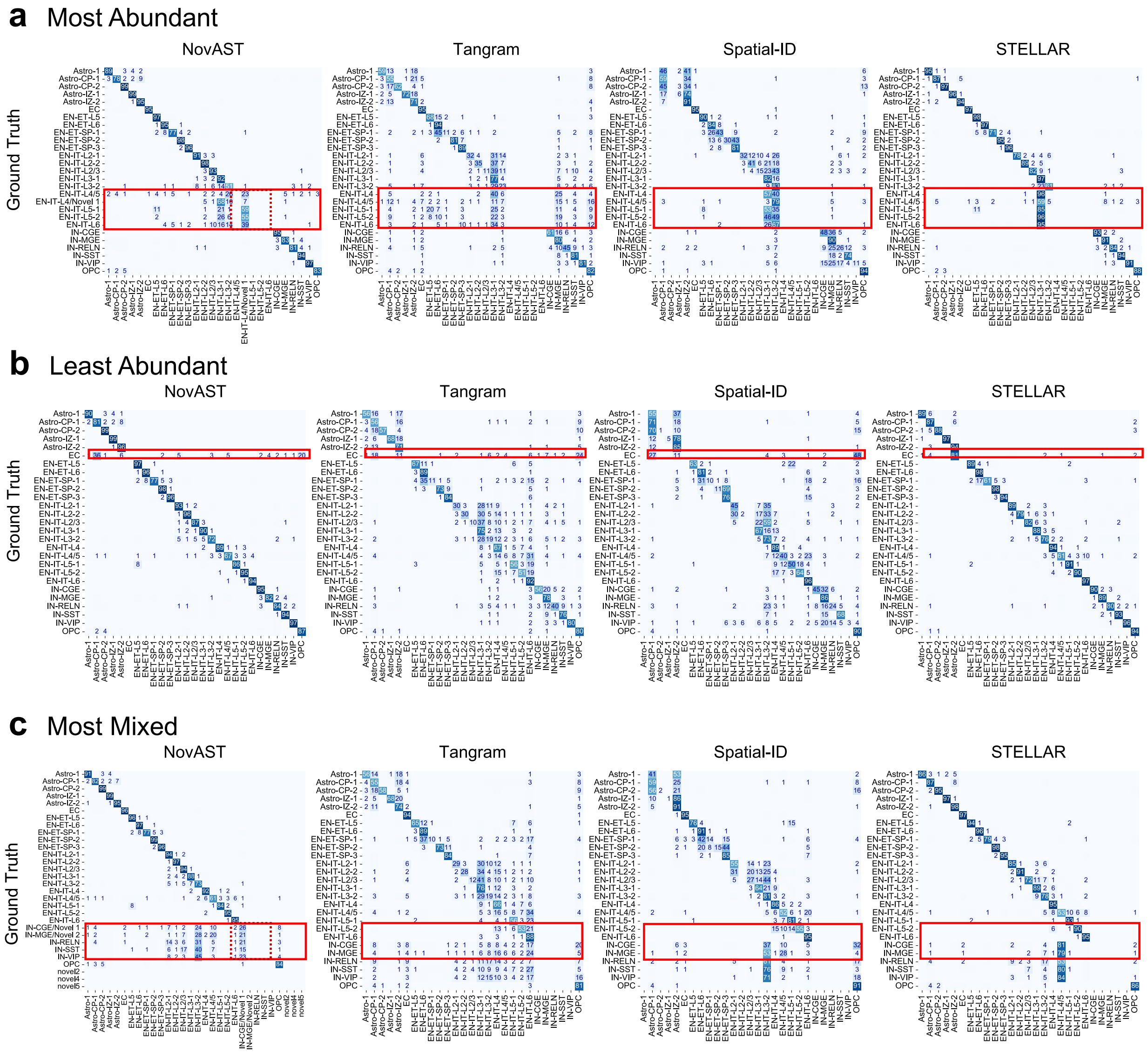

**Supplementary Figure 18: Confusion matrices comparing cell type predictions across all slices in the human MERFISH developmental cortex dataset**

Confusion matrices compare predicted cell type labels with ground truth for NovAST and benchmarking methods. Each matrix is row-normalized by the ground truth labels, with entries indicating the percentage of cells from each true class assigned to each predicted label. Subfigures show results for the **(a)** Most Abundant, **(b)** Least Abundant, and **(c)** Most Mixed settings. Predictions of the novel cluster were merged with the ground truth to account for the intentionally removed reference cell type. A red box highlights the cell types removed from the reference and its corresponding predictions for each method, which are EN-IT-1, EC, and IN at the coarse level for **(a-c)**, respectively.

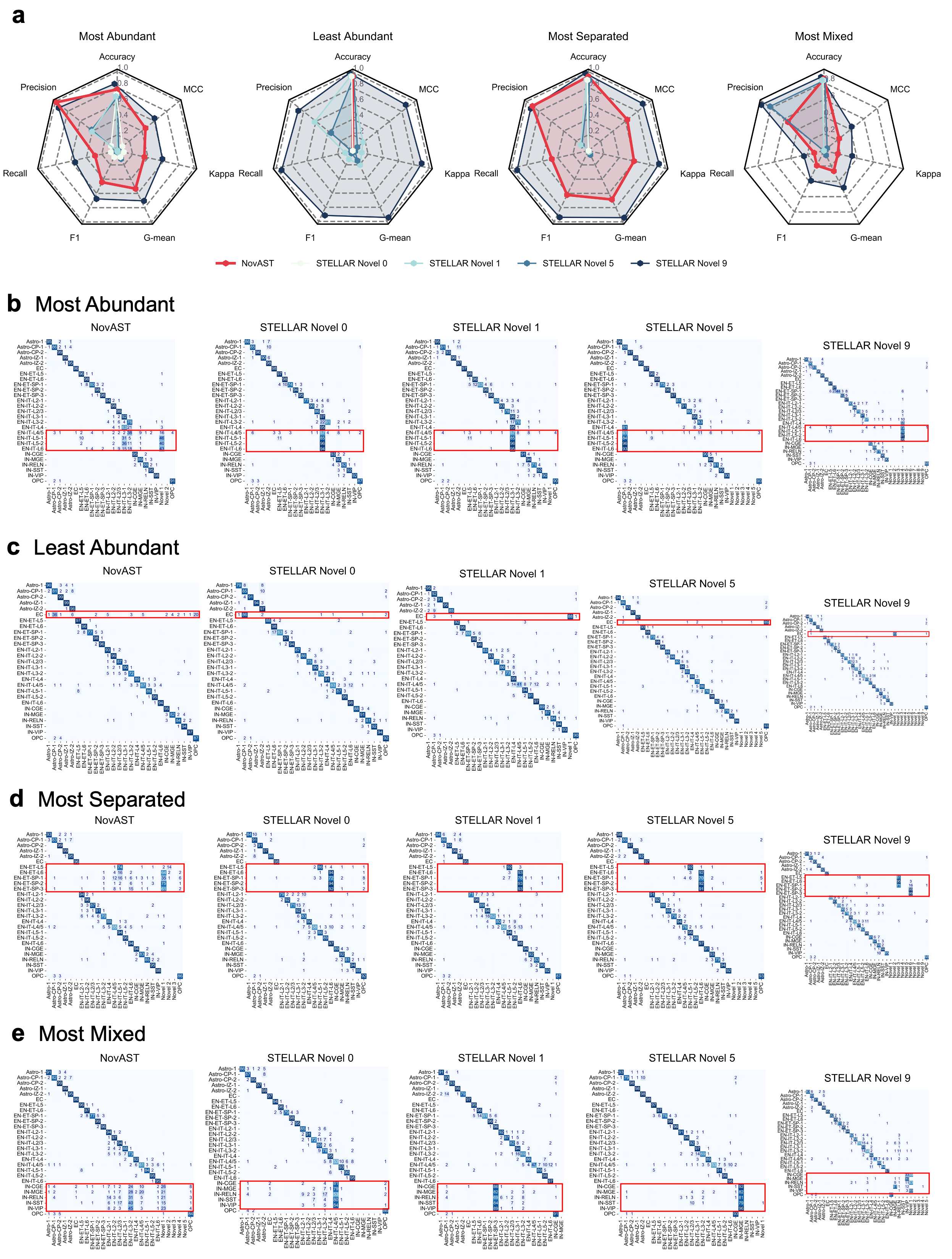

**Supplementary Figure 19**: **Comparative evaluation of novel cell identification between NovAST and STELLAR in the human MERFISH developmental cortex dataset (caption continued next page)**

**(a)** Four radar plots with seven axes (Accuracy, Precision, Recall, F1, G-mean, Cohen's kappa, MCC) and five traces summarize the performance of NovAST and STELLAR, where STELLAR is evaluated under four settings of its novel seed parameter (expected number of novel cell types): 0 (default), 1, 5, and 9. Each panel corresponds to one of the four settings: Most Abundant, Least Abundant, Most Separated, and Most Mixed. Radial values are on a common scale ranging from 0 to 1; higher values are better. **(b)–(e)** Confusion matrices compare predicted cell type labels with ground truth for NovAST and STELLAR under novel seeds of 0, 1, 5, and 9. Each matrix is row-normalized by the ground truth labels, with entries indicating the percentage of cells from each true class assigned to each predicted label. Subfigures show results for the **(b)** Most Abundant, **(c)** Least Abundant, **(d)** Most Separated, and **(e)** Most Mixed settings. A red box highlights the cell types removed from the reference and its corresponding predictions for each method.

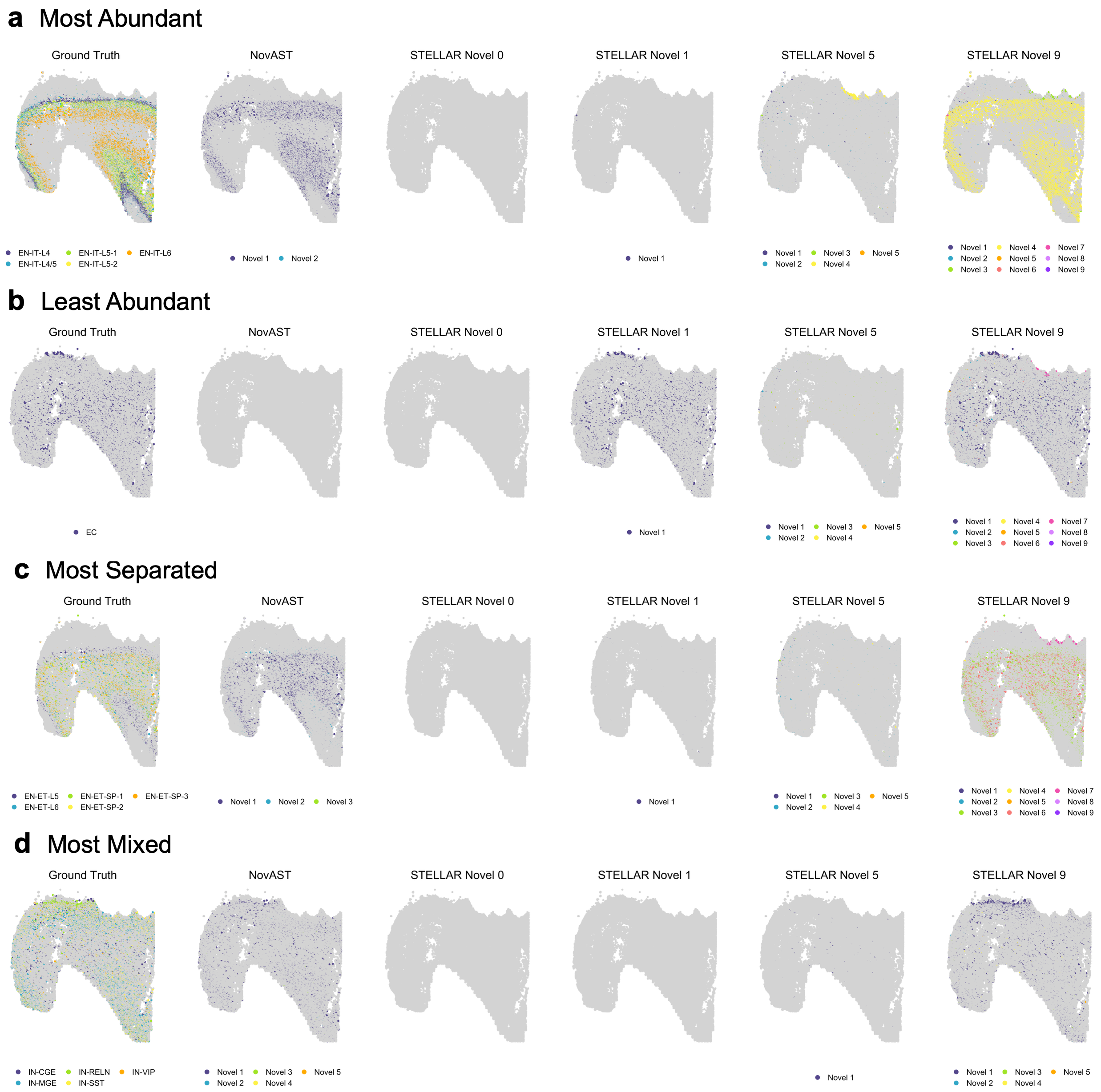

**Supplementary Figure 20: Representative spatial maps of novel cell identification by NovAST and STELLAR in the human MERFISH developmental cortex dataset** **(a)–(d)** Spatial maps for the four pseudo-novel scenarios: **(a)** Most Abundant, **(b)** Least Abundant, **(c)** Most Separated, and **(d)** Most Mixed. Each scenario includes six panels of the annotated tissue slice, showing the ground-truth novel cells and the novel cell predictions from NovAST and STELLAR (with novel seed parameters of 0, 1, 5, and 9). Novel cells are shown in colors, and all other cells are shown in grey.

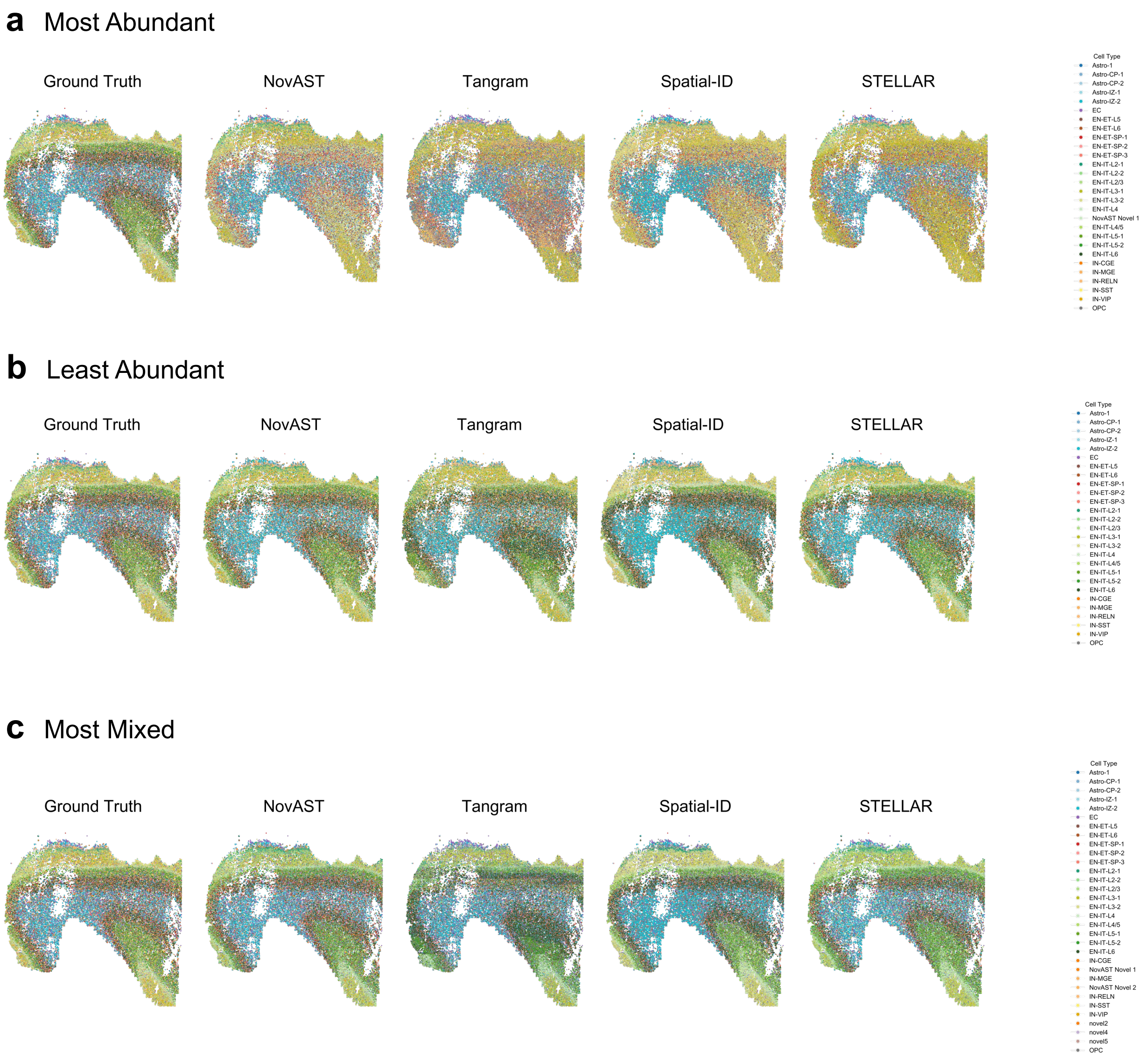

**Supplementary Figure 21**: **Spatial visualization of predicted cell types across additional tissue slices in the human MERFISH developmental cortex dataset**
Spatial maps of ground truth annotation and predictions from NovAST and benchmarking methods under the **(a)** Most Abundant, **(b)** Least Abundant, and **(c)** Most Mixed settings. The Most Separated setting is shown in Figure 4e.

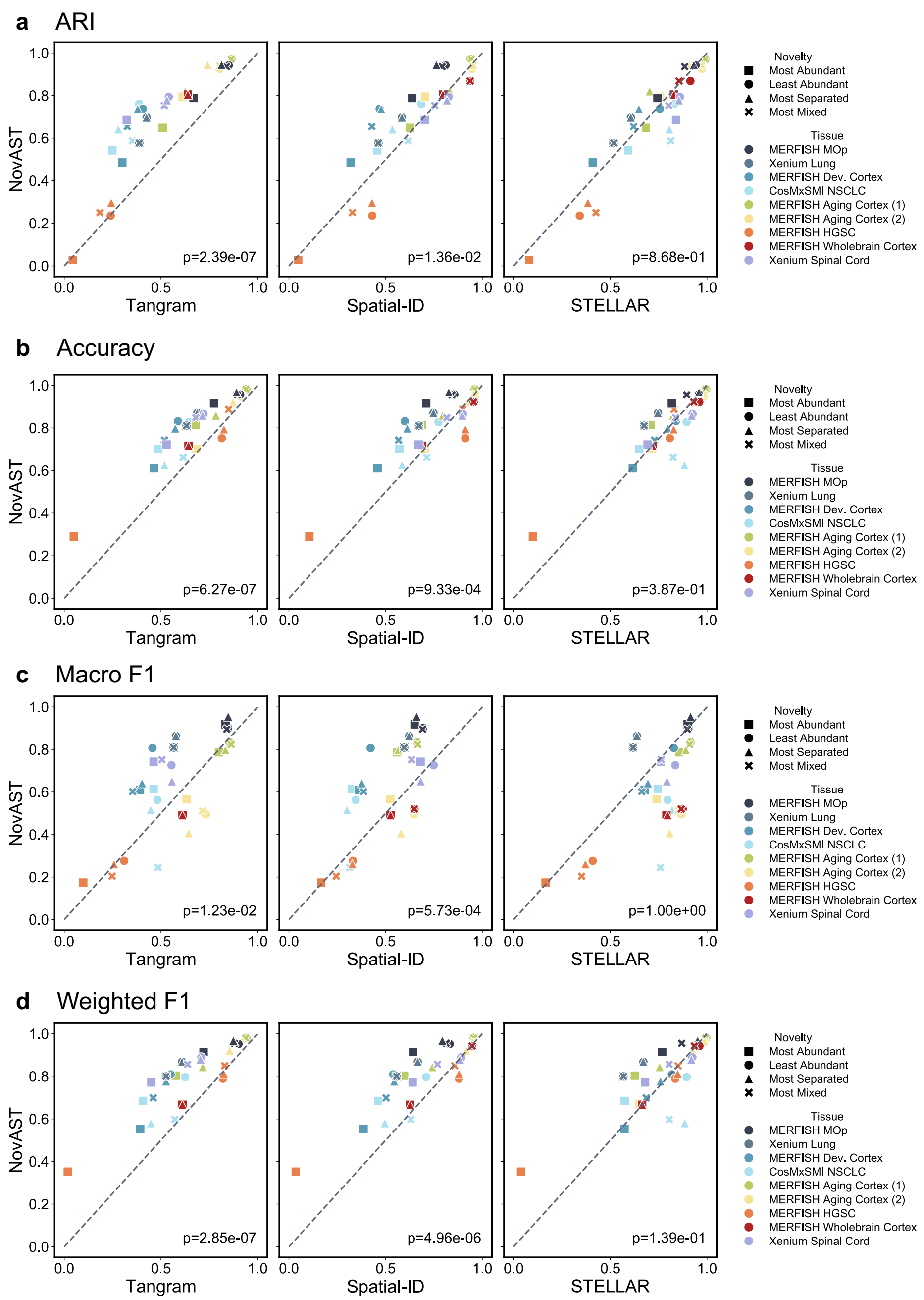

**Supplementary Figure 22**: **NovAST's overall performance across pseudo-novel settings on all datasets**

Subpanels **(a–d)** correspond to four evaluation metrics: **(a)** ARI, **(b)** Accuracy, **(c)** Macro F1, and **(d)** Weighted F1. Each subpanel contains three scatter plots, where the x-axis shows the performance of competing methods and the y-axis shows NovAST performance. Each dot represents one of nine datasets and one of four pseudo-novel settings. Reported *P* values are from one-sided Wilcoxon signed-rank tests, assessing whether NovAST significantly outperforms competing methods.

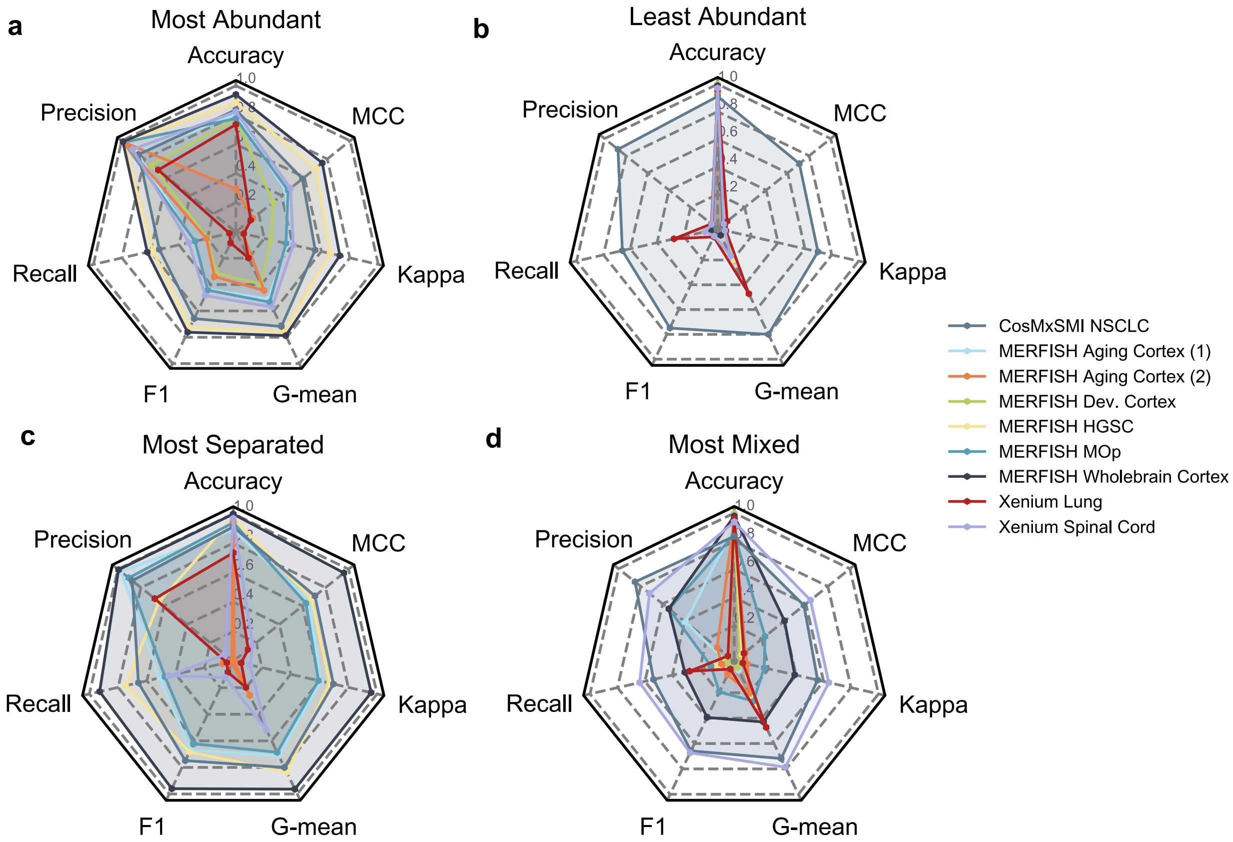

**Supplementary Figure 23**: **NovAST's novel cell identification performance across pseudo-novel settings on all datasets**

**(a)–(d)** Radar plots for the four pseudo-novel scenarios: **(a)** Most Abundant, **(b)** Least Abundant, **(c)** Most Separated, and **(d)** Most Mixed. Each radar plot summarizes performance across seven metrics (Accuracy, Precision, Recall, F1, G-mean, Cohen's kappa, and MCC). Within each panel, each trace corresponds to one of the six datasets. Radial values are scaled from 0 to 1, with higher values indicating better performance.

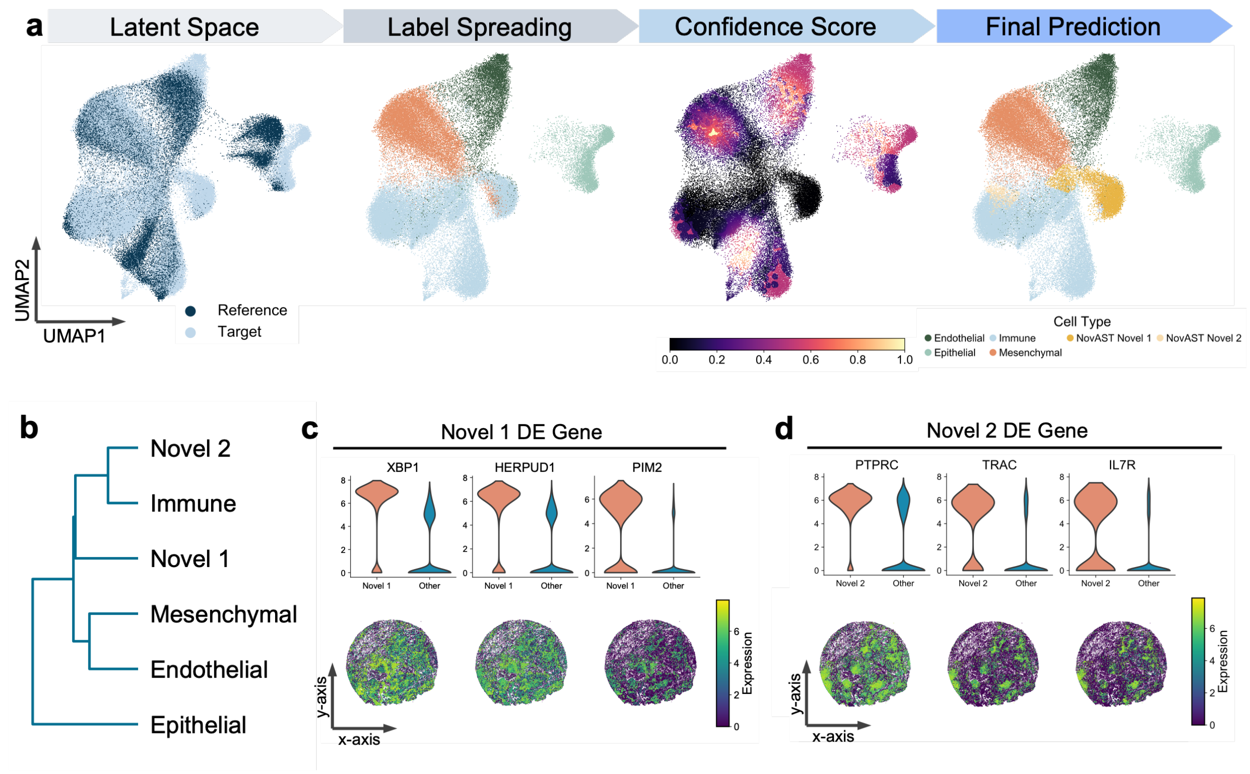

**Supplementary Figure 24**: **Representative UMAP projections of NovAST latent embeddings in the human Xenium IPF dataset**

**(a)** UMAP projections of the latent embedding learned by NovAST when applied to the human Xenium lung dataset, using a healthy donor slice as reference and an IPF patient slice as target. Results from one representative run are shown. Four subpanels are arranged in a row, corresponding to the latent space colored by reference versus target domains (Latent Space), label propagation results (Label Spreading), confidence scores ranging from 0 to 1 (Confidence Score), and the final predicted cell types (Final Prediction). **(b)** Dendrogram of predicted cell types constructed from log-normalized gene expression. **(c-d)** Expression patterns of top DE genes for Novel 1 **(c)** and Novel 2 **(d)** cluster. Violin plots (top) compare expression of selected DE genes in Novel cells versus all other cells, while spatial maps (bottom) show expression patterns for six representative DE genes.

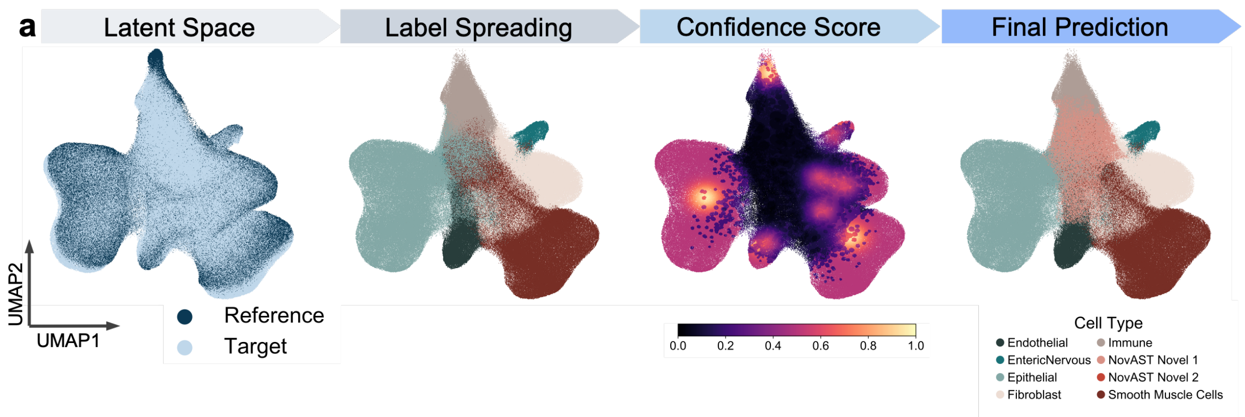

**Supplementary Figure 25**: **Representative UMAP projections of NovAST latent embeddings in the mouse MERFISH inflamed colon dataset**
UMAP projections of the latent embedding learned by NovAST when applied to the MERFISH mouse colon dataset, using slices from donors before DSS treatment as the reference and slices at peak inflammation as the target. Results from one representative run are shown. Four subpanels are arranged in a row, corresponding to the latent space colored by reference versus target domains (Latent Space), label propagation results (Label Spreading), confidence scores ranging from 0 to 1 (Confidence Score), and the final predicted cell types (Final Prediction).

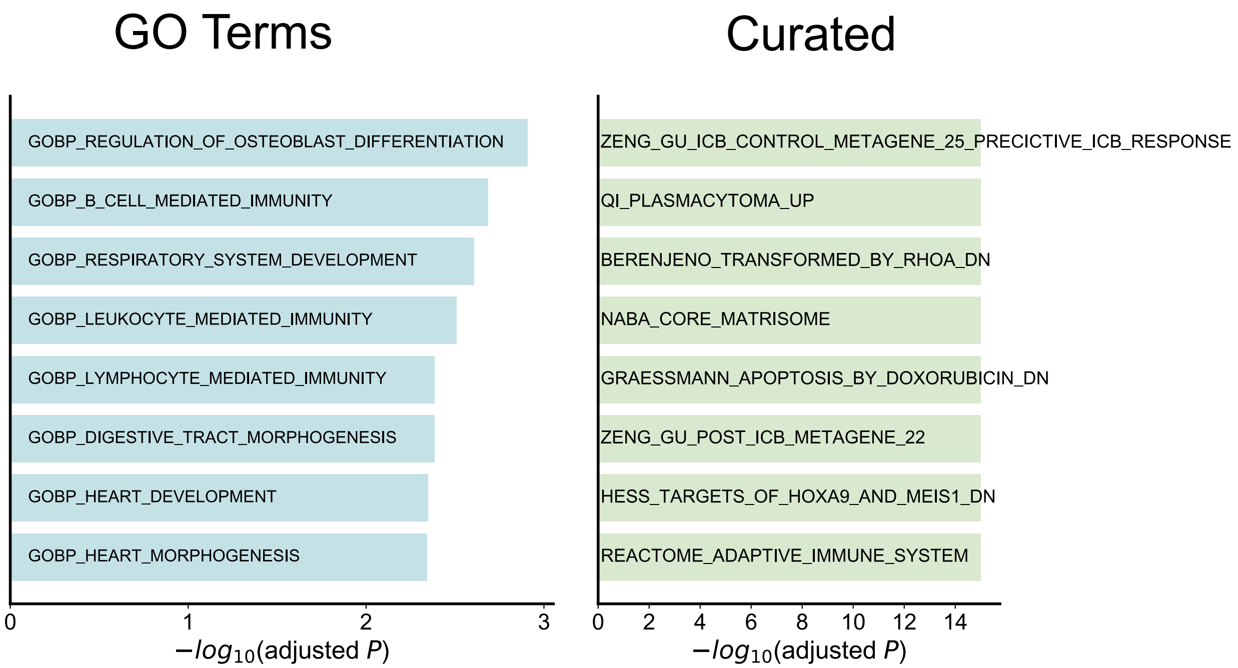

**Supplementary Figure 26**: **Gene set enrichment analysis of DE genes from the Novel 1 cluster in the mouse MERFISH inflamed colon dataset**
Gene set enrichment analysis of DE genes from the Novel 1 cluster. The left panel shows enrichment against GO terms, and the right panel shows enrichment against curated datasets from the MSigDB database. Bars show the top-ranked gene sets ordered by –$\log_{10}$(adjusted *P*), with gene set names labeled on the bars.

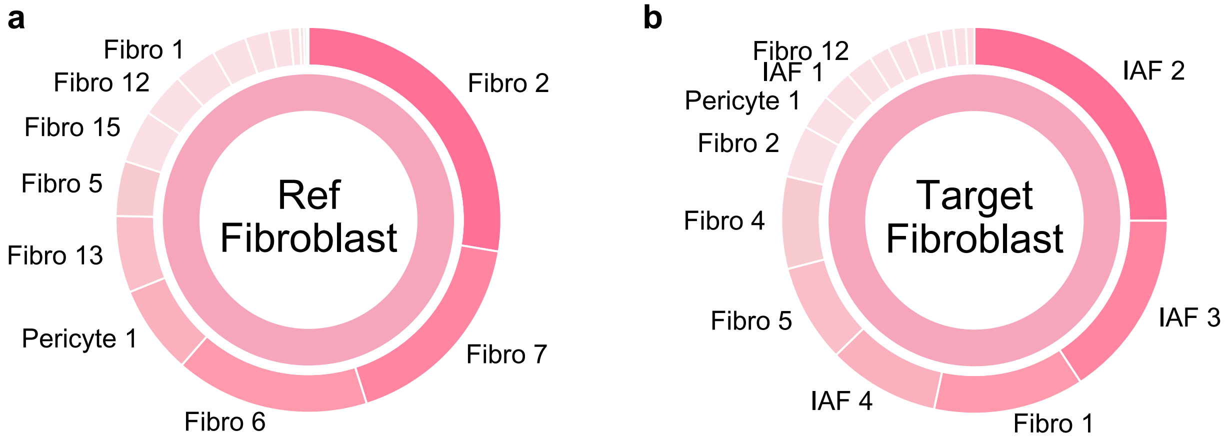

**Supplementary Figure 27**: **Proportional differences between reference and target datasets for immune and fibroblast cells in the mouse MERFISH inflamed colon dataset**
Donut plots with two layers showing subtype proportions at the outer ring for fibroblasts in the (**a**) reference and (**b**) target. The most abundant subtypes are labeled.

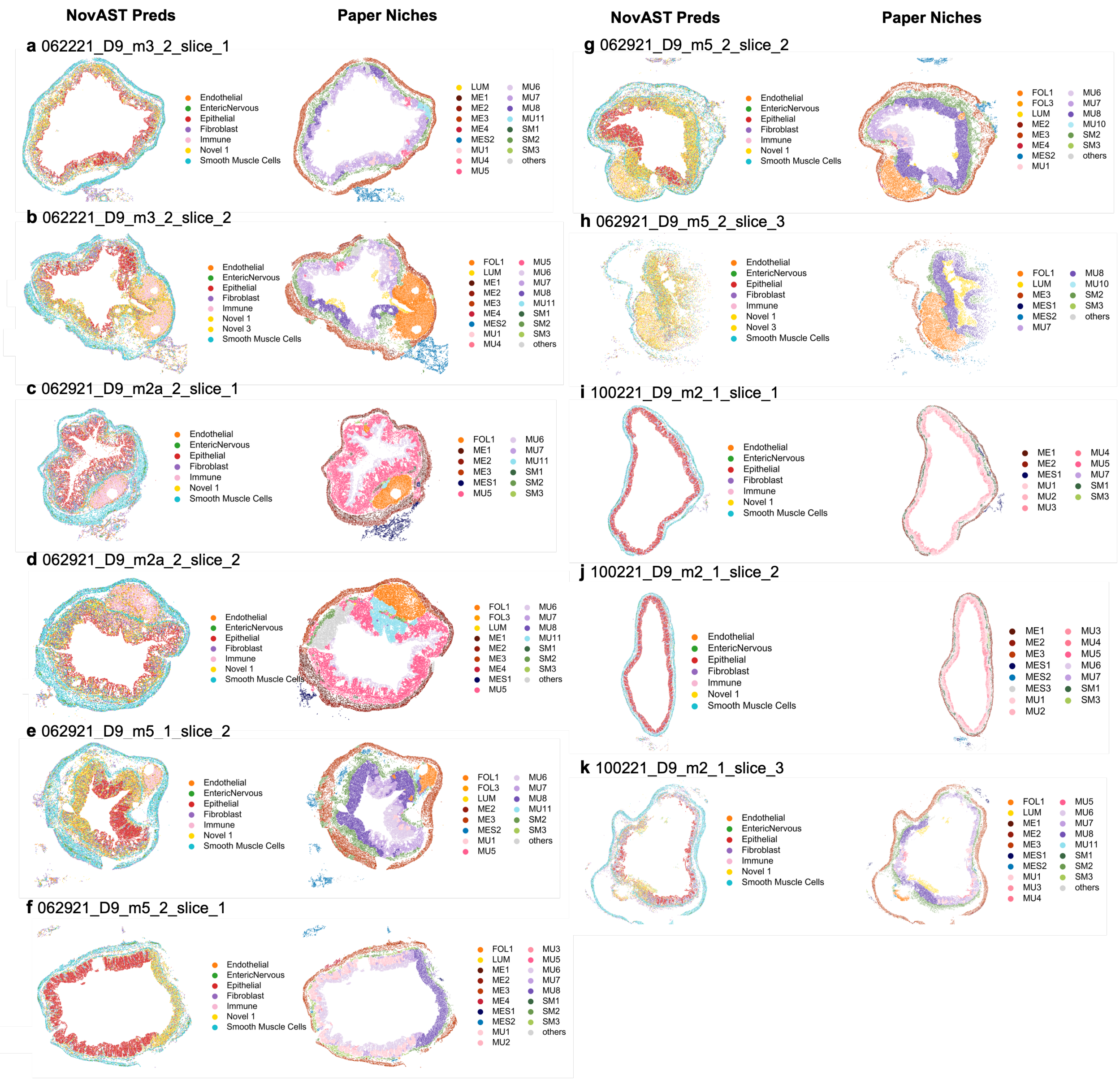

**Supplementary Figure 28**: **Spatial visualization of predicted cell types across additional tissue slices in the mouse MERFISH inflamed colon dataset (set 1)**
Spatial maps show the cell type predictions from NovAST (left column) and niche annotations from [8] (right column). Eleven slices presented here.

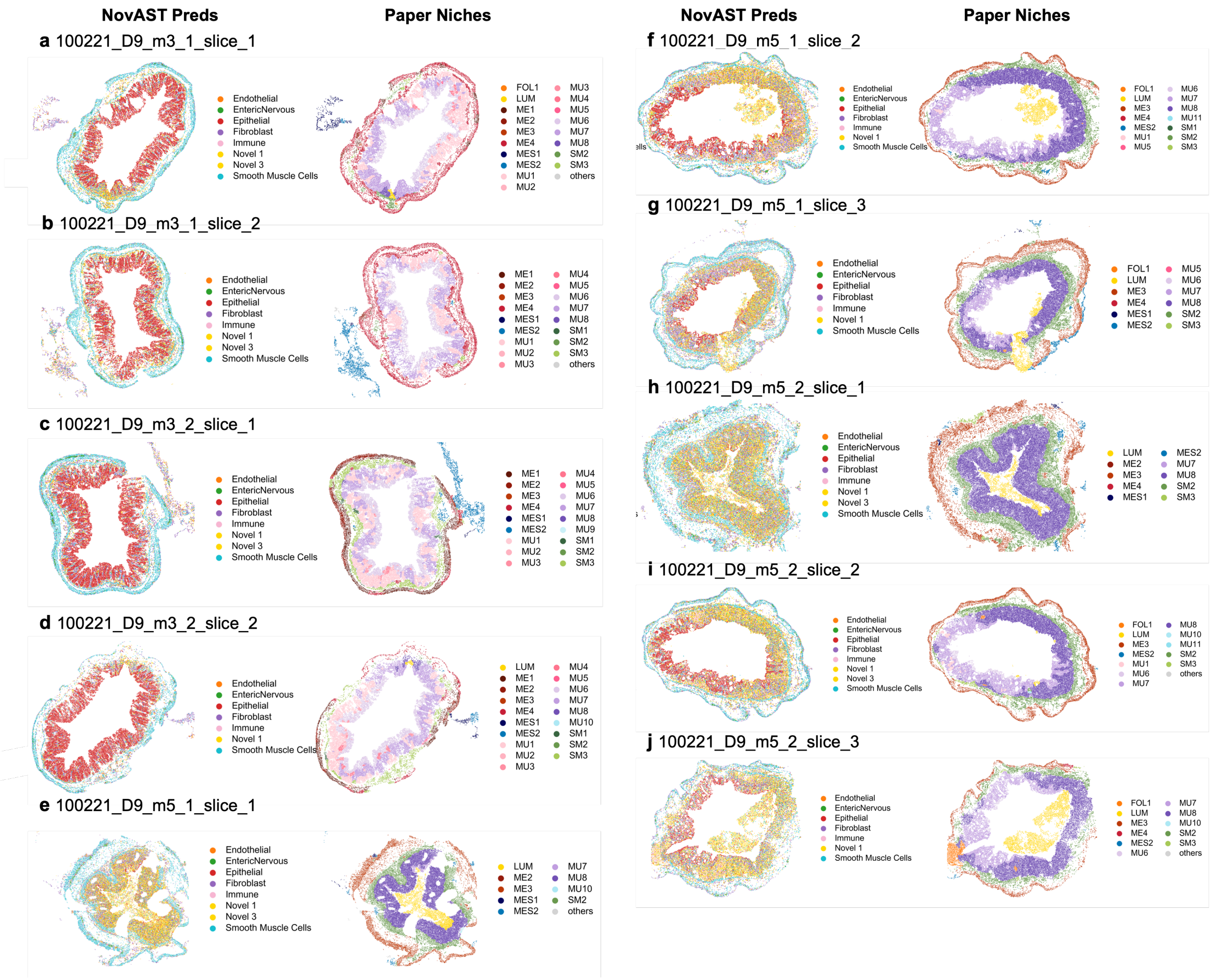

**Supplementary Figure 29**: **Spatial visualizations of additional tissue slices in the mouse MERFISH inflamed colon dataset (set 2)**

Same format as Supplementary Figure 28, showing ten additional slices.

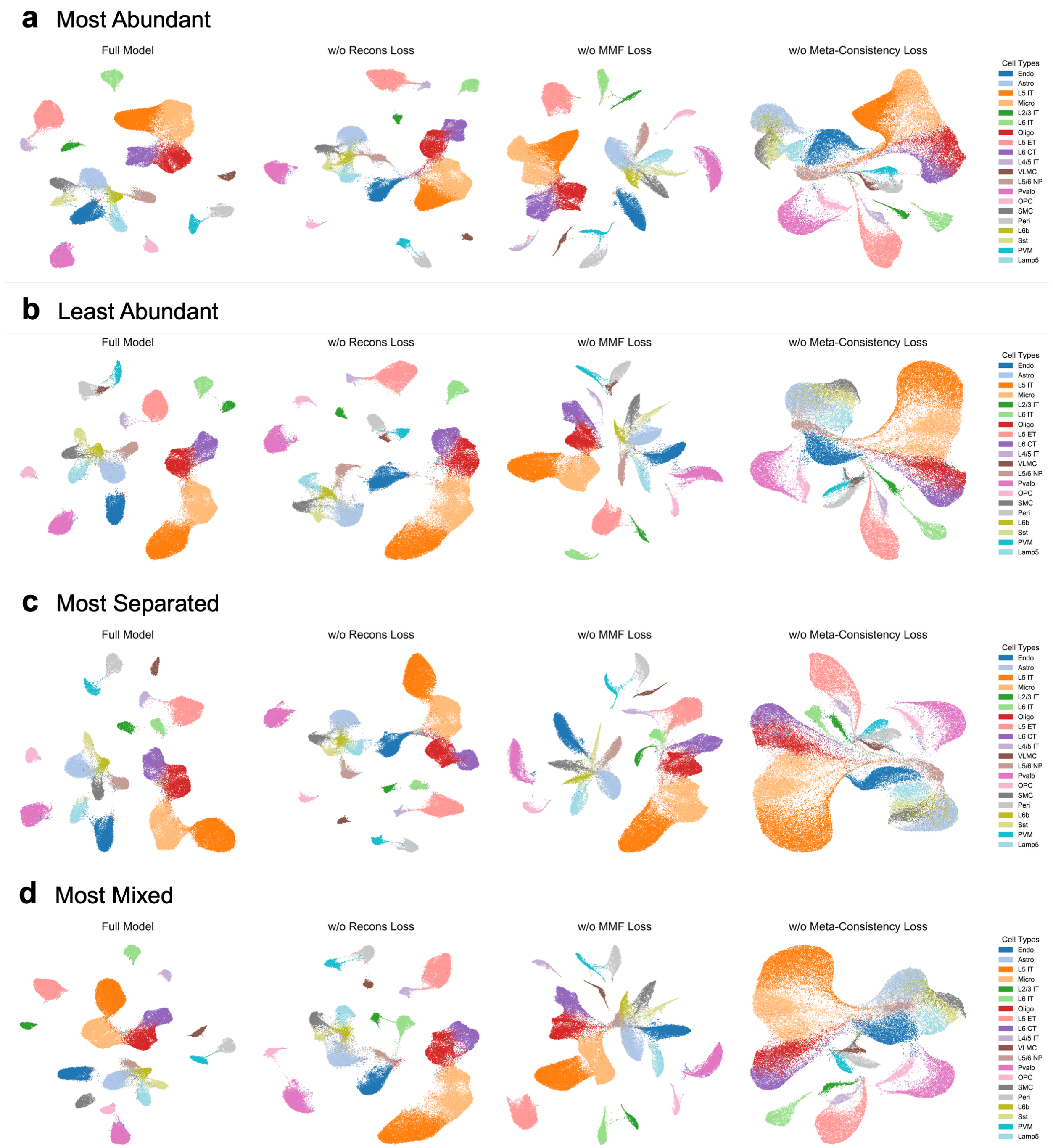

**Supplementary Figure 30**: **UMAP visualization of loss ablation in the constrained autoencoder on the mouse MERFISH MOp dataset**

Each panel summarizes results under one of four pseudo-novel scenarios: **(a)** Most Abundant, **(b)** Least Abundant, **(c)** Most Separated, and **(d)** Most Mixed. Within each row, the leftmost subpanel shows the full model incorporating all loss components, whereas the remaining three subpanels depict models with individual loss terms removed (reconstruction, min-max feature (MMF), and meta-consistency loss, respectively).

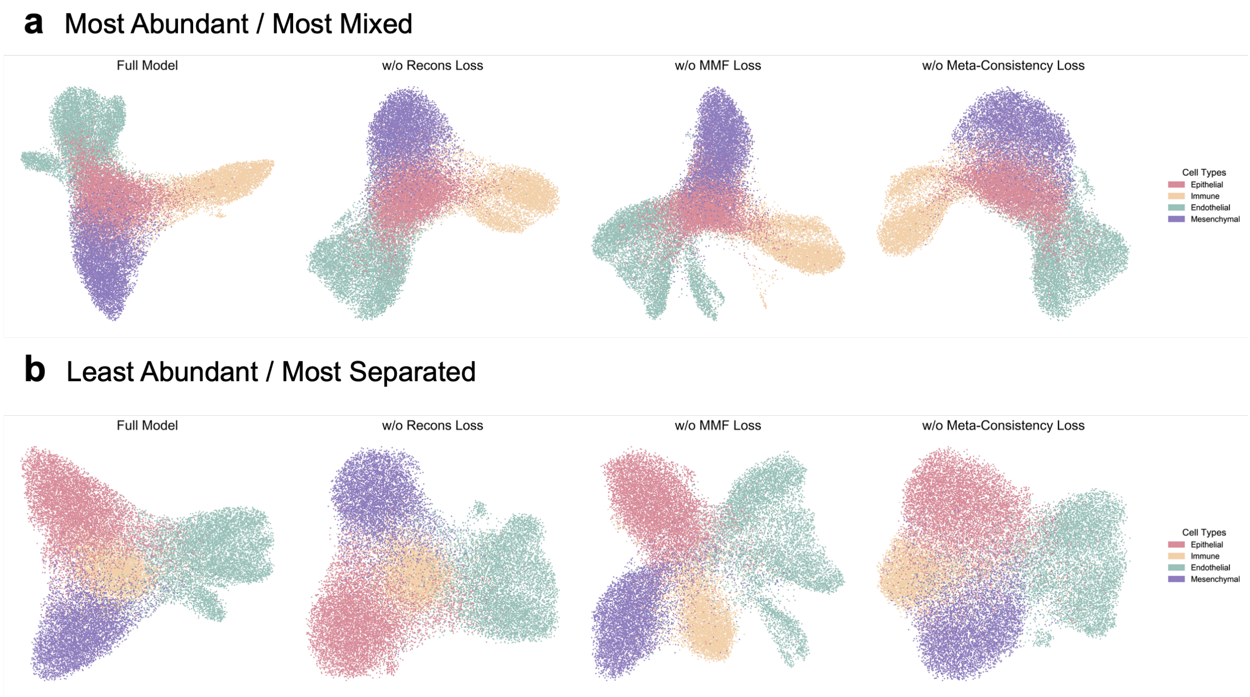

**Supplementary Figure 31**: **UMAP visualization of loss ablation in the constrained autoencoder on the human Xenium lung dataset**
Each panel summarizes results under one of four pseudo-novel scenarios: **(a)** Most Abundant / Most Mixed and **(b)** Least Abundant / Most Separated. Within each row, the leftmost subpanel shows the full model incorporating all loss components, whereas the remaining three subpanels depict models with individual loss terms removed (reconstruction, min-max feature (MMF), and meta-consistency loss, respectively).

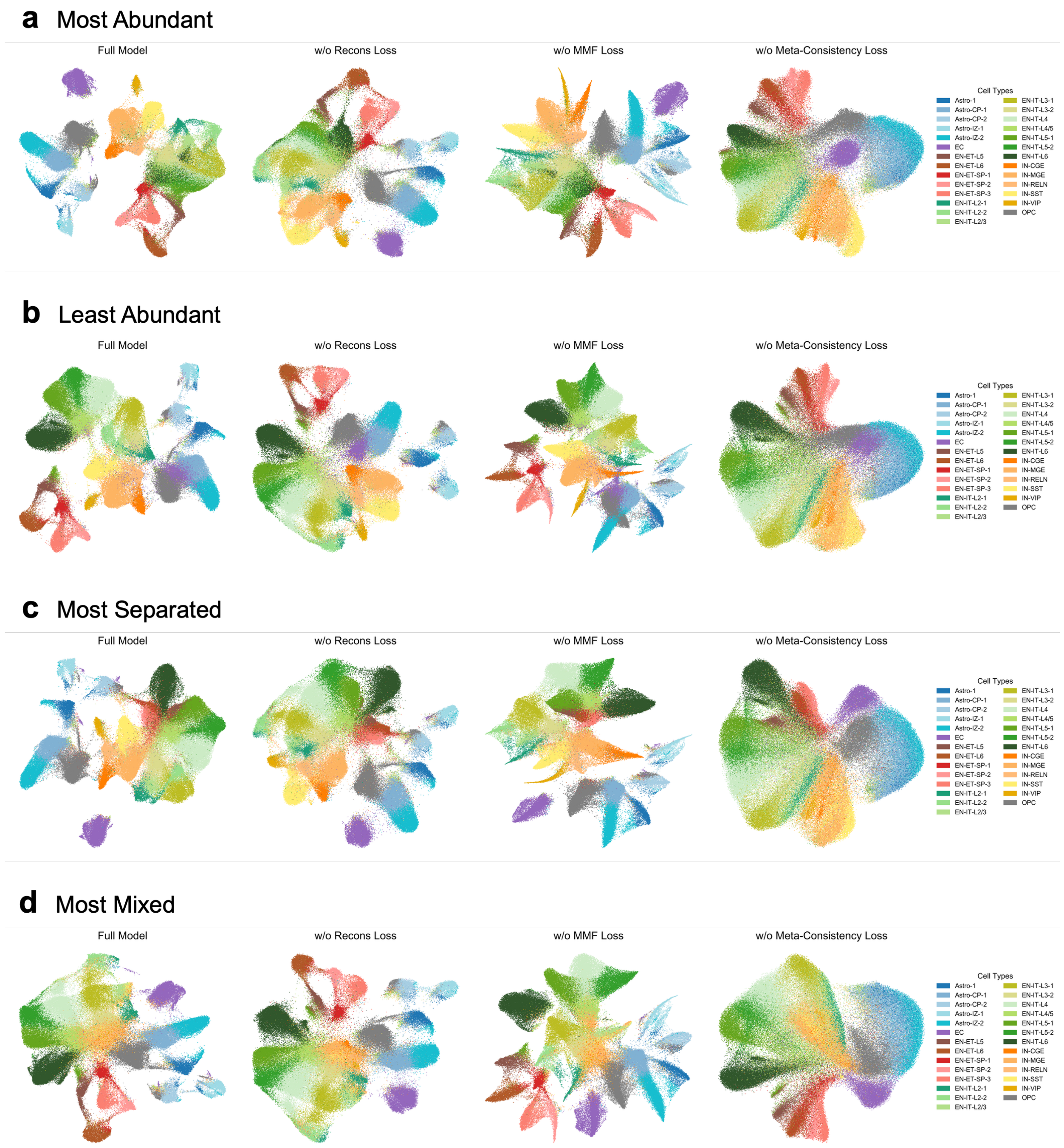

**Supplementary Figure 32**: **UMAP visualization of loss ablation in the constrained autoencoder on the human MERFISH developmental cortex dataset**
Each panel summarizes results under one of four pseudo-novel scenarios: **(a)** Most Abundant, **(b)** Least Abundant, **(c)** Most Separated, and **(d)** Most Mixed. Within each row, the leftmost subpanel shows the full model incorporating all loss components, whereas the remaining three subpanels depict models with individual loss terms removed (reconstruction, min-max feature (MMF), and meta-consistency loss, respectively).

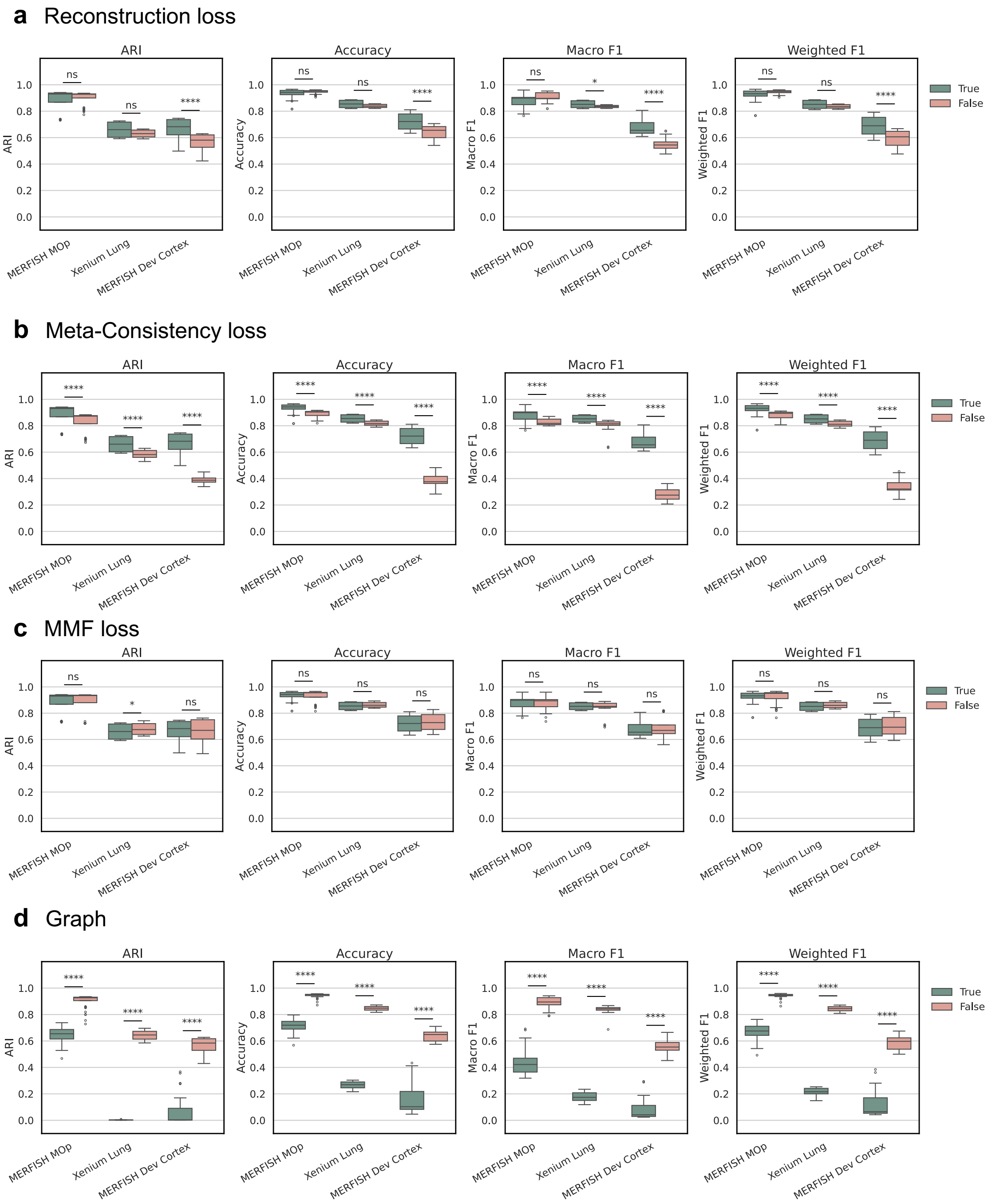

**Supplementary Figure 33**: **Ablation studies on loss terms and the impact of spatial information in the constrained autoencoder (caption continued next page)**

Panels **(a–c)** evaluate the effect of removing individual loss components: **(a)** reconstruction loss, **(b)** meta-consistency loss, and **(c)** min-max feature (MMF) loss, on NovAST performance. Panel **(d)** compares models trained with spatial information (graph = True), where graph attention layers (GAT) operate on a $k$-nearest neighbor (kNN) graph constructed from spatial coordinates, versus without spatial information (graph = False), where the autoencoder uses fully connected layers only. Each panel comprises four subpanels corresponding to evaluation metrics: ARI, Accuracy, Macro F1, and Weighted F1. Within each subpanel, three groups represent results from the mouse MERFISH MOp, human Xenium lung, and human MERFISH developmental cortex datasets under all scenarios. Each group contains paired boxplots showing the distribution of metric values with (True) and without (False) the respective loss term or spatial graph. Statistical significance was assessed using a two-sided Mann–Whitney U test across experimental seeds; significance levels are denoted by horizontal bars above boxpairs (**** indicates $P < 0.0001$, * indicates $P < 0.05$, and ns indicates not significant).

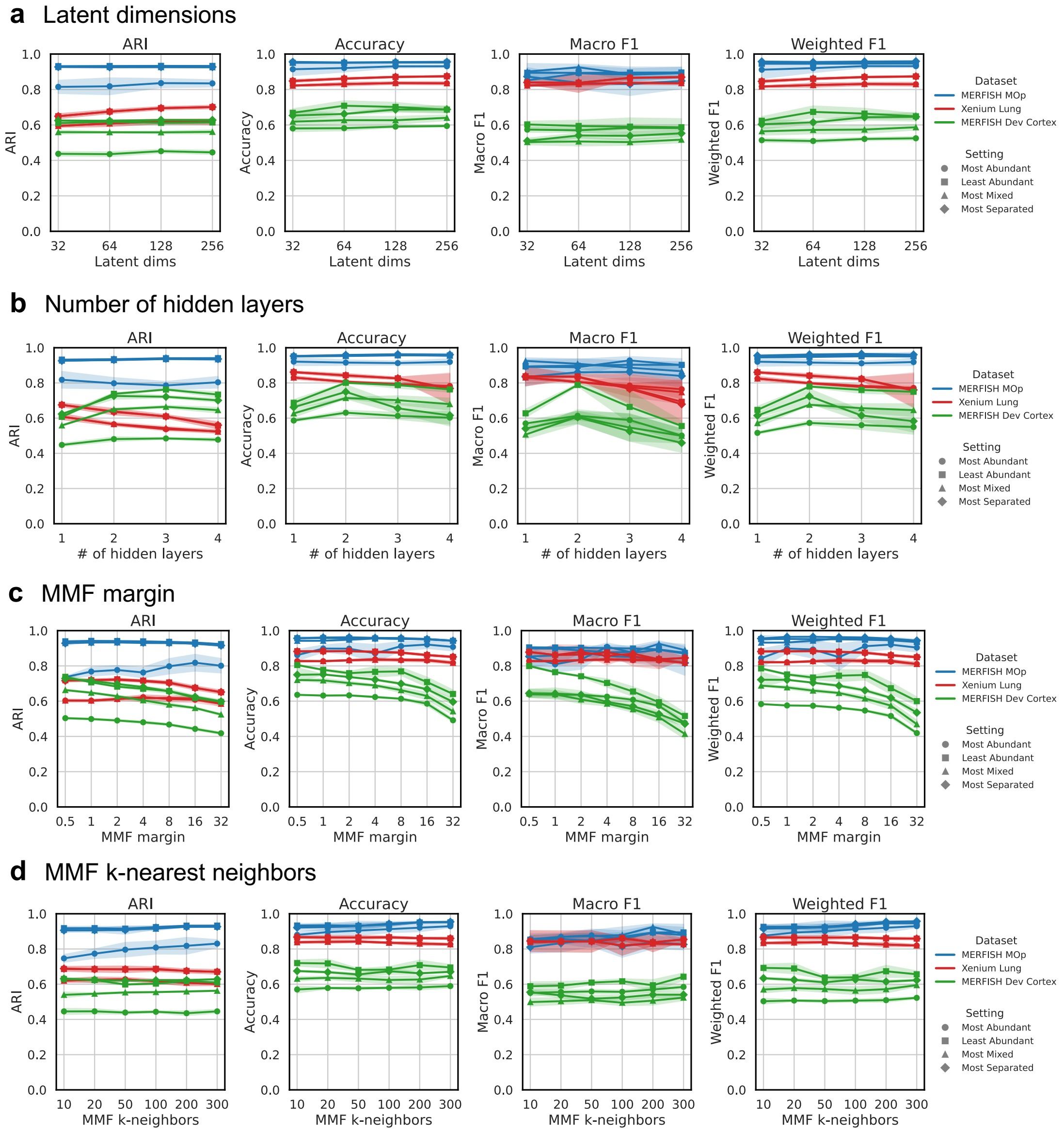

**Supplementary Figure 34**: **Parameter sensitivity analysis for latent dimension, network depth, and MMF loss settings in the constrained autoencoder**
Panels **(a–d)** evaluate the effect of key hyperparameters of constrained autoencoder on NovAST performance: **(a)** latent dimension size, **(b)** number of hidden layers, **(c)** margin parameter of the min–max feature (MMF) loss, and **(d)** number of $k$-nearest neighbors used for the MMF loss. Each panel contains four subpanels corresponding to evaluation metrics: ARI, Accuracy, Macro F1, and Weighted F1. Within each subpanel, line plots depict the mean metric values across a range of parameter choices, with shaded regions indicating standard deviations over experimental seeds. Results are shown for three datasets (mouse MERFISH MOp, human Xenium lung, and human MERFISH developmental cortex) distinguished by color. Each dataset includes results from four pseudo-novel scenarios, represented by distinct markers.

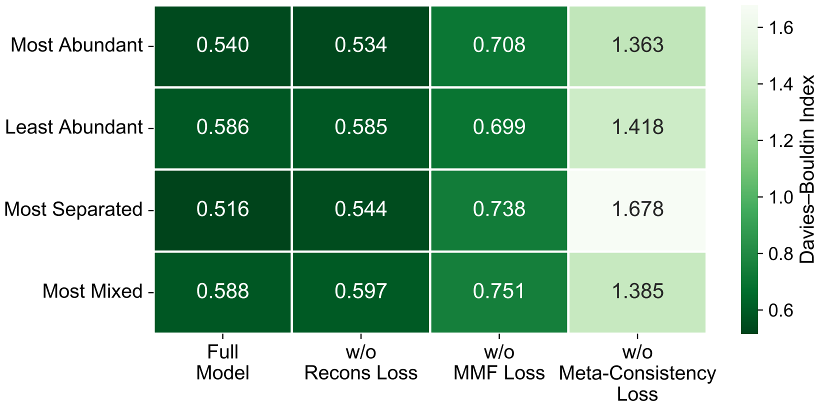

**Supplementary Figure 35**: **Ablation analysis of loss components by cluster compactness on the mouse MERFISH MOp dataset**
The Davies-Bouldin Index on mouse MERFIH MOp across each pseudo-novel setting for the full model incorporating all loss components in contrast to models with individual loss terms removed (reconstruction, min-max feature (MMF), and meta-consistency loss, respectively).

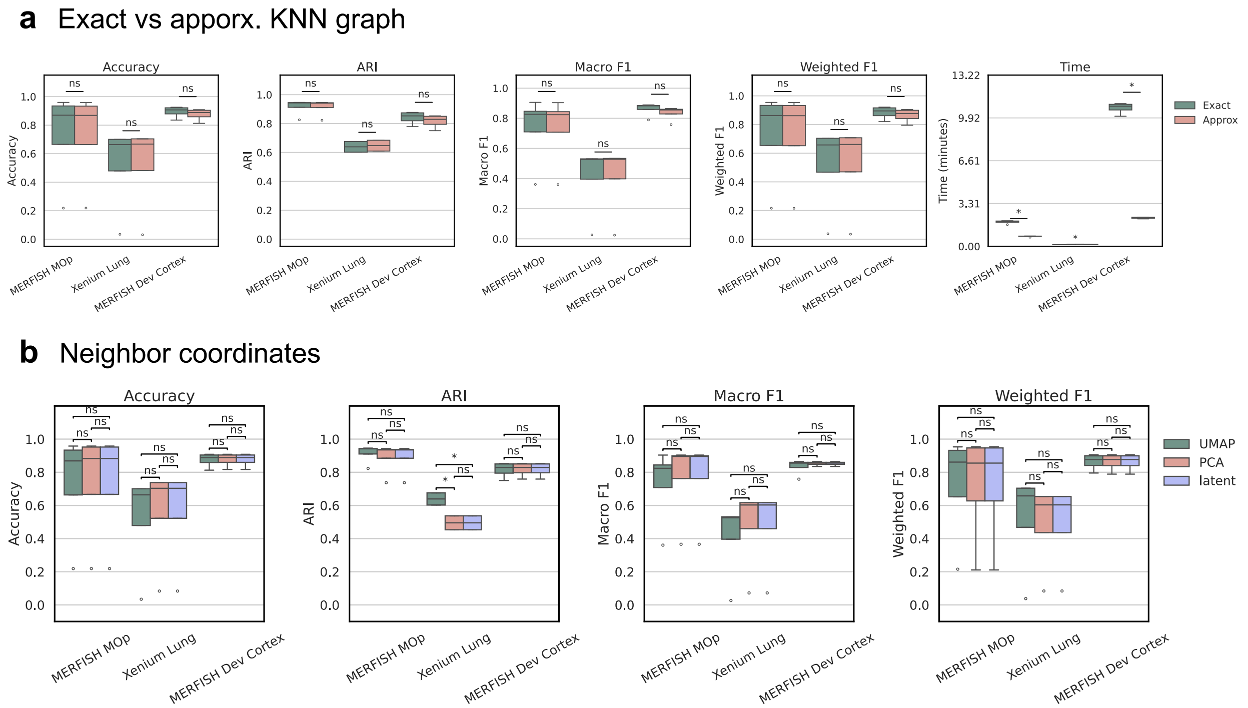

**Supplementary Figure 36**: **Sensitivity analysis of graph construction and coordinate representations**

**(a)** Comparison of exact versus approximate $k$-nearest neighbor (KNN) graphs for label spreading. Five subpanels correspond to evaluation metrics: ARI, Accuracy, Macro F1, Weighted F1, and computation time. Within each subpanel, three groups represent results from the mouse MERFISH MOp, human Xenium lung, and human MERFISH developmental cortex datasets under all scenarios. **(b)** Comparison of neighbor inference using coordinates from UMAP, PCA, or the original latent space for confidence score calculation. Four subpanels correspond to ARI, Accuracy, Macro F1, and Weighted F1. Within each subpanel, three columns represent results from the mouse MERFISH MOp, human Xenium lung, and human MERFISH developmental cortex datasets under all scenarios. Statistical significance was assessed using a two-sided Mann–Whitney U test; significance levels are indicated by horizontal bars above box pairs * indicates $P < 0.05$ and ns indicates not siginificant.

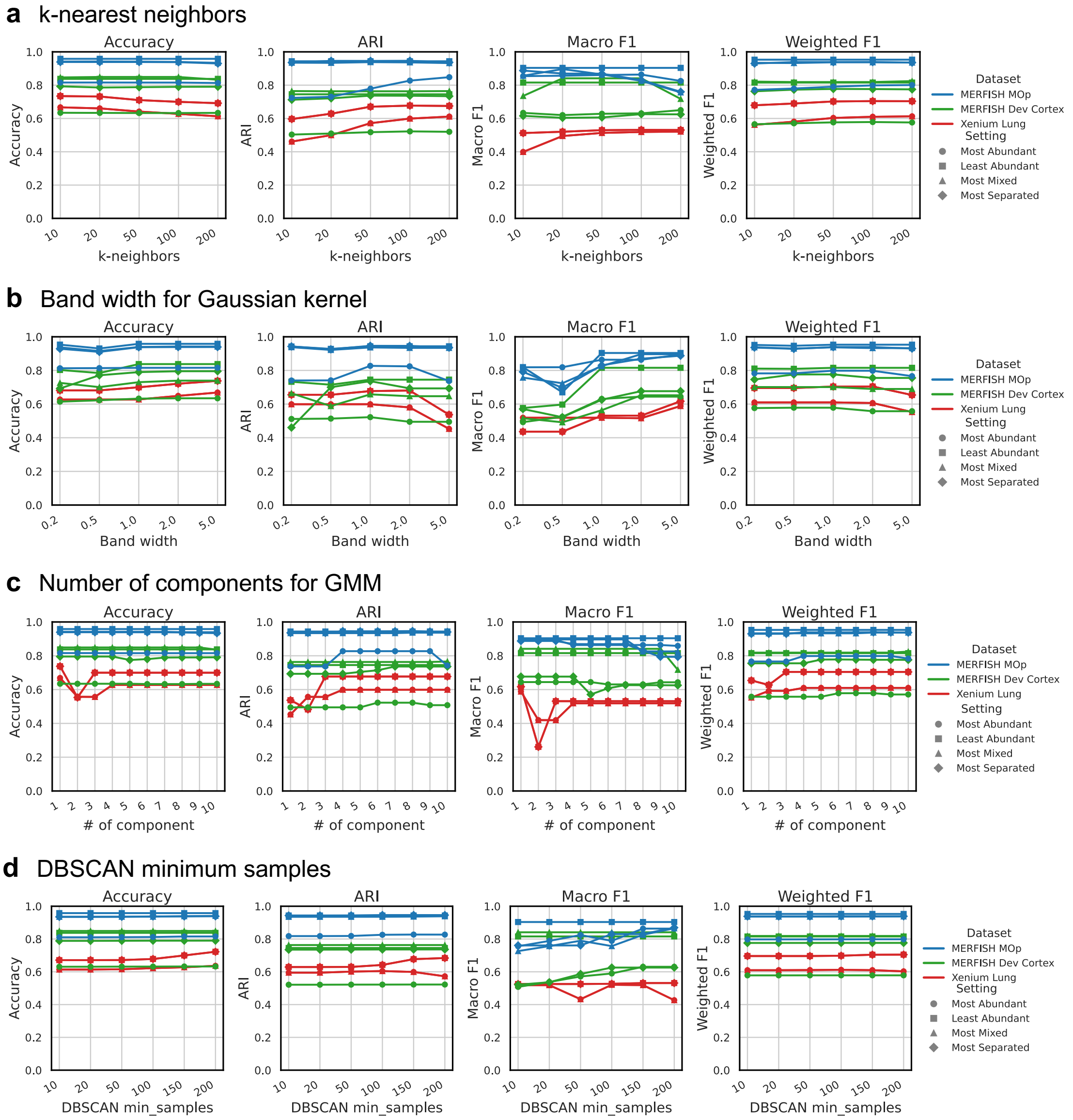

**Supplementary Figure 37**: **Parameter sensitivity analysis in novel cell type identification**

Panels **(a–d)** evaluate the effect of key hyperparameters in novel cell type identification on NovAST performance: **(a)** the number of $k$-nearest neighbors, **(b)** bandwidth of the Gaussian kernel, **(c)** number of components for Gaussian mixture modeling (GMM), and **(d)** minimum number of samples parameter for DBSCAN clustering. Each panel contains four subpanels corresponding to evaluation metrics: ARI, Accuracy, Macro F1, and Weighted F1. Within each subpanel, line plots depict metric values across a range of parameter choices. Results are shown for three datasets (mouse MERFISH MOp, human Xenium lung, and human MERFISH developmental cortex) distinguished by color. Each dataset includes results from four pseudo-novel scenarios, represented by distinct markers.

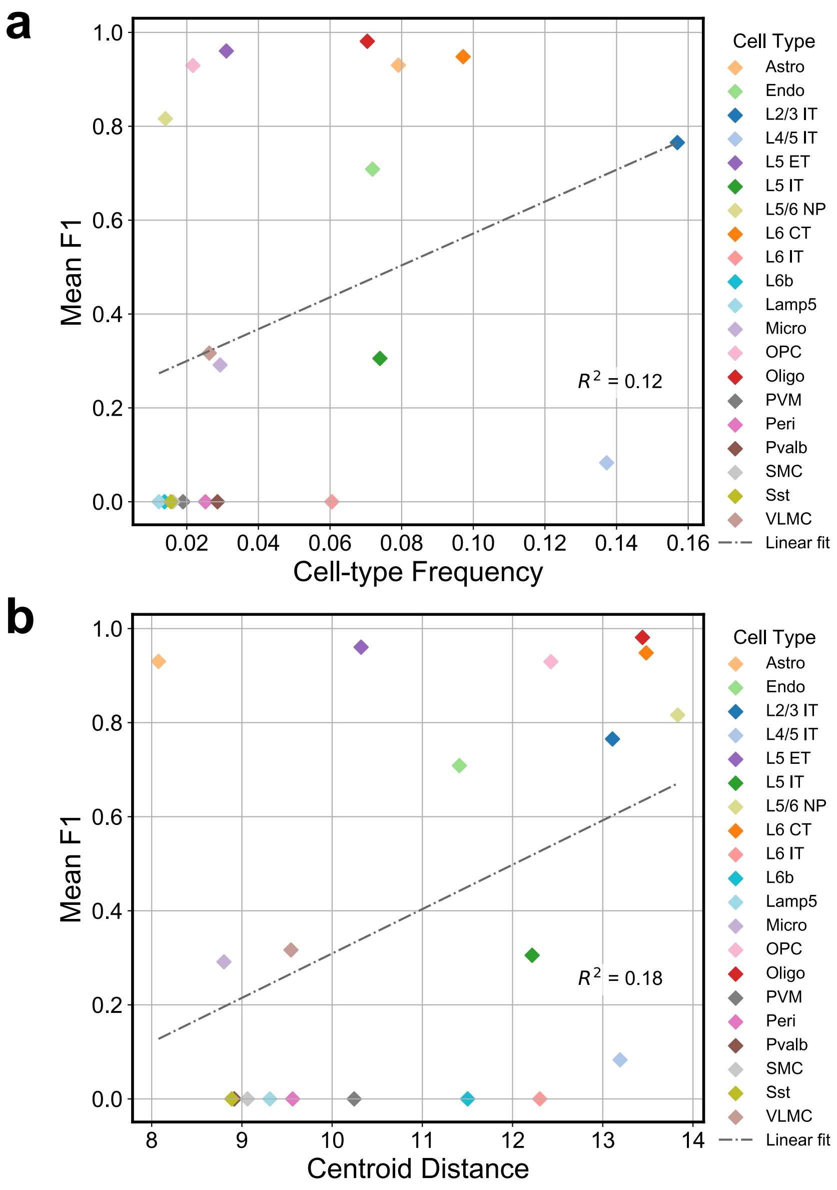

**Supplementary Figure 38**: **Impact of cell type frequency and cluster separation on novel cell type detection accuracy**

**(a)** Scatter plot showing the relationship between cell type frequency in the target dataset and the mean F1 score for novel cell type identification, averaged over 10 runs, with each cell type been removed from reference dataset in turn. The dash-dot line indicates a linear fit between cell-type frequency and mean F1, yielding $R^2 = 0.12$ between the predicted and observed F1 value. **(b)** Same as panel (a), except the x-axis represents the mean centroid distance of each cell type to all others in UMAP space.

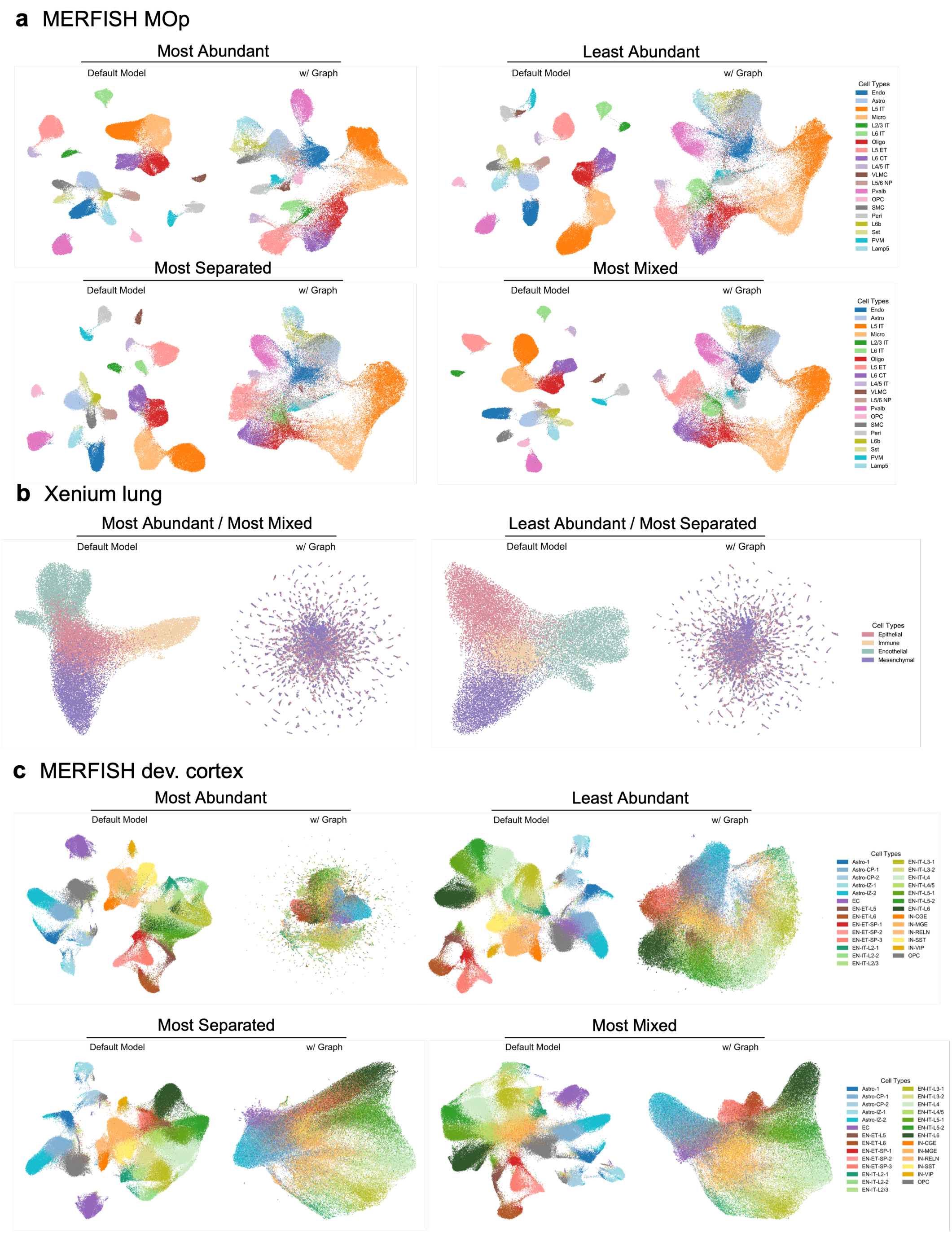

**Supplementary Figure 39**: **UMAP visualization of default and graph-based NovAST variants across datasets and pseudo-novel scenarios**

Each panel summarizes results from three datasets: **(a)** mouse MERFISH MOp, **(b)** human Xenium lung, and **(c)** human MERFISH developmental cortex. UMAPs show the embeddings from the default NovAST model and a variant incorporating graph attention networks (GATs) with graph generated from spatial coordinates. For each dataset, four pseudo-novel scenarios are shown, where one cell type at a time was withheld from the reference and treated as a pseudo-novel population in the target.

#### Supplementary Tables

##### Supplementary Table 1: Summary of datasets used in this study

Detailed information for all datasets used for benchmarking and biological exploration. Green shading denotes the primary benchmarking datasets, blue shading indicates additional benchmarking datasets, and orange shading highlights datasets used for biological analysis to evaluate cell type discrepancies arising from state shifts. For each SRT dataset, we report the dataset name, accessible links, analysis purpose, reference and target samples, number of genes, number of reference and target cells, and number of cell types.

| SRT dataset (Techniques/species/tissue) | Accessible Links | Purpose of Analysis | Reference Samples | Target Samples | No. of Genes | No. of Reference Cells | No. of Target Cells | No. of Cell Types |
|---|---|---|---|---|---|---|---|---|
| MERFISH mouse primary moter cortex (MOp) | https://doi.brainimagelibrary.org/doi/10.35077/g.21 | Quantitative | Mouse 1, Slices 21, 31, 40, 81, 102, 122, 131, 153, 200, 201, 221, 232, 241, 271, 283, 313 | Mouse 1, Slices 1, 10, 50, 62, 71, 91, 112, 162, 170, 180, 190, 212, 251, 260, 291, 301, 326 | 258 | 71,126 | 79,252 | 20 |
| Xenium human lung (control replicates from pulmonary fibrosis paper) | https://www.ncbi.nlm.nih.gov/geo/query/acc.cgi?acc=GSE250346 | Quantitative | VUHD116A | VUHD116B | 343 | 12, 372 | 29,147 | 4 |
| MERFISH human developmental cortex (replicates) | https://doi.org/10.5281/zenodo.14422018 | Quantitative | gw34_UMB5900-BA40a | gw34_UMB5900-BA40b | 300 | 191,819 | 165,520 | 7 (H1); 27 (H2) |
| CosMx SMI human non-small cell lung cancer (NSCLC) | https://zenodo.org/records/15240431 | Quantitative | section_28 | section_34 | 960 | 58,423 | 55,376 | 18 |
| MERFISH mouse cortex, aging study (24 weeks) | https://cellxgene.cziscience.com/collections/31937775-0602-4e52-a799-b6acdd2bac2e | Quantitative | 24wk_donor10 | 24wk_donor11 | 374 | 28,105 | 28,955 | 13 |
| MERFISH mouse cortex, aging study (from 24 weeks to 90 weeks) | https://cellxgene.cziscience.com/collections/31937775-0602-4e52-a799-b6acdd2bac2e | Quantitative | 24wk, all donor | 90wk, all donor | 374 | 90,301 | 164,977 | 13 |
| MERFISH tubo-ovarian high-grade serous carcinoma (HGSC) | https://singlecell.broadinstitute.org/single_cell/study/SCP2650/hgsc-spatial-cohort-discovery-dataset | Quantitative | MER_TB21361 | MER_TB23359 | 140 | 142,788 | 250,526 | 7 |
| MERFISH whole mouse brain | https://cellxgene.cziscience.com/collections/0cca8620-8dee-45d0-aef5-23f032a5cf09 | Quantitative | Mouse C57BL6J-1, Slices 44-56 | Mouse C57BL6J-2, Slices 32-46 | 1122 | 277,315 | 350,229 | 16 |
| Xenium human spinal cord (from multiple sclerosis paper) | https://doi.org/10.5281/zenodo.8037425 | Quantitative | CTRL1 | CTRL2 | 266 | 95,825 | 99,682 | 13 |
| Xenium human lung (healthy vs diseased slices from pulmonary fibrosis paper) | https://www.ncbi.nlm.nih.gov/geo/query/acc.cgi?acc=GSE250346 | Qualitative | VUHD116A & VUHD116B | VUILD107 | 343 | 41,519 | 68,413 | 4 |
| MERFISH mouse inflamation colon | https://doi.org/10.5061/dryad.rjdfn2zh3 | Qualitative | D0, all mosues all slices | D9, all mosues all slices | 940 | 239,450 | 660,977 | 8 |

**Supplementary Table 2**: **Differentially expressed genes of novel cells identified by each method in mouse MERFISH MOp dataset under Most Abundant scenario**
The top 20 differentially expressed (DE) genes identified by each method are listed. For each method, DE analysis was performed using the predicted cluster most overlapping with the ground-truth novel cells, compared against all remaining cells. The bracketed label under each method name indicates the predicted cluster used for testing. The last row lists genes uniquely identified by NovAST.

| Rank | NovAST (Novel 1) | Tangram (L4/5 IT) | SpatialID (L4/5 IT) | STELLAR (L4/5 IT) |
|---|---|---|---|---|
| 1 | Otof | Prdm8 | Unc5d | Unc5d |
| 2 | Lamp5 | Unc5d | Cux2 | Cux2 |
| 3 | Calb1 | Slc30a3 | Prdm8 | Prdm8 |
| 4 | Cux2 | Cux2 | Slc30a3 | Slc30a3 |
| 5 | Wipf3 | Prr16 | Lamp5 | Lamp5 |
| 6 | C1ql3 | Fam19a2 | Cdh12 | Cdh12 |
| 7 | Grm1 | Cdh12 | Fam19a2 | Fam19a2 |
| 8 | Pcdh8 | Slc17a6 | Prr16 | Prr16 |
| 9 | Trp53i11 | Syndig1 | C1ql3 | C1ql3 |
| 10 | Sertm1 | Lamp5 | Syndig1 | Syndig1 |
| 11 | Nos1 | Rorb | Ptprk | Ptprk |
| 12 | Meis2 | Cntnap5b | Satb2 | Sertm1 |
| 13 | Slc30a3 | Ptprk | Sertm1 | Calb1 |
| 14 | Rgs6 | Satb2 | Slc17a6 | Satb2 |
| 15 | Rfx3 | C1ql3 | Calb1 | Slc17a6 |
| 16 | Camk2d | Brinp3 | Cntnap5b | Cntnap5b |
| 17 | Trpc6 | Col25a1 | Wipf3 | Wipf3 |
| 18 | Dscaml1 | Cbln4 | Col25a1 | Col25a1 |
| 19 | Unc5d | Tcap | Flrt3 | Flrt3 |
| 20 | Ccbe1 | Cd24a | Rorb | Rorb |
| **NovAST Unique Genes**: Otof, Grm1, Pcdh8, Trp53i11, Nos1, Meis2, Rgs6, Rfx3, Camk2d, Trpc6, Dscaml1, Ccbe1 | | | | |

**Supplementary Table 3**: **Differentially expressed genes of novel cells identified by each method in human Xenium lung dataset under Most Abundant / Most Mixed scenario**

The top 20 differentially expressed (DE) genes identified by each method are listed. For each method, DE analysis was performed using the predicted cluster most overlapping with the ground-truth novel cells, compared against all remaining cells. The bracketed label under each method name indicates the predicted cluster used for testing. The last row lists genes uniquely identified by NovAST.

| Rank | NovAST (Novel 1) | Tangram (Immune) | SpatialID (Immune) | STELLAR (Immune) |
|---|---|---|---|---|
| 1 | EPAS1 | PTPRC | PTPRC | PECAM1 |
| 2 | PECAM1 | FCER1G | FCER1G | PTPRC |
| 3 | SPARCL1 | LYZ | LYZ | EPAS1 |
| 4 | FCN3 | PECAM1 | S100A8 | FCER1G |
| 5 | BMPR2 | VIM | S100A9 | VIM |
| 6 | IL7R | S100A8 | VIM | LYZ |
| 7 | KDR | EPAS1 | PECAM1 | S100A8 |
| 8 | GNG11 | S100A9 | FCGR3A | IL7R |
| 9 | CLDN5 | FCGR3A | MS4A7 | S100A9 |
| 10 | HES1 | MS4A7 | ITGAX | TGFB1 |
| 11 | BCL2L1 | ITGAX | TGFB1 | FCGR3A |
| 12 | ZEB1 | TGFB1 | CD68 | FCN3 |
| 13 | CA4 | IL7R | S100A12 | ITGAX |
| 14 | POSTN | CD68 | EPAS1 | MS4A7 |
| 15 | ITGAV | S100A12 | AIF1 | BMPR2 |
| 16 | HERPUD1 | ITGAM | ITGAM | S100A12 |
| 17 | CD34 | AIF1 | IL7R | SLC25A37 |
| 18 | RAMP2 | SLC25A37 | SLC25A37 | UBE2J1 |
| 19 | CD274 | MCEMP1 | MCEMP1 | CD68 |
| 20 | IL4R | UBE2J1 | MRC1 | GNG11 |
| **NovAST Unique Genes**: SPARCL1, KDR, CLDN5, HES1, BCL2L1, ZEB1, CA4, POSTN, ITGAV, HERPUD1, CD34, RAMP2, CD274, IL4R | | | | |

**Supplementary Table 4**: **Differentially expressed genes of novel cells identified by each method in MERFISH developmental cortex dataset under Most Separated scenario**

The top 20 differentially expressed (DE) genes identified by each method are listed. For each method, DE analysis was performed using the predicted cluster most overlapping with the ground-truth novel cells, compared against all remaining cells. The bracketed label under each method name indicates the predicted cluster used for testing. The last row lists genes uniquely identified by NovAST.

| Rank | NovAST (Novel 2) | Tangram (EN-IT-L5-1) | SpatialID (EN-IT-L5-1) | STELLAR (EN-IT-L5-1) |
|---|---|---|---|---|
| 1 | FEZF2 | FEZF2 | FEZF2 | FEZF2 |
| 2 | CRYM | NEFM | NEFM | NEFM |
| 3 | ETV1 | CRYM | CRYM | TAFA1 |
| 4 | PEX5L | TAFA1 | PEX5L | CRYM |
| 5 | GLRA2 | PEX5L | ETV1 | PEX5L |
| 6 | TLE4 | GLRA2 | HS3ST4 | ETV1 |
| 7 | BCL11B | NEUROD6 | GLRA2 | GLRA2 |
| 8 | HS3ST4 | ETV1 | CLSTN2 | VSNL1 |
| 9 | TRPM3 | CLSTN2 | KLHL1 | SUSD4 |
| 10 | TRMT9B | VSNL1 | SUSD4 | NEUROD6 |
| 11 | NEFM | SUSD4 | ATP2B4 | CLSTN2 |
| 12 | PCDH17 | KLHL1 | NEUROD6 | KLHL1 |
| 13 | HSPA12A | TMEM163 | VSNL1 | TMEM163 |
| 14 | GABRB2 | KIAA0319 | VAT1L | KIAA0319 |
| 15 | SUSD4 | OPCML | KIAA0319 | RGS6 |
| 16 | TSHZ3 | ATP2B4 | RAP1GAP | PCDH17 |
| 17 | CLSTN2 | HS3ST4 | SORCS1 | FSTL5 |
| 18 | SORCS1 | VAT1L | TAFA1 | SPOCK1 |
| 19 | VAT1L | FOXP1 | OPCML | ATP2B4 |
| 20 | FOXP2 | RGS6 | NEUROD2 | NRN1 |
| **NovAST Unique Genes**: TLE4, BCL11B, TRPM3, TRMT9B, HSPA12A, GABRB2, TSHZ3, FOXP2 | | | | |

**Supplementary Table 5**: **Mann–Whitney U-test of Novel 1 cells vs. non-novel cells across histopathology-annotated regions in the Xenium IPF dataset**

The table reports, for every histopathology-annotated region, the U statistic, p-value, mean distance for Novel 1 and non-novel cells, and the corresponding group sizes. All tests are one-sided, evaluating whether Novel 1 cells lie closer to the region than non-novel cells. The orange shading indicates regions that are statistically significantly closer to the Novel 1 cells than to the non-novel cells ($p < 0.05$), whereas the blue shading denotes regions that are not statistically significant.

| Region | U statistic | p value | Mean Dist. Novel | Mean Dist. Non-novel | # of Novel | # of Non-novel |
|---|---|---|---|---|---|---|
| Muscularized Artery | 191822403 | 6.42E-62 | 4305.217368 | 4850.22122 | 7110 | 61303 |
| Large Airway | 202114672 | 5.39E-24 | 5201.621638 | 5617.92044 | 7110 | 61303 |
| Goblet Cell Metaplasia | 202756653 | 3.08E-22 | 7866.612821 | 8361.15651 | 7110 | 61303 |
| Fibrosis | 203530900 | 2.36E-20 | 2071.224626 | 2289.04294 | 7110 | 61303 |
| Severe Fibrosis | 209904527 | 1.77E-07 | 3727.249835 | 3899.77664 | 7110 | 61303 |
| Remodeled Epithelium | 210799568 | 3.02E-06 | 3811.922725 | 4079.90594 | 7110 | 61303 |
| Hyperplastic AECs | 212659775 | 0.00041194 | 3856.502109 | 3980.23185 | 7110 | 61303 |
| Mixed Inflammation | 221121571 | 0.97865874 | 1474.081385 | 1470.52681 | 7110 | 61303 |
| Fibroblastic Focus | 226275865 | 0.99999994 | 7484.749389 | 7229.99462 | 7110 | 61303 |
| Epithelial Detachment | 227696289 | 1 | 6352.653792 | 6118.25748 | 7110 | 61303 |

**Supplementary Table 6**: **Mann–Whitney U-test of Novel 2 cells vs. non-novel cells across histopathology-annotated regions in the Xenium IPF dataset.**
The table reports, for every region, the U statistic, $p$-value, mean distance for Novel 2 and non-novel cells, and the corresponding group sizes. All tests are one-sided, evaluating whether Novel 2 cells lie closer to the region than non-novel cells. The orange shading indicates regions that are statistically significantly closer to the Novel 2 cells than to the non-novel cells ($p$ < 0.05), whereas the blue shading denotes regions that are not statistically significant.

| Region | U statistic | p value | Mean Dist. Novel | Mean Dist. Non-novel | # of Novel | # of Non-novel |
|---|---|---|---|---|---|---|
| Mixed Inflammation | 35724519 | 6.18E-15 | 1187.67875 | 1476.03851 | 1220 | 67193 |
| Epithelial Detachment | 36540763 | 3.89E-11 | 5480.77303 | 6154.63462 | 1220 | 67193 |
| Severe Fibrosis | 36853050.5 | 7.33E-10 | 3452.99225 | 3889.6329 | 1220 | 67193 |
| Fibroblastic Focus | 37338385 | 4.69E-08 | 6680.088 | 7266.93585 | 1220 | 67193 |
| Hyperplastic AECs | 37746078 | 1.06E-06 | 3633.47518 | 3973.43538 | 1220 | 67193 |
| Remodeled Epithelium | 43106562 | 0.99903043 | 4203.17327 | 4049.31128 | 1220 | 67193 |
| Large Airway | 44393073 | 0.99999968 | 6038.48587 | 5566.23389 | 1220 | 67193 |
| Muscularized Artery | 44572133 | 0.99999992 | 5235.60585 | 4785.55456 | 1220 | 67193 |
| Goblet Cell Metaplasia | 45005871.5 | 1 | 8997.7041 | 8297.26899 | 1220 | 67193 |
| Fibrosis | 46181888 | 1 | 2738.65728 | 2257.83109 | 1220 | 67193 |

### References

[1] Meng Zhang, Stephen W. Eichhorn, Brian Zingg, Zizhen Yao, Kaelan Cotter, Hongkui Zeng, Hongwei Dong, and Xiaowei Zhuang. Spatially resolved cell atlas of the mouse primary motor cortex by merfish. *Nature*, 598(7879):137–143, 2021.

[2] Yu A Malkov and Dmitry A Yashunin. Efficient and robust approximate nearest neighbor search using hierarchical navigable small world graphs. *IEEE transactions on pattern analysis and machine intelligence*, 42(4):824–836, 2018.

[3] Lawrence Hubert and Phipps Arabie. Comparing partitions. *Journal of classification*, 2(1):193–218, 1985.

[4] Juri Opitz and Sebastian Burst. Macro f1 and macro f1. *arXiv preprint arXiv:1911.03347*, 2019.

[5] Jacob Cohen. A coefficient of agreement for nominal scales. *Educational and psychological measurement*, 20(1):37–46, 1960.

[6] Brian W Matthews. Comparison of the predicted and observed secondary structure of t4 phage lysozyme. *Biochimica et Biophysica Acta (BBA)-Protein Structure*, 405(2):442–451, 1975.

[7] David L. Davies and Donald W. Bouldin. A cluster separation measure. *IEEE Transactions on Pattern Analysis and Machine Intelligence*, PAMI-1(2):224–227, 1979.

[8] Paolo Cadinu, Kisha N. Sivanathan, Aditya Misra, Rosalind J. Xu, Davide Mangani, Evan Yang, Joseph M. Rone, Katherine Tooley, Yoon-Chul Kye, Lloyd Bod, Ludwig Geistlinger, Tyrone Lee, Randall T. Mertens, Noriaki Ono, Gang Wang, Liliana Sanmarco, Francisco J. Quintana, Ana C. Anderson, Vijay K. Kuchroo, Jeffrey R. Moffitt, and Roni Nowarski. Charting the cellular biogeography in colitis reveals fibroblast trajectories and coordinated spatial remodeling. *Cell*, 187(7):2010–2028, April 2024.
